# Automated Model Refinement Using Perturbation-Observation Pairs

**DOI:** 10.1101/2023.11.14.567002

**Authors:** Kyu Hyong Park, Jordan C. Rozum, Réka Albert

**Author notes:** Corresponding authors: Kyu Hyong Park, Réka Albert.

## Abstract

Network-based dynamic modeling is useful for studying the responses of complex biomolecular systems to environmental changes and internal perturbations. In modeling signal transduction and other regulatory networks, it is common to integrate evidence from perturbation (e.g. gene knockout) - observation pairs, where the perturbed and observed variables may be distant in the network. For a model to capture these non-local effects, its construction, validation, and refinement necessarily involve trial and error, constrained by domain knowledge.

We propose and implement a genetic algorithm-based workflow to streamline model refinement. This workflow applies to any biological system for which an interaction network and enough perturbation experiments exist. We implement our workflow for Boolean networks, which are a popular and successful tool for modeling biological systems. The algorithm we introduce adjusts the functions of the model to enhance agreement with a corpus of curated experimental results and leverages existing mechanistic knowledge to automatically limit the search space to biologically plausible models. To account for the interdependence of experimental results, we develop a hierarchical scoring technique for assessing model performance. Our implementation is available as the open-source Python library *boolmore*.

We demonstrate *boolmore*’s effectiveness in a published plant signaling model that exemplifies the challenges of manual model construction and refinement. This model describes how plant stomata close in response to the drought hormone abscisic acid. After several hours of automatic refinement on a personal computer, the fittest models recapture and surpass the accuracy gain achieved over two years of manual revision. The refined models yield new, testable predictions, such as explanations for the role of reactive oxygen species in drought response. By automating the laborious task of model validation and refinement, this workflow is a step towards fast, fully automated, and reliable model construction.

## Introduction

Network-based dynamic modeling is an effective avenue toward understanding the response of biological systems to changes in their environment. The system is abstracted into a network, whose nodes represent the components of the system (e.g., proteins, cells, neurons, or species) and whose edges represent the directed, causal interactions among them. The dynamic model assigns each node a state variable and a regulatory function that determines the future state of the node given the current states of its regulators.

Boolean models are the simplest discrete dynamic models. They are used to model a variety of biological systems; examples include gene regulatory networks [1], neuronal networks [2,3], ecological and social communities [4,5]. In Boolean models, the node state variables can take two values: 0, interpreted as low concentration or activity, or 1, interpreted as high concentration or activity. Boolean models are especially suitable for biomolecular networks due to the abundance of nonlinear, sigmoidal regulation in these networks [6,7], and because of these models’ ability to describe perturbation (e.g., gene knockout) experiments. Through integrating the knowledge of the biology community, Boolean models successfully capture key behaviors in the biomolecular system of interest, and make useful predictions such as identifying master regulators or drug targets [8,9] (see Text S1 for examples). Predictions derived from Boolean models were verified experimentally in a variety of biological systems [10,11]. Multiple methodologies and tools can determine the possible long-term behaviors of Boolean models [12–16]. Here we take advantage of minimal trap spaces to describe long-term behaviors. A minimal trap space, also called a quasiattractor, is a minimal set of node states that the system can be “trapped” in and that can be characterized by fixing the values of some subset of the node state variables (see Text S1 for a formal definition).

Due to recent advances in the analysis of Boolean dynamics [12–14,17], the bottleneck in analyzing a biomolecular system is increasingly the time and effort involved in building the model. When full state data (i.e., the state of all the relevant components at a given time) is available, several viable automated Boolean network inference methods are available [18–23]. For example, transcriptome data from multiple cell types can be used to infer gene regulatory network models for cell differentiation processes [20,24,25]. However, high-throughput assays (which can probe the state of many components at the same time) are not the norm in cell signaling systems, which involve difficult-to-track post-translational modifications of proteins.

Traditional experiments, still frequently used in functional biology, measure a single component in two contexts, for example, in the presence or absence of a stimulus, or in the presence or absence of a perturbation of a different component. Compilations of such experimental observations are not equivalent to high-throughput measurements because certain components are more studied than others [26] and because of inconsistencies between reported results [27]. The existing Boolean network inference methods are not suitable for such piece-wise, incomplete, and uneven data. In such systems model construction is done via manual integration of distinct pieces of experimental evidence (see Text S1 for more details of the modeling process). Although some parts of the model can be directly constrained by preexisting experiments, usually many degrees of freedom remain. Modelers often use a process of trial and error informed by the insights of domain experts. Keeping the model up to date involves the same trial and error iteration. An illustrative example of the time and effort needed for model construction, validation, and update is the Boolean model of abscisic acid-induced (ABA-induced) stomatal closure in the model plant *Arabidopsis thaliana*, which was introduced in 2006 [28], significantly updated in 2017 [29] and refined in 2018-20 [30,31].

Our aim is to speed up and automate the trial and error process needed for model validation and refinement. Specifically, we consider the problem of refining and updating an existing baseline model to better agree with existing perturbation-observation data and also incorporate new data. We assume that the baseline model has a relatively complete interaction network, missing perhaps only a few edges. We develop a genetic algorithm-based workflow to adjust the update functions of the model in a manner that optimizes the model’s agreement with curated perturbation-observation results. The workflow, and its implementation in the tool *boolmore* (BOOLean MOdel REfiner), includes multiple ways to incorporate biological expertise that can limit the genetic algorithm’s search space to models that agree with biological knowledge. We demonstrate the effectiveness of our workflow by generating refined models of ABA-induced stomatal closure that agree significantly better with a compendium of published experimental results than the previous models.

### Methods: outline of the *boolmore* tool

Our genetic algorithm-based workflow systematically tackles the huge number of modeling choices that must be considered during model validation and refinement. Genetic algorithms are a type of heuristic optimization that produce candidates through stochastic mutation and retain or eliminate them depending on a fitness score [32]. In the context of the problem considered here, the sought-after solution is the optimal refinement of an existing Boolean model, which consists of a signed interaction graph and the Boolean functions of each node. Our proposed model refinement tool, called *boolmore*, uses the starting model to build a large number of mutated offspring models with different regulatory functions. These offspring models stay consistent with modeler-specified biological constraints and with the interaction graph, unless the addition of interactions is allowed. *Boolmore* scores the fitness of each model by comparing the model’s predictions to a compendium of experimental perturbation - observation pairs. *Boolmore* is meant to be used in conjunction with domain expertise to interpret the refined model and extract new predictions from it.

The previous works most relevant to our method involve genetic-algorithm-based inference of a Boolean model based on a directed network that integrates prior knowledge of interactions and regulatory relationships [19–23,33,34]. These algorithms take as input information a compendium of steady state values of all the nodes in the unperturbed system, complemented by steady state values obtained for perturbations. Another type of relevant prior work develops answer-set-programming methods to infer a Boolean model [25] using reachability relationships between initial and final states (as available for cell differentiation) or to revise an existing Boolean model to better align with steady state or time course measurements [35,36]. We summarize in Text S2 the goals and use cases of relevant previous algorithms. We provide a comparison of *boolmore* with the algorithms BoNesis [25] and Gitsbe [34] in the Results.

*Boolmore* takes as input a compilation of experimental results, each of which describe the observed state of a node (biomolecule) in a certain context (e.g., in the presence of a signal, or in case of a perturbation of another component). It also incorporates known biological mechanisms as constraints that have to be satisfied by the regulatory functions. The model refinement process of *boolmore* repeatedly iterates through the steps depicted in Figure 1. Here we provide a brief overview; see Text S3 for the details. First, *boolmore* mutates the functions of the starting model to form new models, using a novel representation of monotonic Boolean functions. This representation has the advantage that it can easily be converted to a truth table or Blake canonical form. The mutation is constrained to preserve all the knowledge in which there is high confidence, for example the sign of each edge, which represents positive or negative regulatory influence. Every edge can be deleted and later recovered. Edges not in the original network can be added from a user-provided pool. *Boolmore* also generates crossover models in which each node’s regulatory function is randomly chosen from one of two models.

**Figure 1.**
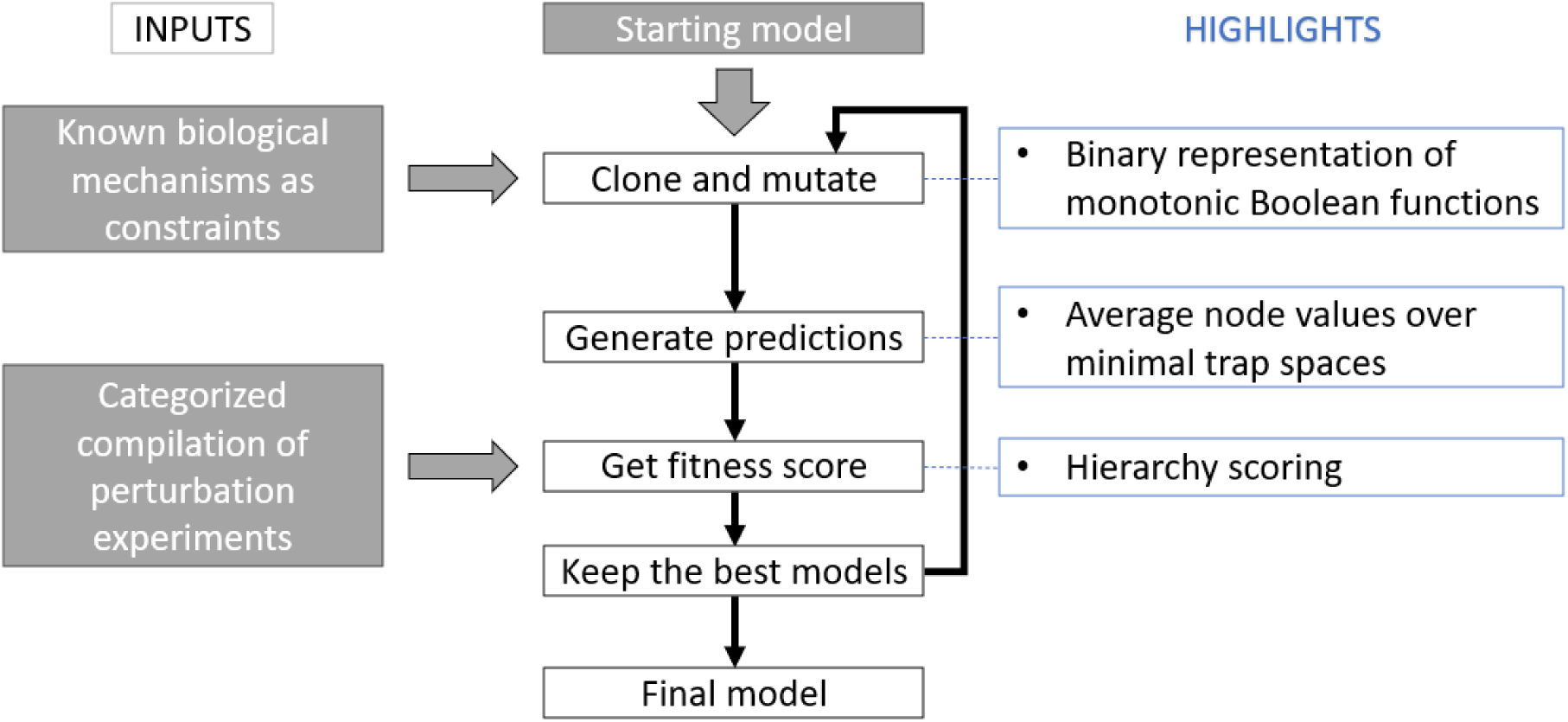
The outline of the *boolmore* tool. The starting model, known biological mechanisms and a categorized compilation of perturbation experiments are taken as input. These inputs are integrated directly into the model refinement process, as they are used to generate and score every single model. We introduced a new representation to allow the mutation of Boolean functions while maintaining consistency with the interaction graph and the biological constraints. *Boolmore* generates model predictions using minimal trap spaces and employs hierarchy scoring to ensure that biologically realistic models receive higher scores.

Second, *boolmore* generates the predictions of the models by calculating the minimal trap spaces of the model under each setting, and identifying nodes that are ON, OFF, or oscillate. Third, *boolmore* computes the model’s fitness score by quantifying the agreement of the model predictions with the experimental results. The scoring incorporates two new features. The results of each experiment are classified into five categories, including OFF, ON, and three intermediate categories. The agreement of the predictions with these categories are calculated using the curves in Table S1. The “Some” intermediate category is best satisfied by oscillating nodes or nodes that have different values in different minimal trap spaces (see Text S4 for the interpretation of this category). Furthermore, to improve the alignment of biological relevance and scoring, the experiments are grouped hierarchically and the scoring of an experiment takes into account the experiments at previous levels of the hierarchy. For example, for a model to get a score on the result of a double intervention experiment, it must also agree with the constituent single interventions. Finally, *boolmore* keeps the models with the top fitness scores, while also preferring models with fewer added edges. *Boolmore* contains multiple tunable parameters including the mutation probability and the number of models generated in each step, which the users can freely customize. We present a parameter analysis and explain our choices in Text S5.

## Results

As is common practice in network inference, we first demonstrate the performance of *boolmore* through *in-silico* benchmark studies. We used 40 published Boolean models from the Cell collective repository [37] to generate artificial experiments, each consisting of a perturbation (fixing the state of a set of nodes) and the observation of the state of a different node. We used 80% of the artificial experiments as a training set, i.e., to refine an initial model that had the same interaction graph as the actual model but its regulatory functions were randomly selected. The remaining 20% of the artificial experiments were used as a validation set to test the predictive power of the refined model. We describe the details of these benchmark studies in Text S6 and Table S2.

We found that the starting models had on average a 49% accuracy on the training set, and *boolmore* improved the models to 99% accuracy on average (Figure 2). Notably, *boolmore* also increased the accuracy of the models on the validation set from 47% on average to 95% on average. This indicates that *boolmore* does not overfit the training set and that the refined models give valid predictions.

**Figure 2.**
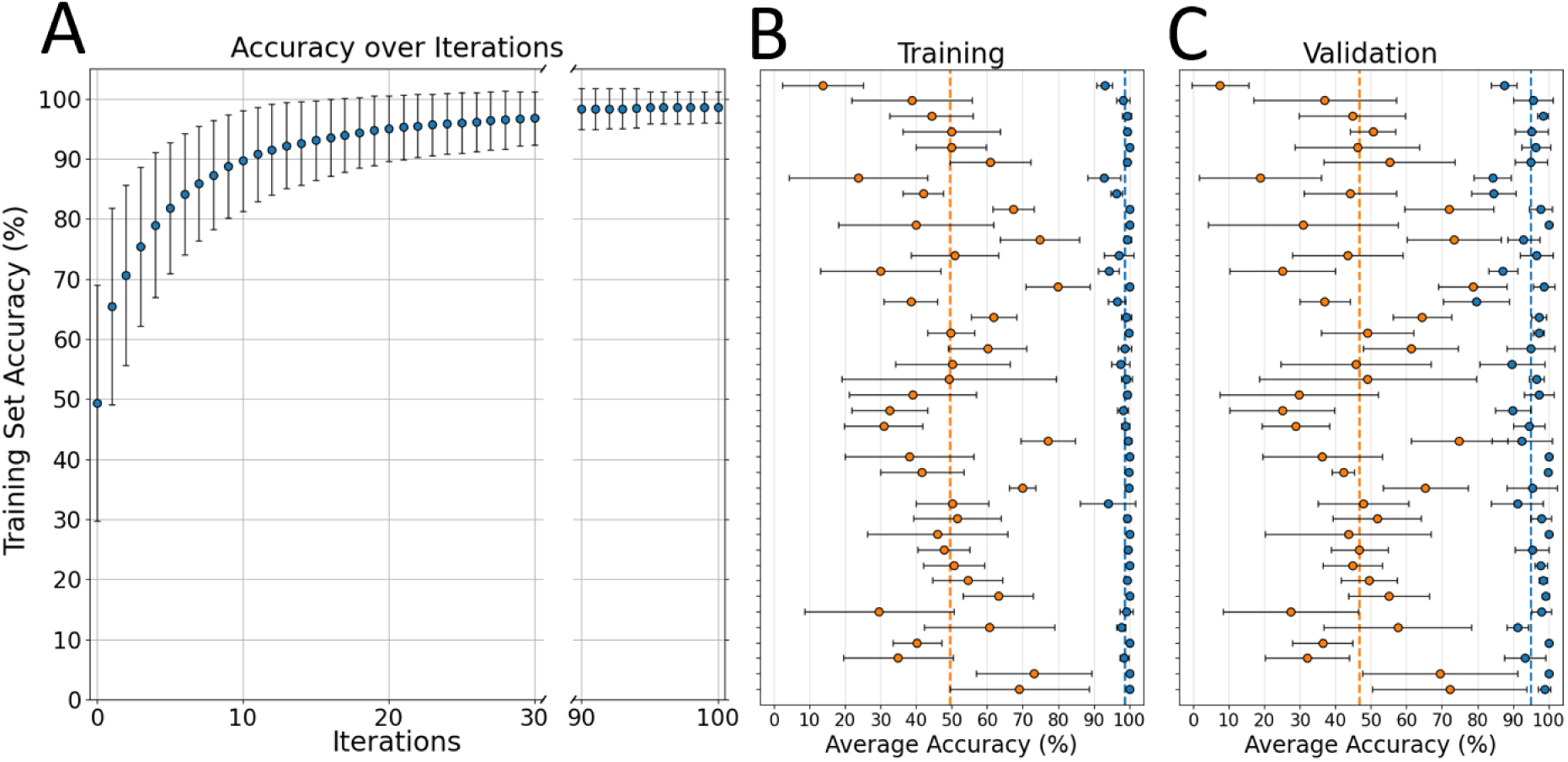
Results of benchmark studies using an ensemble of 40 published Boolean models. (A) Illustration of the accuracy improvement on the training set (80% of the artificial experiments) over the iterations of the algorithm. The blue circles represent the highest accuracy obtained at each iteration, averaged over 200 independent runs, with the error bars representing the standard deviation over the runs. (B) For each model the average accuracy on the training set (80% of artificial experiments) of 5 starting model variants is shown in orange, and the average accuracy of the 5 refined models is shown in blue. Error bars represent the standard deviation across the 5 replicates. The names of each model and their accuracies are listed in Table S2. Orange and blue dashed lines indicate the average accuracy of the starting models (49%) and the refined models (99%) respectively. (C) The average accuracies on the validation set (20% of artificial experiments) of the 5 starting model variants (orange) and the 5 refined models (blue). *Boolmore* had no information on the validation set in its model refinement process, but was still able to improve the accuracy greatly, from 47% on average to 95% on average.

### Improving the ABA-induced stomatal closure model

To go beyond simple benchmarks and use our approach to search for new biological insights, we apply our method in a case study. We selected a system that is representative of the challenges inherent in constructing a Boolean model of a complex biological phenomenon and keeping it up-to-date with current literature. Specifically, we analyzed a group of Boolean models, each aiming to integrate the mechanisms through which the hormone abscisic acid (ABA) leads to the closure of plant stomata (see Text S7 for the biological details of this process). The first model of ABA-induced closure, published in 2006 by Li et al. [28], included 42 nodes. This was expanded to 81 nodes in 2017 by Albert et al. [29]. An alternative expansion to 60 nodes was published in 2018 by Waidyarathne and Samarasinghe [38]. The 2017 model was refined in 2019 by the addition of a new edge and a simultaneous attractor-preserving reduction to 49 nodes [30]. We decided to use the 2017 model (whose node names are given in Table S3) as the basis of model refinement, as it is the most comprehensive and its analysis included a thorough comparison with 112 experiments. The reported accuracy of the 2017 model was 95/112=85%.

Despite the overall high accuracy, the 2017 model also exhibited two important weaknesses. First, there are 13 nodes that were shown experimentally to lead to closure when perturbed by external interventions. The 2017 model failed to recapitulate nine of these results according to the simulation-based criteria used at the time. Second, plant stomata reopen after the removal of the ABA signal, enabling the plant to resume photosynthesis. This reversibility of stomatal closure is not captured by the model.

Two follow-up publications aimed to address each of these two weaknesses of the 2017 model by making parsimonious hypotheses. Maheshwari et al. [30] hypothesized in 2019, and experimentally confirmed, the existence of an additional edge, thereby recapitulating five of the nine experimental observations of closure that were not captured by the 2017 model. A second follow-up to the model in 2020 [31] identified that the source of the irreversibility of stomatal closure in the model is the assumption of self-sustained activity of four nodes. The 2020 model achieved reversible closure and preserved the 2019 model’s success in capturing the five experimental observations of closure. We describe the 2017 model and its two follow-ups in more detail in Text S7. We emphasize that both of these improvements were proposed after an in-depth analysis of the 2017 model’s weaknesses and individual exploration of many hypotheses to address these weaknesses while preserving the strengths of the model.

We aimed to refine the 2017 model such that it reproduces the interventions that lead to stomatal closure and it exhibits reversible closure. Importantly, we aimed to do this without including these criteria as explicit constraints of the refinement algorithm, and instead we used the scoring process described in the Methods. We undertook an extensive search of the experimental literature and expanded the number of experimental observations to 505 (see Table S4). We considered two starting models, baseline A and baseline B, which correct an error in the 2017 model in two different ways. The key difference between the two models is that in the presence of ABA the baseline model A features the oscillation of multiple nodes, driven by transients in the elevation of the cytosolic Ca^2+^ level, while baseline model B defines an abstract node ‘Ca^2+^_c_ osc’ and leads to a fixed state for all nodes (see Text S8 for a description of the two baseline models). We used *boolmore* to find refined models that have an improved agreement with the corpus of 505 experiments performed for ABA-induced closure. In our application of *boolmore* on either baseline model, we adopted specific constraints for the regulatory functions of 26 nodes and specific criteria for allowing 13 additional, experimentally-supported edges, as we describe in Text S9. The algorithm added 8 edges from the pool (see the next section for more information).

It took roughly 10 hours to go through 100 iterations of the algorithm and generate 10,000 models on a PC with an AMD Ryzen 5 3600 6-Core CPU at 3.8GHz. The majority of the computation time was spent on model evaluation; the evaluation of each model took roughly 7 seconds on average.

The application of *boolmore* significantly improved both baseline models. We will refer to the genetic-algorithm-refined (GA-refined) baseline model A as GA1-A and the GA-refined baseline model B as GA1-B. We indicate the regulatory functions of the GA1-A and GA1-B models in Text S10. Our first specific model refinement goal was to reach better agreement with the experimental interventions that cause closure in the absence of ABA but were not recapitulated by the 2017 model.

### Improvement 1. Better agreement with experimental interventions that yield closure

Table 2 summarizes these experimental observations and the model results for the node Closure. The categorization of the experimental results, ranging from Some to ON, reflects the observed degree of closure (decrease of the stomatal aperture) induced by each intervention. The table indicates the results of the 2017 model, the Maheshwari et al. 2019 model, and the two GA-refined models. The GA1-A model preserves the 2017 model’s agreements and has an improved score for three additional responses. The GA1-B model preserves the agreement for three interventions, has a lower score than the 2017 model in one case (supplying InsP3), and receives a higher score for three additional interventions. Note that the elements whose closure-inducing nature is newly recapitulated by the

### GA-refined models lie at the core of the system, as reflected in the large number of experiments studying them

**Table 1.** Summary of the two sources of the improved score of the GA1-A and GA1-B models compared to the 2017 model [29]. The score gains (indicated in parentheses in the second column) of the GA-refined models arise from resolving 50% or more of the disagreements of the 2017 model with experimental results (third column) and achieving reversibility (fourth column). A small number of experiments were no longer captured by the refined models, decreasing the actual score gain (fifth column).

| Model | Score<br>(max 505) | Score gained from<br>resolving original<br>disagreements<br>(max +167.1) | Score gained from<br>reversibility<br>(max +27.4) | Score change from<br>new disagreements |
| --- | --- | --- | --- | --- |
| 2017 | 310.5 (61.5%) | - | - | - |
| GA1-A | 426.9 (84.5%, +116.5) | +101.4 | +25.3 | -10.2 |
| GA1-B | 407.9 (80.8%, +97.4) | +85.9 | +26.5 | -14.9 |

**Table 2.**
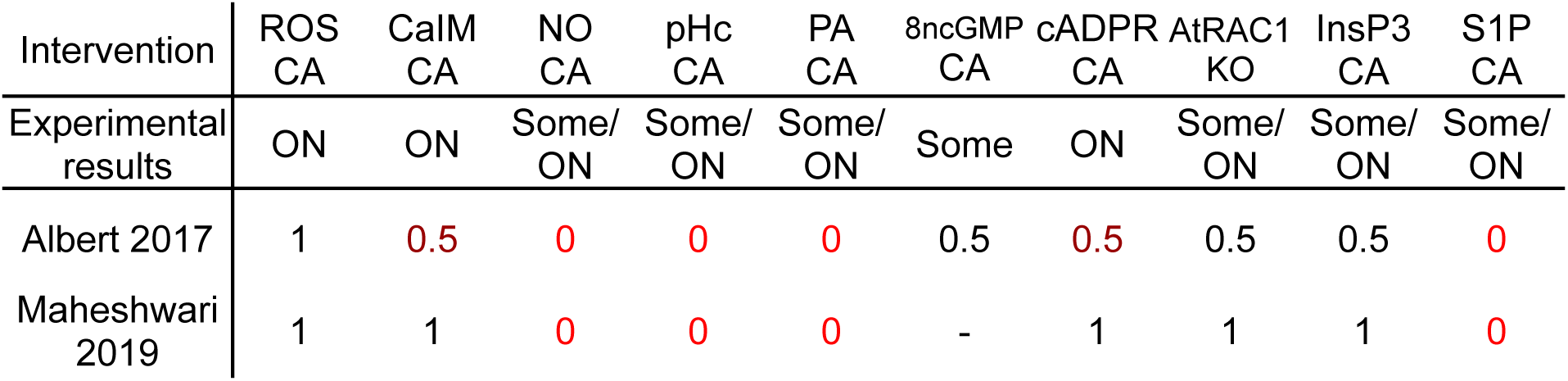
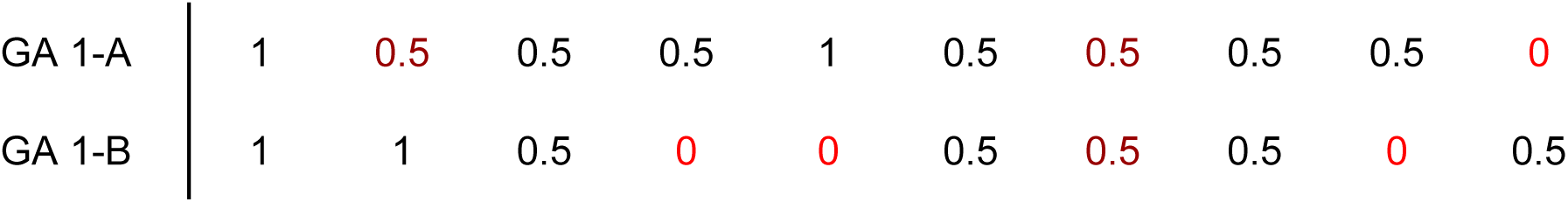
The experimental value of Closure in response to interventions in the absence of ABA and the corresponding predictions of four models. “CA” refers to constitutive activation or supply, which is implemented in the model by fixing the node in the ON state, “KO” refers to knockout, implemented by fixing the node in the OFF state. The interventions are listed in the decreasing order of the number of experiments that observed or manipulated the corresponding elements. The model entries indicate the categorization from lack of closure (0) to closure (1) described in the Methods. To aid the comparison of the models in recapitulating the experimentally observed closure responses, the entries are color-coded as black (agreement), dark red (partial agreement), or bright red (disagreement). The quantitative scoring of each result is indicated in Table S1.

### Improvement 2. Reversibility

The 2017 model has 17 minimal trap spaces in the absence of ABA; 16 with Closure=0 and 1 with Closure=1. In the presence of ABA, the model has a single minimal trap space with Closure=1. If the model starts in this trap space, then ABA is taken away, the model can only reach the single minimal trap space with Closure=1, and therefore fails to achieve reversibility. The elimination of this trap space would yield a one-to-one correspondence between the signal ABA and the closure response. *Boolmore*’s scoring method prefers models whose minimal trap spaces are aligned with the experimental observations.

Specifically, when an experimental observation corresponds to lack of closure (Closure=0) but the model result (i.e., the average value of the node Closure in the minimal trap spaces) is close to but not equal to zero, the model receives a partial score. Both GA1-A and GA1-B succeeded in eliminating the trap space with Closure=1 in the absence of ABA and thus achieving reversibility (see Table 1). It is particularly remarkable that this reversibility was achieved without the introduction of time-dependent regulatory functions as done in the 2020 revision [31].

### Improvement 3. Better agreement with experimental results

As 1 point in the score of a model means recapitulating the outcome of one experiment, the highest possible score for a GA-refined model is 505, the number of experiments in our compilation. The 2017 model recapitulated 314 experiments, out of which it received partial scores for 292 experiments, and disagreed with 191 experiments. An improved model would need to preserve the original model’s agreements with the experimental results and reach agreement with the experiments the original model did not capture. Indeed, we confirmed that the increase in the score of both GA1-A (from 61.7% to 84.5%) and GA1-B (from 36.5% to 80.8%) is due to resolving the majority of the original model’s discrepancies with experiments (see Table 1). An additional source of score increase is due to the elimination of the trap space with Closure=1 (as described above). Note that the GA-refined models were not able to gain the maximum score improvement from reversibility due to trade-offs in capturing some of the experiments (i.e., due to the fact that achieving agreement with one experiment may create a disagreement with another experiment).

To place these results into context, we present the accuracy of the previous manually constructed models as well as two additional versions of GA-refined models in Figure 3. In the GA0 models, we did not allow the addition of edges, therefore the GA0 models stay consistent with the original interaction graph. In contrast, for GA2 models we allowed more assumed edges, without requiring experimental support. The figure indicates that all versions of GA-refined models surpass all published models, while at the same time decreasing the time needed for refinement.

**Figure 3.**
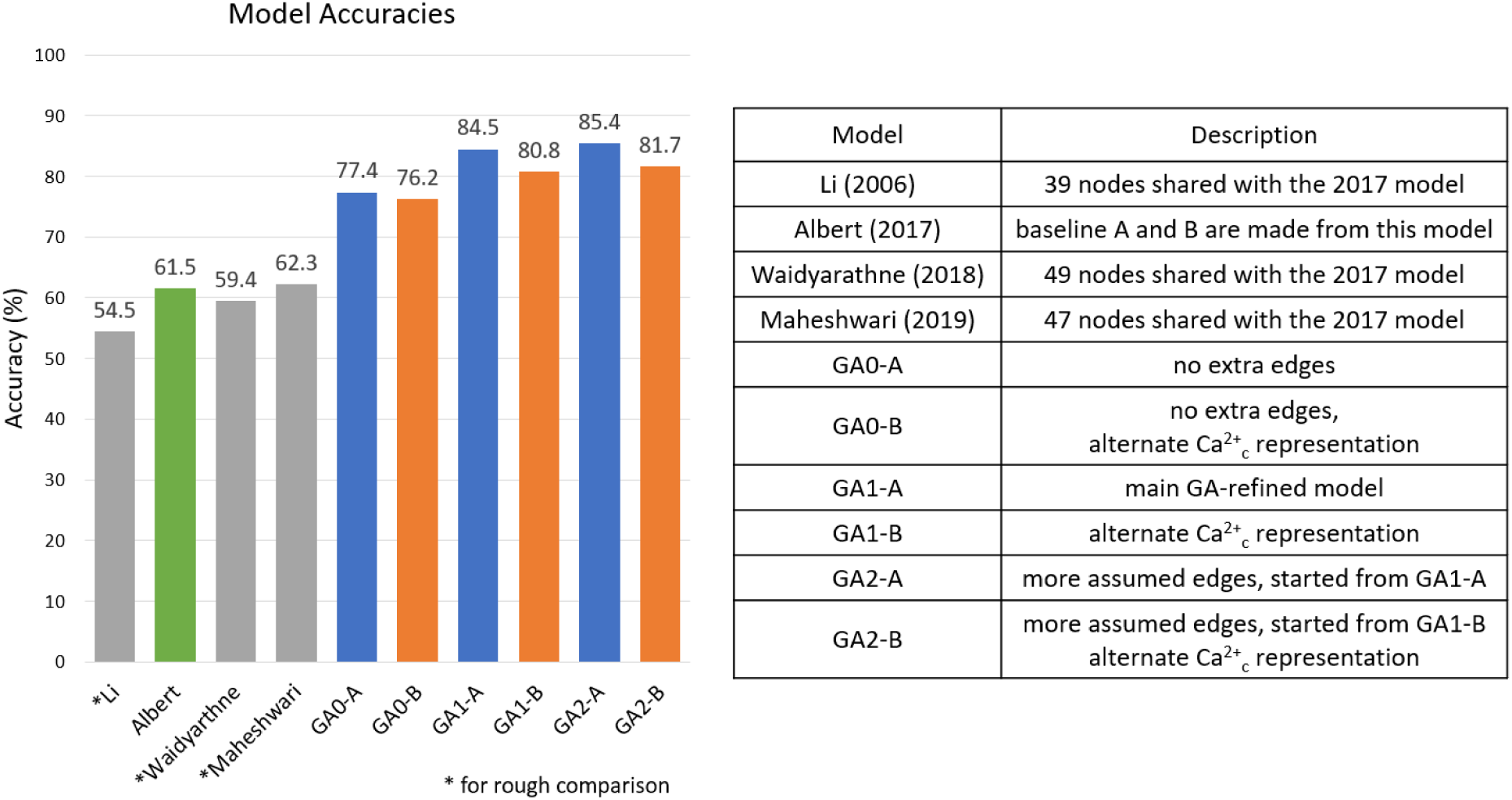
The accuracy of the previous ABA-induced stomatal closure models and the GA-refined models in reproducing a compilation of 505 perturbation experiments. The accuracy of the models marked with * is scaled to the percentage (50-75%) of experiments that apply to them. We note that 100% accuracy is not possible due to intrinsic limits on the agreement between the experiments on ABA-induced closure and Boolean models (see Text S11). Detailed results are in Text S12.

We evaluated the reproducibility of the genetic algorithm-based refinement by performing 16 independent GA1 runs starting from the baseline A model. The mean of the final scores was 81.5% with a standard deviation of 2.1%. The GA1-A model had the highest final score, 84.5%. There was a significant consensus among the GA1 model variants (e.g., 16 of the 35 functions that could change converged into an identical or logically equivalent form) as well as subtle dissimilarities that explain the differences in the score. The fitness score saturated before 100 iterations in each run. This shows that although the algorithm was not able to find the global maximum in every run, it was able to converge into a local maximum. This is a remarkable result considering the extremely high number of possible models for this case study. Indeed, we calculated using Dedekind numbers that the rough lower bound of the number of Boolean models consistent with the interaction graph and the constraints is 10^153^.

The goal of our model refinement workflow was to find biologically relevant improvements to an existing model, mirroring a manually refined model. Comparison with two publicly available tools for Boolean model inference or refinement, BoNesis [25] and Gitsbe [34], indicates that neither tool is able to achieve this goal for the ABA induced closure process (see Text S13). BoNesis could not return a single model within 24 hours even when restricting the experimental input to the experiments satisfied by a baseline model. All the models generated by Gitsbe in over 13 hours scored less than the worst-scoring *boolmore*-refined models. Furthermore, the Gitsbe-generated models contained many biologically invalid modifications to the regulatory functions, such as the loss of a key regulator of the node Closure.

As we describe next, the model refinements identified by *boolmore* have both explanatory and predictive power. The existence of two alternative baseline models allows the identification of changes implemented by *boolmore* in both models (see Text S14). These changes have a high likelihood of being biologically meaningful. In addition to an increased understanding of the biological system, it is also possible to extract novel predictions from the GA-refined models and suggest new experiments. In the following, we describe a selection of new predictions.

### New predictions of the GA-refined models of ABA-induced closure

#### Modifications to the interaction graph

Both GA-refined models feature significant changes to the interaction graph, which can serve as new predictions. GA1-A deleted 24 edges out of 152 starting edges. GA1-B deleted 17 edges out of 145 starting edges. Ten edges were deleted in both models; these edges are shown in Figure 4. A significant fraction of these edges represented assumptions of the 2017 model based on indirect evidence. These assumptions are no longer needed due to the improvements to the regulatory functions made possible by *boolmore* (see Text S14 for examples). In some of these cases, we could identify a shortcoming in the reasoning that led to the regulatory function in the original model. These findings illustrate how an automated method using a genetic algorithm can overcome modelers’ bias in selecting edges and functions, revealing optimal possibilities. Note that the deletion of an edge does not necessarily mean lack of influence; it indicates that in the context considered the influence is not significant enough to overcome the effects of the other regulators in a phenotypically relevant way.

**Figure 4.**
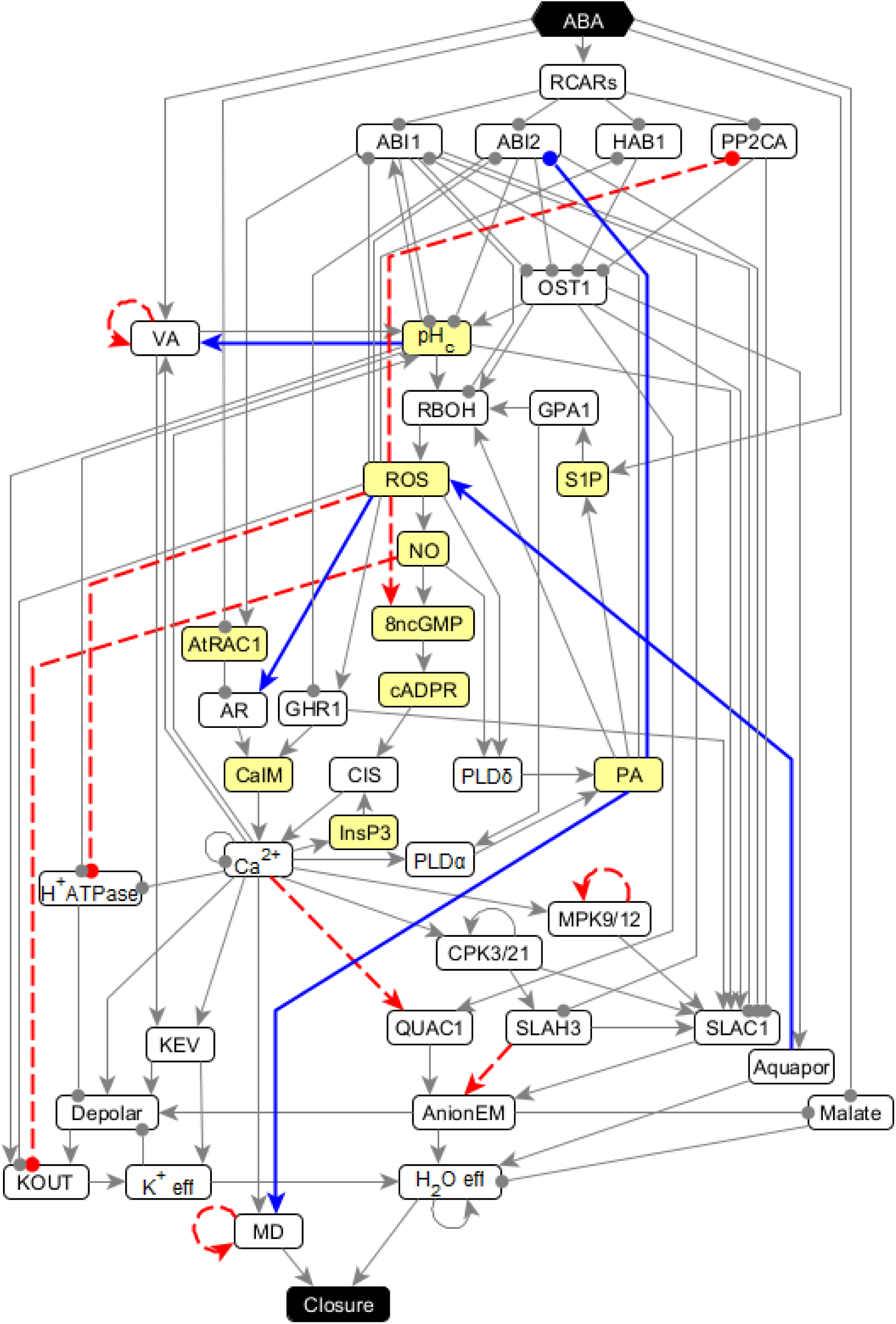
Simplified interaction graph illustrating the modifications to the network topology (edge deletions and additions) shared by the GA-refined models GA1-A and GA1-B. Each edge that terminates in an arrowhead indicates a positive regulation, and a round tip means negative regulation. Blue edges are shared additions of the GA-refined models, and red dotted edges were deleted by both GA-refined models. The nodes with yellow background are the key intervention nodes whose perturbation leads to some degree of Closure. The full names of the elements are indicated in Table S3.

GA1-A added 6 new edges and GA1-B added 7 new edges from the pool of 13 experimentally supported new edges. The added edges present in both GA-refined models indicate biological mechanisms that were not included in the 2017 model. As shown in Table 3, five new edges were added in both models. Importantly, the added inhibitory edge between PA and ABI2 recapitulates the experimentally supported prediction of the 2019 follow-up to the 2017 model [30]. The success of these shared additions confirms the improvements possible from the incorporation of new biological information.

**Table 3.** Added edges in the GA-refined models compared to the 2017 model [29]. Note that the inhibitory edge from PA to ABI2, which was incorporated in the 2019 revision of that model by Maheshwari et al. [30], was added by both GA-improved models.

| | PA $\dashv$ ABI2 | Aquaporin<br>→ ROS | ROS → AR | PA → MD | pH<sub>c</sub>→VA | Other Edges |
| --- | --- | --- | --- | --- | --- | --- |
| Maheshwari<br>2019 | O | X | X | X | X |  |
| GA 1-A | O | O | O | O | O | PA $\dashv$ HAB1 |
| GA 1-B | O | O | O | O | O | Ca<sup>2+</sup><sub>c</sub>osc $\dashv$ ABI2<br>AR → RBOH |

AR: Actin Reorganization, MD: Microtubule Depolymerization, VA: Vacuolar Acidification

#### Predictions that can be verified experimentally

The GA-refined regulatory functions also serve as new biological predictions. We summarize selected testable predictions in Table 4 and explain them in Text S15. Here we illustrate the types of predictions with a few examples.

The GA-refined models modify the regulatory function of the anion channel SLAC1 such that it is easier to activate. As a consequence, they recapitulate the experimental observation that ROS activates SLAC1. A follow-up prediction is that the resulting anion flow brings the malate concentration below threshold.

Causal relationships mediated by chains of interactions (pathways) can also yield new predictions. The 2017 model and the GA-refined models agree in predicting that ROS is sufficient to induce PA production. This is experimentally testable. As the GA-refined models incorporate the new observation that PA is sufficient for microtubule depolymerization, a follow-up testable prediction is that ROS can induce microtubule depolymerization; this can be tested by methods used in [39].

The shared features of the GA-refined models’ minimal trap spaces, which are described in detail in Text S16, identify further predictions. One such prediction is that external Ca^2+^ would yield no or a very limited amount of ROS production. While ROS production in ABA-induced closure has been experimentally documented, the production of ROS in response to external Ca^2+^ has not yet been studied experimentally.

**Table 4.** Illustrative biological predictions that can be made based on the GA-refined models of ABA-induced closure.

| Context | Prediction |
| --- | --- |
| Constitutive activation of ABI1 in the presence of ABA | The aquaporin channels will not activate (will not open) |
| Externally provided ROS in the absence of ABA | The malate concentration decreases below threshold. |
|  | PA is produced |
|  | The microtubules are depolymerized |
| Disruption of aquaporins (*pip2;1* KO) in the presence of ABA | Lack of NO production |
|  | No cADPR production |
| | PLDδ is not activated |
| Externally provided ROS in the absence of ABA | PP2CA is active |
| Externally provided ROS, knockout of RCARs receptors, in the presence of ABA | PP2CA is active |
| Externally provided cADPR in the absence of ABA | pH<sub>c</sub> increases to a lesser extent than in response to ABA |
|  | No ROS production |
|  | PA is produced |
| Externally provided 8-nitro-cGMP in the absence of ABA | pH<sub>c</sub> increases to a lesser extent than in response to ABA |
|  | No ROS production |
|  | PA is produced |
| Externally provided Ca<sup>2+</sup> in the absence of ABA | No ROS production |

A more general insight can be gained from observing that GA1-A and GA1-B rely on two different mechanisms to yield minimal trap space results of 0.5 that achieve agreement with experimental observations classified as “Some” in Table 2. The attractors of GA1-A feature oscillations in Closure along with a significant number of other nodes, which are due to the oscillations of Ca^2+^. In contrast, GA1-B has two attractors, one featuring Closure=1 and the other featuring Closure=0. Although different in a technical sense, these results are consistent with each other in that they both suggest population-level heterogeneity of the stomatal responses. Due to the challenges of tracking individual stomata in real time, there are few studies of individual stomata. Nevertheless, the studies that exist include observations of multiple types of oscillations in the stomatal aperture, including Ca^2+^-induced oscillations (reviewed in [40]). In addition, [28] reported a significant bimodality of the ABA-treated stomatal aperture distribution. Our results suggest that more in-depth analyses of the time-dependent status of signaling mediators in individual guard cells may reveal a richer dynamic picture than previously thought.

## Discussion

Small, local changes in a Boolean model, such as to the regulatory function of a single gene, can lead to global changes in the model’s attractor repertoire and thus to the predicted phenotypes. This makes the process of iteratively building a model, or incorporating new biological knowledge into an existing model, extremely challenging. Here we automate the process of refining or updating an existing Boolean model and implement it as the tool *boolmore*. We use a genetic algorithm to adjust the regulatory functions of the model to improve its agreement with curated experimental results. Automated exploration and quantitative scoring of modeling decisions can alleviate the immense cognitive burden of integrating myriad experiments into a causal model, while reducing human error and modeler bias. *Boolmore* uses a flexible scoring method that appropriately handles low-throughput state data and the subtleties that arise in discretizing experimental data. In the case study considered here, for example, we used this flexibility to introduce additional outcome categories (e.g., “Some”, “OFF/Some”) to describe intermediate outcomes (e.g., reduced closure) and inconsistency between repeat experimental observations. By considering the interdependence of perturbation experiments, our scoring achieves an unprecedented level of biological realism even with limited data.

Our method allows modelers to automatically explore many more modeling choices than previously possible, enabling systematic evaluation of alternate modeling assumptions (e.g. compressing an oscillating negative feedback loop into a single non-oscillating node, or introducing multiple levels of activation for a node). Such core modeling decisions can have large, global effects on the model dynamics, and hence switching to a different description of an important node’s regulatory logic often requires many adjustments to other nodes’ functions to reach an accuracy comparable to that of the original model. While it is difficult and time-consuming to manually evaluate and compare multiple core decisions, *boolmore* allows such adjustments to be made in a reasonable amount of time. As an illustrative example, we considered a model of ABA-induced closure in which the oscillating negative feedback between cytosolic calcium and calcium ATPase was replaced by a single node whose activity indicates calcium oscillation (baseline model B). Using *boolmore*, we refined baseline model B, which had an accuracy of only 36.5%, into model GA1-B, with an accuracy of 80.8%.

Broad exploration of modeling choices can be further enhanced by workflow customization. The scoring system can be readily augmented by additional measures, such as the number of attractors, or the phenotype transitions under different environments, to produce more realistic models. Model prioritization can be customized so that only models with certain key behaviors are selected. Furthermore, the modeler has the freedom to determine how individual experimental results should be encoded; for instance, allowing for finer-grained response categories than used here.

There is an intrinsic limit to the degree to which a Boolean model can recapitulate quantitative experimental results (see Text S11 for specific examples in our case study). These limitations would be ameliorated by introducing multiple levels of activation for nodes. As a lossless mapping between a multi-level variable and a set of Boolean variables has been worked out [41,42], *boolmore* can be expanded to accommodate multiple levels. We indicate in Text S17 the details of this adaptation and a proof of concept application to a model of nutritional regulation of lisosomal lipases in *C. elegans* [43]. A multi-level representation of the output variable “Closure” is especially needed in future models that integrate the response to the various signals that lead to various degrees of stomatal closure. These signals include ABA, high CO_2_, darkness, and their combinations. *Boolmore* can be a useful tool for identifying the number of levels that yield the best agreement with experiments.

Though *boolmore* worked well on models with randomized regulatory functions, it performs best when it starts from a pre-existing baseline model with reasonable regulatory functions. Starting with a high-quality interaction graph is especially important, as it can fundamentally constrain dynamical behaviors regardless of the Boolean functions. We found that a change in a single edge may lead to a great decrease in the agreement of the model with experimental results. When we corrected the 2017 model of ABA-induced stomatal closure by deleting the erroneous edge from Vacuolar acidification to KEV, *boolmore* could not refine the model to recapitulate ABA-induced closure without adding a self-edge on H_2_O efflux to compensate. Fortunately, many methods have been developed to construct or infer biomolecular interaction graphs [44,45], and curated interaction graphs are available in multiple databases [46–48], increasing the likelihood of a high-quality interaction graph for any cellular process to be modeled.

New discoveries that refine the interaction graph will improve the final fitness of the GA-refined model. We illustrate this trend by evaluating 3 versions of GA-refined models (see Figure 3 and Text S12). The GA0-A and GA0-B models, which did not allow edge addition, showed an overall 5.8% lower score than the GA1 models. We also considered GA-refined models that allowed 50% more additional edges than the 13 experimentally observed edges. We chose the assumed edges manually to help achieve agreement with top-of-the-hierarchy experiments. Two such edges were incorporated in the GA-improved models GA2-A and GA2-B, both of which suggested new regulation of ion flows. For example, the suggested connection between CIS and anion flow allows the recapitulation of impaired anion flow in case of disruption of PLC or cADPR. The GA2-A and GA2-B models showed a 1.0% increase in scores on average, suggesting the existence of undiscovered interactions. This example shows that our workflow can be used to explore putative interactions whose incorporation can enhance model accuracy.

Although gaps of knowledge in the interaction graph can be filled by allowing the addition of edges, such addition is only effective in moderation; each additional edge doubles the size of the search space. Although *boolmore* can in principle search the whole space of Boolean functions consistent with the constraints, it can only search the local vicinity of the starting functions in the practical time limit. Model evaluation is the major computational bottleneck of *boolmore* and we believe that no trivial speedup is possible. By relying on the Python package *pyboolnet* [13], *boolmore* could find the minimal trap spaces in our 82-node case study in a hundredth of a second. Even with this remarkable speed, evaluating more than 8000 models under more than 260 perturbation settings resulted in 10 hours of runtime for a single run on a personal computer.

Any method that involves fitting to data has a chance of overfitting, i.e., inferring more parameters than can be justified by the data [49]. The parameters of a Boolean model are the Boolean functions of individual nodes. In our case study, *boolmore* deleted many edges that were included in the manually constructed starting models. Deletion of edges is analogous to removing parameters, and hence our workflow actually helped reduce the risk of overfitting that exists in the manual refinement process. Our workflow also helps introduce new edges in a much more conservative manner by only allowing edges that increase the accuracy of the model as a whole. The predictive power of the models refined by *boolmore* is evident in the benchmarks, where the refined models showed 95% accuracy over the validation set, which was not used in the refinement process.

Our automation of model refinement streamlines model-building by lowering the hurdle for the initial model. Once the experimental database and an interaction network for the model are set, a quickly achievable preliminary model can be refined in a fraction of the time that would be needed for manual model building. We envision that *boolmore* may also be used as an exploratory tool, allowing biologists and modelers to evaluate whether, and how, a model can accommodate hypothetical experimental results. *Boolmore* can also be integrated with other steps such as automated evidence gathering [50] and model expansion [51] to progress toward fully automated and extremely fast model construction.

## Supporting information

**Text S1**. Key concepts of Boolean modeling and key methodologies to identify the long-term behaviors of a Boolean system

**Text S2**. Summary of prior work on Boolean model inference or revision

**Text S3**. Methodological details of *boolmore*

**Text S4**. Details of the interpretation of the “Some” category of experimental results

**Text S5**. Parameter analysis

**Text S6**. Details of the benchmark analysis used to test the overall efficiency of *boolmore*

**Text S7**. Detailed description of abscisic acid-induced closure and its Boolean modeling

**Text S8.** Correcting an error in the 2017 model and choosing baseline models

**Text S9**. Details of the constraints and extra edges used for improving the ABA-induced closure model

**Text S10**. The Boolean functions of the 2017 model and of the two GA-improved models

**Text S11.** Intrinsic limits on the agreement between experiments on ABA-induced closure and models

**Text S12**. Summary of the scores and features of the genetic algorithm-modified models as compared to the 2017 model of ABA-induced stomatal closure

**Text S13**. Considering BoNesis or Gitsbe as alternatives to derive a Boolean model from the experimental data on ABA-induced stomatal closure

**Text S14**. Regulatory function revisions shared by the two GA-improved models

**Text S15**. Details of the experimentally testable biological predictions presented in Table 4

**Text S16**. The dynamics of the GA-refined models

**Text S17**. Adapting *boolmore* to multi-level models

**Table S1**. The agreement functions used in our workflow

**Table S2**. Detailed benchmark results

**Table S3**. The full names of the abbreviated node names in the ABA-induced closure models

**Table S4**. Compilation of experimental results used to score the ABA-induced closure model

## Acknowledgements

We are grateful for the advice of Prof. István Albert, Prof. Sarah M. Assmann, Prof. Dezhe Jin, and the helpful suggestions of Dr. Xiao Gan, Dr. Fatemeh Sadat Fatemi Nasrollahi, and Dr. Eli Newby.

## Data availability statement

The Python package *boolmore* is available in the github repository https://github.com/kyuhyongpark/boolmore.

