## Supporting Information for "Automated Model Refinement Using Perturbation-Observation Pairs"

### Text S1. Key concepts of Boolean modeling and key methodologies to identify the long-term behaviors of a Boolean system

#### Boolean model construction

In systems for which the information-gathering is piecewise, model construction is done via manual integration of distinct pieces of experimental evidence. Experimental observations of interactions (e.g. chemical reactions and protein-protein interactions) and positive or negative regulation (e.g. transcriptional or allosteric regulation) inform the edges of the network. Another type of experimental evidence is a single observation that is nevertheless influenced by the whole system (e.g., it indicates that the knockout of a gene impairs the system's response to a signal). It is not possible to directly assimilate this type of global evidence into any individual node's regulatory function, as the function only incorporates the effects of the node's direct regulators (i.e., it is purely local). Although some parts of the model can be directly constrained by preexisting experiments, usually many degrees of freedom remain. Fully exploring the space of all regulatory functions that are consistent with local information and systematically testing the agreement of the resulting model with global system-level data is prohibitively time-consuming. In practice, modelers often tackle this challenge through a process of trial and error informed by the insights of domain experts.

#### Biological predictions of Boolean models

Boolean models of biomolecular systems successfully capture key behaviors of the system, and make useful predictions such as identifying master regulators or drug targets. An example of capturing key behaviors is a Boolean model of T cell differentiation in response to the cellular microenvironment which recapitulated 8 documented T cell types such as helper and regulatory T cells [8]. An example of the predictive power  of Boolean models is the model of the cell fate change that represents the first step toward metastasis in liver cancer, which identified the interventions that are able to prevent or reverse this fate change [9].

#### Updating Boolean models

Experiments performed after the construction of a model may disagree with the model, or identify important new elements that the modeler would like to capture in the model. Keeping a hand-constructed model up-to-date involves the same trial and error iteration as its construction. For example, the earliest version of a model of T cell differentiation, introduced by Luis Mendoza, initially had 17 nodes and reproduced the three T helper cell types known at the time [52]. After the identification of T regulatory cells, the model was expanded to 35 nodes [53] and reproduced five T cell types. After a significant expansion to 85 nodes, and a subsequent logic-preserving reduction, the model reproduced eight T cell types [8].

#### Analyzing Boolean models

Boolean models typically use a discrete and implicit time variable. Frequently used implementations are synchronous update, in which every node is updated at each time step, and asynchronous update, in which a single randomly selected node is updated at each time step. The biological predictions of models are typically, but not always, robust to the update scheme [54].

The state transition graph, made up by the $2^N$ states of the system and the possible, update-scheme-dependent, transitions between them, provides a complete description of the model's dynamics, including the attractors and the states that lead to them. As the state transition graph is computationally expensive to calculate, modelers often choose to run simulations, which can be thought of as sampling trajectories (random walks) in the state transition graph. An effective alternative method to obtain a summarized state transition graph and determine the attractors of a Boolean model is via the identification of so-called stable motifs [55]. A stable motif is a self-sufficient feedback loop that can sustain an associated state of its constituent nodes regardless of the other nodes of the system. A conditionally stable motif (CSM) is a motif that acts like a stable motif as long as certain conditions are satisfied, for example, as long as a node external to the motif maintains a fixed state [56].

A subspace of a Boolean network is a subset of the state space characterized by a set of fixed node values, and a subspace is called a trap space if there are no dynamical trajectories that exit it. Once locked in, a stable motif is irreversible, and traps the system into a trap space [57]. A minimal trap space is a trap space that does not contain any smaller trap space within it. The successive locking-in of stable motifs and CSMs traps the system into smaller and smaller trap spaces, eventually reaching a minimal trap space [55,57]. A minimal trap space, also called quasiattractor, is a good approximation of an attractor. It is either a single state (a point attractor) or a set of states in which the nodes that are not fixed by stable motifs may oscillate between the 0 and 1 state. Calculating trap spaces is much faster than calculating attractors. In rare cases, there may be multiple complex attractors in the same minimal trap space, or there may exist motif-avoidant attractors that lie outside of a trap space. These behaviors are dependent on the update scheme [57,58], and are unlikely in biological systems, which need to function robustly despite variations and stochasticity.

### Text S2. Summary of prior work on Boolean model inference or revision

As our goal is to refine an existing Boolean model using a genetic algorithm, the prior research on Boolean model refinement (by any method) or on Boolean model inference using genetic algorithms is the most relevant. We categorize this prior work based on the goal of the authors to infer a network together with a Boolean model, infer a Boolean model using a directed and signed prior knowledge network, or refine/extend an existing Boolean model. We organize the key information about each algorithm in the table below, listing the type of state input they require, whether the key methodology is a genetic algorithm (GA), answer set programming (ASP), or clause learning (CL), whether or not they use a prior network, and describe a representative case study in which the algorithm was used.

Almost all algorithms (with the exception of those by Sayed et al. [59] and Flobak et al. [34]) require system-wide input information, i.e., the knowledge of the state of all the nodes, in multiple conditions, including the wild type (WT) system and a variable number of node knockouts (KOs) and constitutive activations (CAs). When the number of conditions is of the same order of magnitude as the number of nodes, we refer to the state input as a "matrix". (Indeed, fewer conditions than the number of nodes make the model inference problem under-determined). Most studies assume that each state in the input reflects a point attractor (steady state). Four studies allow multi-state (oscillating) attractors as well, in which case we write “attractor”.

Due to the different goals, the biological comparisons through which the output of the algorithm is evaluated also differ. If the goal is to infer a network together with a Boolean model, the accuracy of the output is mainly tested by evaluating the resulting edges. Most of the existing algorithms for Boolean model inference (with Bonita [22] being a notable exception) evaluate the model solely on its attractors. The two ASP-based algorithms for model revision (ModRev [35] and ARBoLoM [36]) are solely tested on their ability to recapitulate the input (steady states or trajectories of the wild type system). It is important to note that these two model revision algorithms are not set up to handle perturbation data. It is evident from the last column that almost all studies (with the exception of Terfve et al. [33] and Flobak et al. [34]) used only artificial measurements to evaluate the algorithm and did not apply it to real data.

| Reference | State input | Key method | Prior network | Representative case study and result |
|---|---|---|---|---|
| Goal: Inference of an interaction network and Boolean model | | | | |
| Trinh & Kwon[18] | Matrix of point attractor(s) of the WT system and for multiple perturbations | GA | No prior network. | Artificial gene expression data generated from a 100 node network. More than 100 point attractors were used, including the point attractors of the WT system and the point attractor(s) in case of KO of each gene. The inferred network had a structural accuracy of ~95%; the regulatory functions were not evaluated. |
| Goal: Inference of a Boolean model | | | | |
| Dorier et al. [19] | Matrix of point attractors of the system and the transitions between them in case of perturbations | GA | The inferred network is a subset of a prior network. | Artificial measurements generated from a published 25-node Boolean model of cell fate decisions. The input consisted of 256 transitions between point attractors; the starting attractors are the four point attractors of the WT system and both KO and CA of each node are considered. The inferred model recapitulated the attractors obtained from individual KO of 14 key nodes. |
| Ghaffarizadeh et al. [20] | Attractors of the WT system and for certain perturbations | GA | If used, the prior network is fixed. The regulatory functions are assumed to be nested canalyzing functions. | Artificial expression profiles generated from a 11-node published Boolean model of myeloid cell differentiation. The edges of this model were used without information on their signs. The input contained four point attractors corresponding to four cell types and two constraints indicating the reduction in available cell types in case of KO of a certain gene. The inferred edge signs agree with prior literature on 26/30 edges. |
| Munoz et al (Griffin) | The attractors of the WT | CL | The prior network can include | The network structure and set of 10 point attractors of a previously published |

| | | | | |
|---|---|---|---|---|
| [23] | system, possibly expanded by attractors obtained in case of perturbations | | hypothetical regulators and ambiguous effects | 13-node Boolean model of *Arabidopsis thaliana* flower development are consistent with more than 300,000 Boolean models. |
| BoNeSis[25] | Time course information that reflects trajectories from an initial state to different attractors. The experimental observations do not necessarily need to include all the genes. | ASP | The prior network is fixed. | A network of 12 genes that determine the differentiation of neural stem cells into neurons, astrocytes, and oligodendrocytes. There are 9 observations of 5-12 genes, information on 6 positive reachability properties (e.g. the initial state in which Pax6 is active only reaches the state in which 5 marker genes are also active) and on 3 negative reachability properties (the state in which every gene is inactive cannot reach three of the other states). The paper identifies thousands of inferred networks consistent with this information. |
| CellNOpt/CNO [33,60] | Matrix of point attractors of the WT system under certain signals and perturbations | GA | The prior network's edges can be pruned; new edges can be added one by one | Phospho-proteomics measurements of 16 proteins in response to one of 7 external molecules and for 7 cases of inhibition of a single protein. The prior knowledge network was reduced to include only the measured or inhibited nodes. The inferred model agreed well with experiments on combined inputs (to the same extent as with the input data). |
| Müssel et al. (CANTATA) [21] | Time series or attractors of the system | GA | The prior network's edges can be pruned; new edges can be added randomly | Artificial measurements generated from a 10-node published Boolean model of the yeast cell cycle. The input consisted of the two most relevant point attractors and the most important sequence of 10 states. The actual network was modulated by 5 random changes. The inferred models reproduced the training set; the networks had a mean accuracy of ~98%. |
| Palli et al (Bonita) [22] | Matrix of states of the system under various conditions | GA | The prior network is fixed. Each observed state is assumed to be a steady state (point attractor). | Artificial measurements generated from a 108-node KEGG pathway combined with biology-inspired random expression. Ninety percent of the inferred model's regulatory functions were equivalent to the functions used to generate the artificial measurements. |
| Goal: Extension or revision of an existing Boolean model | | | | |
| Sayed et al [59] | The steady states of key nodes under various | GA | A group of new edges (each with one or two new nodes) are added | A previously published 39-node Boolean model of T cell signaling was made incomplete by removing 8 nodes. An ensemble of candidate extensions that |

| | | | | |
|---|---|---|---|---|
| | conditions | | to the prior network. | contained the removed nodes was analyzed. The input was obtained from the original model and consisted of the expression of five key nodes under two levels of an input signal. The inferred models restored at least four of the removed nodes and recapitulated the input. |
| Azpeitia et al [61] | The attractors of the WT system | GA | New edges are added to the prior network one by one. The new edges satisfy biological constraints. | After updating a previously published 10-node Boolean model of the gene regulatory network of the root stem cell niche with newly discovered interactions, it did not recapitulate two of the nine known attractors (all of which are point attractors) and had non-biological attractors. The application of the algorithm included 3 new interactions and yielded 10 very similar models that recapitulate all 9 attractors and have only 3 non-biological attractors each. |
| Gouveia et al. (ModRev)[35] | The point attractors of the WT system | ASP | Edges of the prior network can be deleted or change sign. New edges can be added. | A previously published 40-node Boolean model of T cell receptor signaling was made incorrect by probabilistically changing regulatory functions, changing edge signs, removing or adding edges. The algorithm-revised model could recover the point attractors of the original model for the first three types of error, but timed out in most instances of added edges. |
| Aleixo et al (ARBoLoM)[36] | The point attractors or several trajectories of the WT system | ASP | Edges of the prior network can be deleted or change sign. New edges can be added. | A previously published 40-node Boolean model of T cell receptor signaling was made incorrect by probabilistically changing regulatory functions, changing edge signs, removing or adding edges. The input consisted of the point attractors of the original model, or up to 5 trajectories with up to 20 timepoints. The algorithm-revised model could recover the correct point attractors or trajectories. |
| Flobak et al. (Gitsbe)[34] | The steady states of a list of biomarkers of the WT system, possibly expanded by their states in case of perturbations. | GA | Edges of the prior knowledge network can be deleted. | The 144-node interaction network of a previously constructed Boolean model of growth-promoting signaling in the AGC gastric adenocarcinoma cell line, as well as Boolean functions following a default template, served as the starting point of a genetic algorithm-based model calibration. The model was trained on the observed states of 21 biomarkers in proliferating AGS cancer cells. The calibrated model was used to predict synergistic drug combinations, recovering the results of a drug screen with an ROC area under the curve of 0.69. One predicted synergistic combination was tested and validated |

| | | | | |
|---|---|---|---|---|
| | | | | experimentally. |

As Boolean models of biological systems are judged on their explanatory and predictive power for the biological process they describe, the same standards should be used to evaluate inferred Boolean models as well. If a prior network is used, the accuracy of the model can be tested by evaluating its regulatory functions as well as its dynamical outcomes. If algorithm-driven additions to the prior network are allowed, these should be evaluated against the biological literature and may represent experimentally testable predictions. Such evaluation has not been a focus so far.

### Text S3. Methodological details of *boolmore*

#### Mutating functions such that edge signs are preserved

The interactions and regulatory relationships in biological networks are locally monotonic in the vast majority of cases. This means a regulator either inhibits or activates its target; it is not an inhibitor in one context (i.e., for a certain state of other regulators of the target node) and an activator in another context. The signs of the interactions are built from the literature, and form the foundation of the model. These interactions are often well established and hence changing the signs will lead to completely unrealistic networks.

To achieve random mutations of Boolean functions that preserve the original signs, we propose a degenerate binary representation of each function based on a disjunctive normal form of the function. The key idea is the following: A function with p positive regulators and n negative regulators can be expressed (not necessarily uniquely) as the disjunction (OR composition) of a subset of the $2^{n+p}$ conjunctions (AND compositions) consistent with the regulatory signs. Each of these conjunctions is assigned a location in a binary string of length $2^{n+p}$; the binary string is interpreted as the disjunction of the conjunctions corresponding to the locations in which the string has a 1. Each mutation changes a randomly selected digit of this binary representation and is guaranteed to preserve the signs of the regulatory relationships.

The specific representation for a positive regulatory function (i.e., one for which all regulators are activators) with k inputs is

$$\begin{aligned} f(X_1,\ldots,X_k) = \; & a_{\{\}} \;|\\ & a_{\{1\}}\&X_1 \;|\; \ldots \;|\; a_{\{k\}}\&X_k \;|\\ & a_{\{1,2\}}\&X_1\&X_2 \;|\; \ldots \;|\; a_{\{1,k\}}\&X_1\&X_k \;|\; \ldots \;|\\ & a_{\{2,3\}}\&X_2\&X_3 \;|\; \ldots \;|\; a_{\{2,k\}}\&X_2\&X_k \;|\; \ldots \;|\\ & \vdots\\ & a_{\{1,\ldots,k\}}\&X_1\&\ldots\&X_k \end{aligned}$$

Here $X_i$ represents the state of the $i^{th}$ input node out of k input nodes. The notation "|" means logical "OR" and "&" indicates logical "AND". A constant Boolean coefficient $a_S$ is assigned to each subset S of input nodes, from S={} to S={1,...,k}. Note that this representation is not unique. The coefficients $a_S$ can be ordered to obtain a binary representation of the function. A natural ordering interprets each S as the binary representation of the numbers from 0 to $2^k$. This ordering is also traditionally used in the truth table representation of Boolean functions.

For example, let us consider three variables A, B, C and the function f(A,B,C) = B&C | A&B. This function can be written as f(A,B,C) = 0 | 0&C | 0&B | 1&B&C | 0&A | 0&A&C | 1&A&B | 0&A&B&C. Note that this representation includes all the possible subsets of ABC, leading to $2^3$=8 clauses; the original three clauses are the terms that start with 1. The update function can be represented by a string of the $a_S$ coefficients, i.e., f(A,B,C) is represented by 00010010. The benefits of this representation are that any possible combination of 0s and 1s represents a positive function, and that the combinations span all the positive functions. Exemplifying the degenerate nature of this representation, 00010011 also represents f because A&B&C is implied by B&C | A&B. However, the representation that has the maximal number of 1s (max representation) is unique and is equivalent to the truth table of the function (00010011 in this example). The representation that has the minimal number of 1s (min representation) is also unique and is equivalent to the Blake canonical form of the function (00010010 in this example).

To obtain a mutated function, each digit of this representation is changed with a certain probability. For example, the mutation may change the sixth digit and lead to 00010110. The mutated function is thus f’(A,B,C) = 0 | 0&C | 0&B | 1&B&C | 0&A | 1&A&C | 1&A&B | 0&A&B&C, which can be further simplified to f’(A,B,C) = B&C | A&C | A&B.

This method can be extended to mutate locally monotonic functions that contain the NOT operator (which we will represent as !). If g(A,B) = !A&B, we can use the change of variables A’ = !A and represent the function as the positive function g’(A,B)=A’&B. As long as we keep the original records of the signs, any locally monotonic function can be switched to a positive function, mutated, and switched back.

Note that the above representation naturally has a bias toward 1. For example, if the first digit of the binary representation is mutated to 1, the whole function becomes 1. To remove this bias, we also consider a binary representation of the negation of the function. For example, the negation of the above function, !f(A,B,C) = !B | !A&!C = !f(A’,B’,C’) = B’ | A’&C’ can be represented as 0010100. In the negated representation, mutating the first digit to 1 makes the negated function 1, and thereby makes the function 0. To remove bias toward one output state or the other, we introduce a 50% chance to mutate the negation of the function rather than the function itself.

**Preserving known mechanisms via constraints**

In some cases, the biochemical mechanisms of certain regulatory relationships are known. We encode such knowledge as constraints to the regulatory functions, in a similar vein as [61]. These constraints not only ensure reasonable models, but they also reduce the search space greatly. For example, if it is known that the activation of a regulator A is necessary for the activation of the target B, then the form of the update function of B is constrained to be “$f_B$ = A & (other regulators)”. For a function with 4 inputs, this reduces the number of possible Boolean functions with fixed signs from 168 to 20. The constraints of the regulatory functions are enforced by their binary representations. For example, if A is constrained to be a necessary regulator, any term that does not contain A will have a coefficient of 0 in the binary representation. We implemented five types of constraints in our case study described in the main text (see Text S9). The currently implemented constraints are based on our current experience with using *boolmore* on a signal transduction network. Future applications will likely reveal new constraints, which can be implemented.

#### Allowing the addition of new edges from a limited pool of experimentally-supported hypotheses

*Boolmore* modifies the interaction graph by deleting or adding edges. The deletion of edges is done implicitly through the modification of regulatory functions. The addition of edges was implemented in a restrictive way: *boolmore* can only select edges from a predetermined pool. Initially, a newly added regulator is integrated with the rest in a random way, and the function can mutate through the iterations. For example, consider that node X, which had the original function $f_X$(A,B,C) = B&C | A&B | A&B&C, acquires a potential new positive regulator D. Its binary representation now has 16 digits instead of 8, thus acquiring 8 free parameters, which we mark with the symbol "?": $f_X$(A,B,C,D) = 0 | ?&D | 0&C | ?&C&D | 0&B | ?&B&D | 1&B&C | ?&B&C&D | 0&A | ?&A&D | 0&A&C | ?&A&C&D | 1&A&B | ?&A&B&D | 1&A&B&C | ?&B&C&D = 0?0?0?1?0?0?1?1?. If each "?" is set to zero, then the additional regulator is fully redundant, whereas the regulator is sufficient for activation if the first "?" is equal to one. When adding an additional regulator in *boolmore*, we initialize each "?" randomly, with a 50% chance to be 0 or 1.

Models are given internal penalties when adding edges, so that models in which the addition of an edge did not lead to a score increase are less likely to survive through the iterations. This is done by counting the number of prime implicants (or equivalently the number of 1s in the binary representation) and prioritizing the model with a smaller number whenever there are two models with the same agreement with the experiments. This method helps prevent the models from deviating too much from the original interaction graph and from increasing their complexity. However, even with these preventive measures, each edge in the pool makes the search space exponentially larger, and can preclude the algorithm from finding an optimal model in a reasonable amount of time. Hence we only allowed the addition of user-provided edges, often limited to edges with experimental support.

#### Interpreting experimental results in a Boolean context

*Boolmore* computes each model's fitness score using input data consisting of experimental interventions and a coarse-graining of the corresponding experimental outcomes. We used five categories of experimental results for each node: OFF, OFF/Some, Some, Some/ON, ON; we describe below how we assigned these categories, though we note that our workflow is flexible and allows other choices.

Experimental interventions such as knocking out a gene or providing excess amounts of a protein have natural Boolean interpretations; the corresponding nodes are considered to be fixed OFF for the former and ON for the latter. The observations of mRNA or protein concentrations in the unperturbed (wild type) or perturbed (e.g., mutant type) systems have a continuous spectrum of outcome. We use a comparative method to express these outcomes in a form that is compatible with Boolean dynamics. In the case of a signal transduction network, we use the levels of activation of the nodes in the presence/absence of the signal (in the wild type system) as two points of reference, akin to a positive and negative experimental control. We coarse-grain node activities in response to interventions by comparison to these two points of reference.

We illustrate in Figure S1A the case of a node R, which has higher activation when the signal S is present in the wild type. We consider this level of activation to be the ON state (R=1), and in any intervention that leads to a similar activation level or higher, R is considered ON. The levels are assigned to the OFF state in a similar manner. In addition to the preferred OFF and ON categories, we also introduced intermediate and mixed categories as necessary. If the activation level is an intermediate between the OFF and ON states, it is categorized as “Some”. Although a Boolean model has no simple way of describing such an intermediate level, it can be realized by an attractor in which the node oscillates or by multiple attractors, with the node being with ON in some of the attractors and OFF in the others. We discuss the interpretation of the “Some” category and its possible customization in Text S4. We assigned the OFF/Some or Some/ON categories in cases when there are multiple reported experiments for the same intervention that have non-identical results, or in cases where a clear comparison with the reference was impossible.

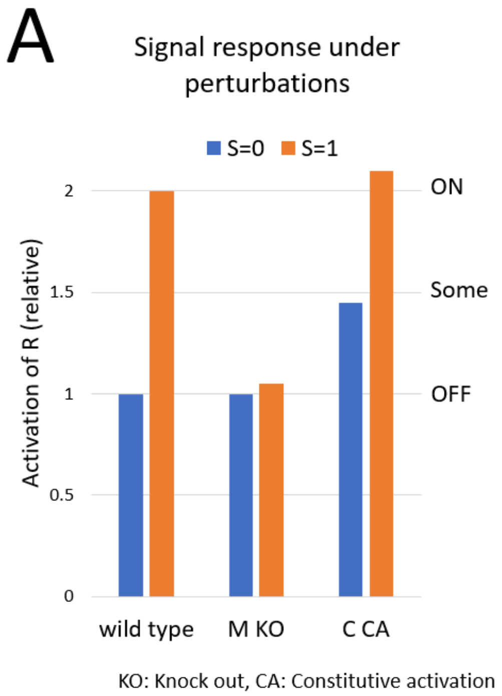


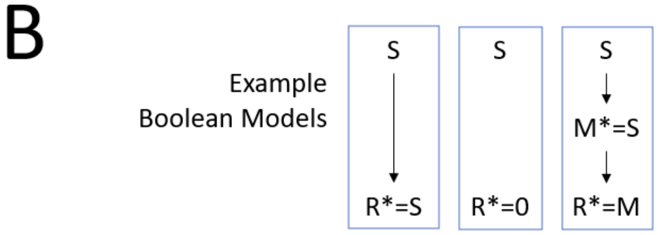


| | R*=S | R*=0 | R*=M |
|---|---|---|---|
| S=0 -> R=OFF | ✓ | ✓ | ✓ |
| S=1 -> R= ON | ✓ | ✗ | ✓ |
| S=1, M KO -> R=OFF | ✗ | ✓ | ✓ |
| Minimal trap space agreements | 2/3 | 2/3 | 3/3 |

| | R*=S | R*=0 | R*=M |
|---|---|---|---|
| S=0 -> R=OFF | ✓ | ✓ | ✓ |
| S=1 -> R= ON | ✓ | ✗ | ✓ |
| S=1, M KO -> R=OFF | ✗ | ✗ | ✓ |
| Hierarchy-based score | 2/3 | 1/3 | 3/3 |

Figure S1. (A) Illustrative example of perturbation experiments. The reported results are normalized by the activation of the response node R in the absence of the signal (S=0). The wild type shows higher activation of R in the presence of the signal S. M and C represent mediators of the signal.”KO” means knockout and “CA” means constitutive activation, thus “M KO” refers to M=0 and “C CA” refers to C=1. The wild type response under S=0 is categorized as OFF; the wild type response under S=1 is categorized as ON. If the value of R obtained under a perturbation is similar to one of these two reference values, it will be included in the same category as the reference. For example, the response to [S=1, C=1] is categorized as ON. Notably, the response to [S=0, C=1] is categorized as Some. (B) Illustrative example of determining the minimal trap space agreement and the hierarchy-based score of putative models. Each model indicates the next state of R (denoted R*) as a function of the current state of S or M. The model with R*=0 has the same minimal trap space agreement as the R*=S model, but it receives a lower hierarchy-based score because of its discrepancy with a wild-type experiment. In general, the difference between the minimal trap space agreement and the hierarchy-based score (highlighted in yellow) is more prominent in more complex perturbation experiments.

#### Detailed description of the model's outcome

We use *pyboolnet*, a Python package for analyzing Boolean models [13] to determine the minimal trap spaces of the model under various constraints that mimic experimental settings. Minimal trap spaces are a close, update-scheme-independent approximation of attractors and their identification is more computationally efficient (see Text S1). For each minimal trap space, nodes constrained to be ON are assigned the value of 1, nodes constrained to be OFF are assigned the value of 0, and unconstrained (oscillating) nodes are assigned the value of 0.5.

Although there are alternative tools for minimal trap space calculation that can outperform *pyboolnet* in typical settings, it is optimal for *boolmore*. This is because *pyboolnet* allows very fast computation of minimal trap spaces using the Blake canonical form of the functions, and its main bottleneck is calculating the Blake canonical form from the given functions. In *boolmore*, the binary representation allows the computation of the Blake canonical form at a very small cost.

In each comparison with the experimental result obtained in a perturbation condition, the model outcome is the average value of the observed node in the minimal trap spaces corresponding to that condition. For example, if there are two minimal trap spaces and the node oscillates in one and has the state 1 in the other, the average node value is 0.75.

#### Scoring model fitness with a hierarchy-based method

Each model receives one point toward its fitness score per recapitulated perturbation experiment outcome. To ensure that the simulated intervention is causally linked to the model outcome, a perturbation experiment is considered to be recapitulated only if the model predictions agree with measurements obtained for subsets of the perturbation as well, resulting in a hierarchy of experimental observations. The top of the hierarchy is the 'resting state' observation of the wild type system in the absence of any signal. The signal-response pairs of the unperturbed (wild type) system are one step down, as are the responses to perturbations of single intermediary nodes in the absence of any signal. The signal-response pairs under perturbations of single mediators are two steps from the top. Perturbations of multiple mediators are at increasingly lower levels of the hierarchy.

As an illustrative example, consider a system in which a signal S leads to a response R through a mediator M (see Figure S1B). The knockout of the mediator (M=0) inhibits the response to the signal (R=0 even if S=1). A hypothetical model that says R is inactive regardless of the signal (R*=0) recapitulates the result that M=0 leads to R=0. However, this model does not respond to the signal even when M=1. Our scoring method ensures that a high-scoring model satisfies the top-of-the-hierarchy experiment [S=0]→R=0, the one step down experiment [S=1]→R=1, as well as the more complex experiment [S=1, M=0]→R=0. Note that S=0 is the default and should be included in the specification of the experiment unless S=1.

*Boolmore* scores each model in a two-step process. First it determines the agreement of the model with each perturbation experiment, and then it scores the model by considering all agreements with the experiments at higher levels of the hierarchy. For each perturbation condition, the model prediction is the average value of the observed node in the minimal trap spaces. The model's agreement for that perturbation experiment is given depending on how well the prediction agrees with the categorization of the experimental outcome. Each agreement function's output ranges from 0 to 1 (see Table S1). For example, if the experimental outcome is categorized as ON, a model with a prediction of 1 receives an agreement of 1 for that experiment and another model with a prediction of 0.5 gets an agreement of 0.5. The final score is the product of all the attractor agreements of the subset perturbation experiments. For example, in Figure S1B, if we are considering the perturbation [S=1, M=0], the score is given by multiplying the agreements of 4 experiments, namely [S=0], [S=1], [S=0, M=0], and [S=1, M=0] itself. Going back to our previous example, the model with R*=0 has an agreement of 1 for the experiment [S=1, M=0]→R=0. However, as its agreement for [S=1]→R=1 is 0, it will not receive a score from this experiment.

**Details of running the genetic algorithm**

The steps described in Figure 1 of the main text are performed on a population of 100 models generated in each iteration. We do 100 iterations in each run. The top 20 models with highest fitness scores from the previous iteration are carried over. Twenty pairs selected randomly (fitter models having a higher chance of being selected) from this top 20 are mixed pairwise to produce 20 new models. Finally, from this pool of 40 models, we repeat 80 times the process of selecting a model and mutating it, to generate the remaining 80 models of the new iteration. We used a mutation probability of 0.01 in the functions. Fitter models have a higher chance of being selected in this process as well. Ten thousand networks are generated in a single run. These numbers were chosen such that the best score saturates by the end of the run in the benchmarks and the case study. We performed a parameter analysis and found significant robustness to changes in these parameter values (see Text S5 for more details).

### Text S4. Details of the interpretation of the “Some” category of experimental results

We assigned the “Some” category to experimental observations that show an intermediate node activation level between the OFF state (which is determined based on the negative control illustrated in Text S3 and Figure S1) and the ON state (which is determined by the positive control). In our implementation, the model is considered to agree with the “Some” category if the average value of the node in the minimal trap spaces is near 0.5. This can be realized by an attractor in which the node oscillates, by multiple attractors with the node being with ON in some of the attractors and OFF in the other, or by a combination of oscillating and point attractors. However, there are multiple biological mechanisms that can yield such an intermediate activation level, and our scoring method can be customized to reflect them if they are known.

Most observations on the activation of cellular elements are population-averaged, which also yields an implicit time-averaging when the cells in the population are not synchronized. Hence, the biology underlying an observed intermediate activation level can belong to one of 3 broad categories. First, it could happen that the element’s activation level oscillates in each cell, but the population-average shows an intermediate value. An example of such oscillation in our case study is the transient spikes of cytosolic calcium level in stomatal guard cells. Such oscillation can be verified when time-course observations of single cells are available [62]. Second, the population can be bimodal, with the element being ON in some cells and being OFF in the others. This behavior can be verified when the distribution of the element’s activation level in individual cells is available. Last, it could happen that the element’s activation level is indeed intermediate in every individual cell over time.

If there is evidence for oscillation or bimodal distribution, our workflow can be customized to give a score only when the generated models reflect the actual dynamics. In our case study this was unnecessary as the calcium oscillation was guaranteed by the network structure. A Boolean model has no simple way of reflecting the last case, when the activation level is actually intermediate. One possibility, used in previous modeling [63], is to interpret oscillations of the Boolean model as excursions (repeated increases and decreases) around a dynamic equilibrium, and equate the fraction of the ON state with the observed intermediate level. The other possibility is to introduce an intermediate activation level, either by making the model multi-level, or by introducing a new node, which represents the intermediate activation level, into the Boolean model.

### Text S5. Parameter analysis

*Boolmore* contains multiple tunable parameters: the mutation probability, the number of iterations, the number of models generated in each step, the number of top-scoring models kept at each step, and the number of models generated by crossing existing models. We explored multiple values of each parameter for a compilation of 8 models from the Cell collective. These 8 models were selected to exemplify a variety of properties: small (<20) or large (>70) number of nodes, presence or absence of source nodes, single or multiple point attractors, single or multiple complex attractors. We used each model to generate 5 sets of 10*N artificial experiments. We generated five starting models for each model by randomizing the binary representation of the update functions of the original model. Here we illustrate the optimization of the mutation probability and present the general conclusions of our parameter analysis. See the github repository for results on other parameters.

#### Analysis of mutation probability

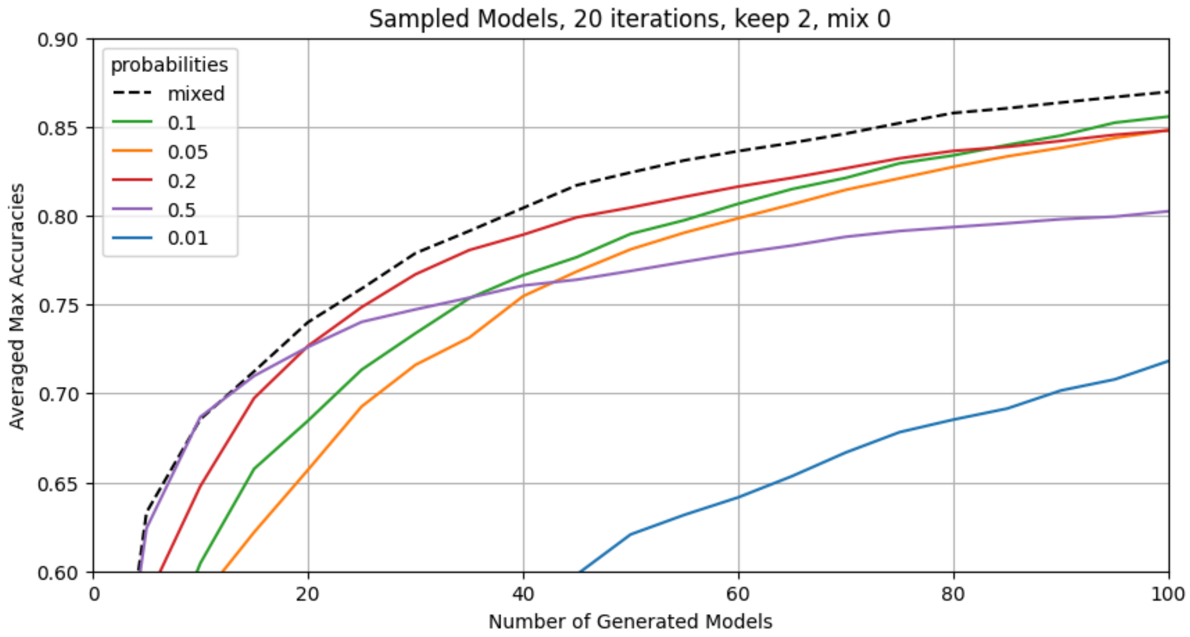

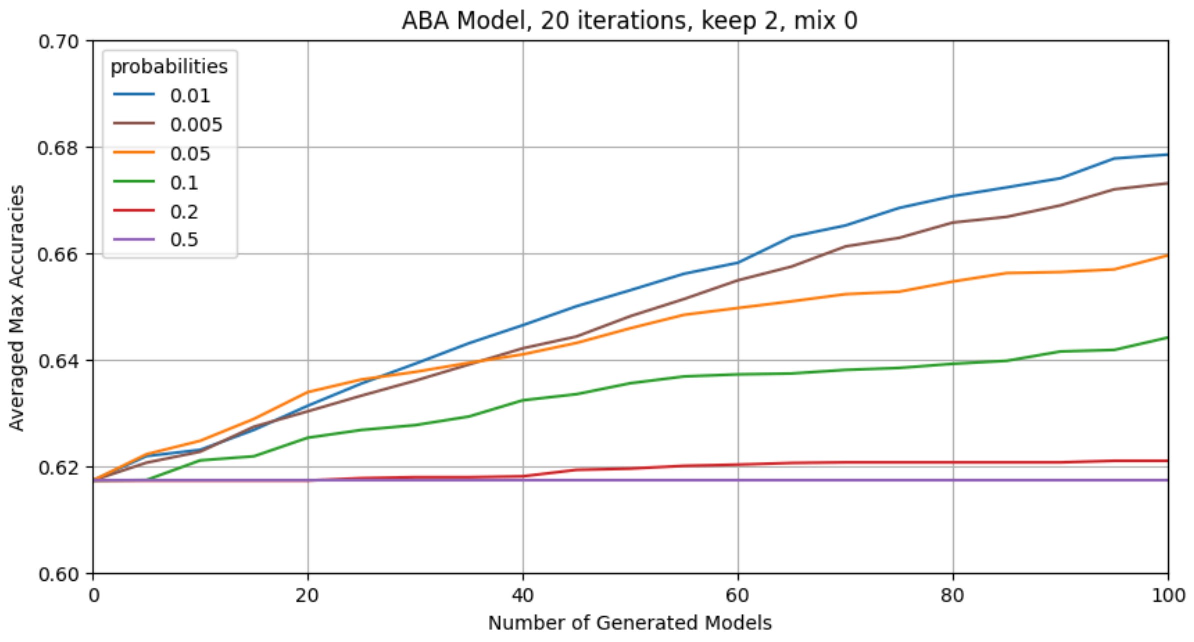


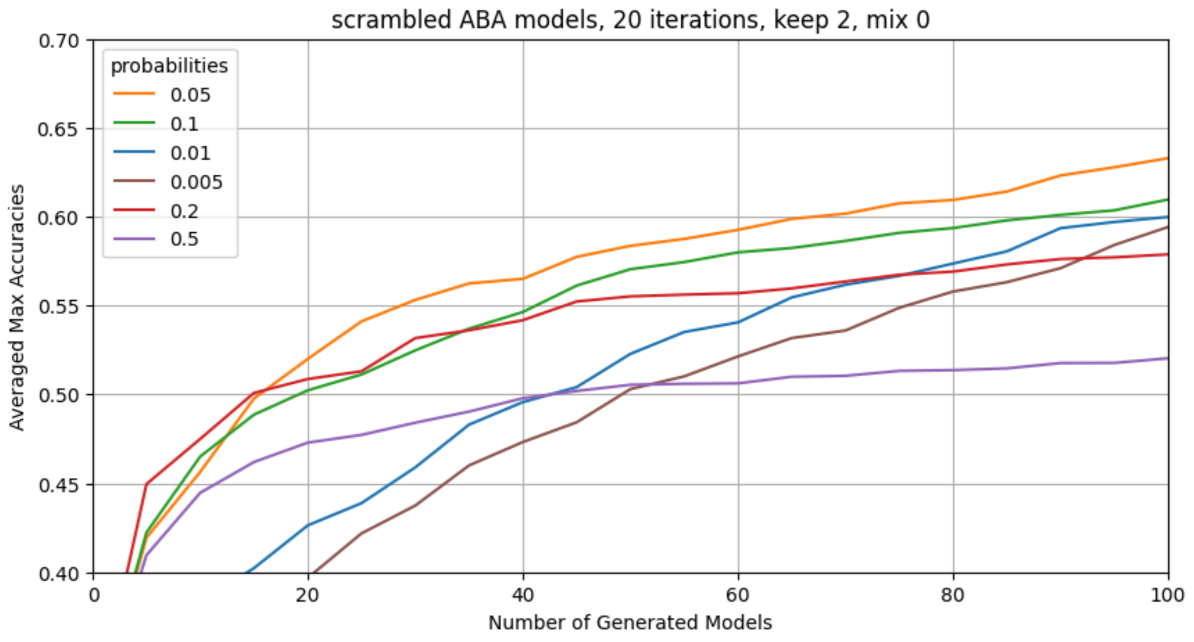


The mutation probability denotes the probability of changing (flipping) each bit of the binary representation of each regulatory function. For every run we generated 100 models over 20 iterations, keeping 2 models with the highest scores and no mixing. For the sampled models, we tested 5 probability values, from 0.01 to 0.5. We observed that the probability value of 0.5 yielded the highest accuracy in the first 4 iterations (20 models generated), but then it was surpassed by the models with probability of 0.2, and at iteration 17 also by the models with probability of 0.1. Based on this result, we designed an adaptive probability setting with a value of 0.5 for the first 3 iterations and a value of 0.1 afterward. This probability setting yielded the best accuracy (see top panel of Figure S2).

We also varied the mutation probability used for the refinement of the ABA model. As the starting model is much fitter than a randomized model, it is expected that a lower mutation probability would yield better accuracy. Accordingly, we tested 6 probability values, from 0.005 to 0.5 with 50 independent runs. We found that the probability of 0.1 led to the highest accuracy, followed by the probability of 0.005 (see middle panel of Figure S2). The mutation probability of 0.5 did not achieve any improvement over the original model. If instead of the published model we started with randomized update functions (that nevertheless satisfy the biological constraints described in Text S9), the probability that yielded the best accuracy was 0.05, and both high and low mutation probabilities yielded poor accuracy (see bottom panel of Figure S2).

We conclude that high mutation probabilities are an appropriate choice for starting models with a low fitness and low mutation probabilities are an appropriate choice for higher-fitness starting models. Probability settings with iteration-dependent probabilities (e.g. high mutation probability for the initial exploration, followed by low mutation probability for refinement) may be the best choice in certain use cases.

#### Conclusions

We found that the performance of the genetic algorithm does not depend sensitively on the choice of the parameters. The most important choice is the selection of the mutation probabilities, as the most appropriate mutation rate depends on the fitness of the starting model. Another consideration is that in general, larger models require lower mutation probabilities to allow fine-tuning of the well-performing models. However, any mutation probability in the range of 0.01-0.1 can sufficiently refine the model and reach saturation with enough iterations. We found that the other parameters have negligible impact on the performance. When an equal number of models were generated, the number of iterations did not make a significant difference as long as it was over a certain threshold, i.e. larger than 10 when 100 models are generated. Similarly, the number of models kept to the next iteration did not make a significant difference as long as it was comparable to the number of models generated for each iteration, i.e. lower than 5 when 5 models are generated at each iteration. The optimal number of models to generate using mix (crossing) is small, i.e., 1 for the sampled models when 5 models were generated for each iteration.

### Text S6. The details of the benchmark analysis to test the overall performance of *boolmore*

Following the common practice in such benchmarks, we used existing Boolean models to generate artificial experiments. We used 80% of the artificial experiments as the training set to refine a randomized starting model generated using the original model's interaction graph; we used the remaining 20% as the validation set to test the accuracy of the refined model in recapitulating newly encountered experimental results.

We used 40 Boolean models in the Cell Collective (a repository of peer-reviewed, experimentally supported Boolean models [37]) and ran 5 replicate benchmark runs for each of them. We selected models with fewer than 30 nodes. For each model with N nodes, we generated five sets of 10*N artificial experiments, 80% of which were used for training, for a coverage that is comparable to that of our case study (505 experiments for 68 not fully constrained nodes). Each artificial experiment consists of a set of nodes whose state is controlled (kept fixed) and a node whose state is observed. The controlled set of nodes always included the source nodes of the network; each non-source node had a 1/N chance of also being included. The observed node was one of the sink nodes or, with a lower priority, it was a randomly selected node. We fixed the values of the nodes in the controlled set randomly and determined the average node value of the observed node in the minimal trap spaces. Depending on the average value, the result was classified into one of the 5 categories described previously, using thresholds that ensure that the original model would have a perfect fitness score.

We generated five starting models for each model by randomizing the binary representation of the update functions of the original model. This randomization keeps the functions monotonic and consistent with the original interaction graph, but may yield fewer regulators than the original. The missing regulators can be added back in during the iterations of the algorithm. We used *boolmore* to refine the models over 100 iterations and score them by comparing their results to the results of a subset (80%) of the artificial experiments following the procedure described in the Methods.

### Text S7. Detailed description of abscisic acid-induced closure and its Boolean modeling

Stomata are microscopic pores on the leaves through which the plants take in $CO_2$ necessary for photosynthesis and through which the plant inevitably loses water. The stomata open in response to environmental signals such as light. When the plant is dehydrated, it produces a stress hormone called abscisic acid (ABA) whose role is to help the plant conserve water. ABA acts as a signal in a network of protein interactions, enzyme-catalyzed reactions and ion flows, whose ultimate effect is the relaxation of plant guard cells and subsequent closure of stomata. The mechanisms involved in this network include activation of enzymes, production of molecules such as reactive oxygen species (ROS), nitric oxide (NO), and phosphatidic acid (PA), transient increases in the cytosolic $Ca^{2+}$ level ($Ca^{2+}{}_c$) due to $Ca^{2+}$ flow through the membrane (CaIM) or to $Ca^{2+}$ release from stores (CIS), and ion flow across the cellular membrane.

Understanding this network provides an avenue for gaining insight about the regulatory mechanisms underlying drought tolerance, which is of paramount importance for plant survival and crop resilience. The variety of processes involved in this network also exemplifies the potential complexity of the signal processing of living systems. The first Boolean model of ABA-induced closure was developed starting in 2003 and was published in 2006 [28]. At that time, the receptor proteins to which ABA binds were not known. Following the identification and confirmation of the RCAR receptor family [64–66], the ABA-induced closure model was significantly expanded and an updated version was published in 2017 [29]. The 2017 version of the model was subsequently improved in 2019 to better account for the possibility of stomatal closure in response to intracellular stimuli [30] and then in 2020 to reproduce the reversibility of stomatal closure [31].

We use the 2017 model as the basis of model refinement, as it is the most comprehensive and its analysis included a thorough comparison with experiments. The full and abbreviated names of all the elements included in the 2017 model are listed in S3 Table. One notable feature of the model is that it includes a negative feedback loop between the node $Ca^{2+}{}_c$, which represents an increase in the cytosolic level of $Ca^{2+}$ , and the node $Ca^{2+}$ATPase, which represents the pumps and channels that decrease the cytosolic $Ca^{2+}$ level to avoid damage to the cell. This negative feedback loop leads to an irregular $Ca^{2+}{}_c$ oscillation in the model in the presence of ABA, which represents the transient $Ca^{2+}{}_c$ increases of the real system [62]. The attractor of the model in the presence of ABA features the sustained state of 71 nodes (including the outcome node Closure) and the oscillating state of 10 nodes (including $Ca^{2+}{}_c$ and $Ca^{2+}$ATPase). Albert et al. used simulations to classify perturbations (e.g., genetic or pharmacological knockouts, constitutive activation of elements) into five categories in terms of the closure response to ABA: close to wild type response, hypersensitivity (faster closure), hyposensitivity (slower closure), reduced closure, and insensitivity to ABA (i.e., lack of closure). When comparing model results and experimental results, any defects in closure were considered consistent with each other. Based on a comparison with 112 perturbation experiments, the reported accuracy of the 2017 model was 95/112=85%.

Despite the overall high accuracy, a significant weakness of the 2017 model is that it does not recapitulate the reversibility of stomatal closure. This is because the model has an attractor with Closure=1 out of 17 attractors in the absence of ABA. After the model converges to the closure attractor in response to ABA, the removal of ABA will lead to this single attractor, and not cause loss of closure.

Two follow-up publications aimed to address each of these two weaknesses of the 2017 model by making parsimonious hypotheses. Maheshwari et al. [30] hypothesized in 2019 the existence of an additional edge. They found that the model augmented by an inhibitory edge starting from cytosolic $Ca^{2+}$ and targeting any of four PP2C phosphatase family members recapitulates five of the nine experimental observations of closure that were not captured by the 2017 model. They confirmed this inhibitory effect by experimentally observing a decrease in PP2C phosphatase activity in response to externally provided $Ca^{2+}$. Considering the possibility of this effect being indirect, Maheshwari et al. also demonstrated that it could be mediated by PA with very similar model results.

The 2019 model [30] shared the irreversibility of stomatal closure exhibited by the 2017 model. A second follow-up to the model in 2020 [31] identified that the source of the irreversibility of stomatal closure in the model is the assumption of self-sustained activity of four nodes regulated by $Ca^{2+}_c$. This assumption was made in the 2017 model because if the activity of these four nodes would decrease when $Ca^{2+}_c$ decreases, then sustained closure would not be possible. The 2020 model incorporated the additional edge introduced in the 2019 model and made the regulatory function of these four nodes time-dependent. This time dependence serves as a memory of past $Ca^{2+}_c$ increase that makes the activity of these nodes less likely to decrease, making the assumption of their self-regulation unnecessary. The 2020 model achieved reversible closure and preserved the 2019 model's success in capturing five of the experimental observations of closure in response to interventions.

### Text S8. Correcting an error in the 2017 model and choosing baseline models

The original 2017 model had an erroneous edge between the node Vacuolar Acidification and the node KEV, which represents vacuolar $K^+$ channels necessary for the ion flow that precedes stomatal closure; the inaccuracy of this edge was recognized in a later publication [67]. We found that the model corrected by deleting this edge had a low accuracy (see model "VA-KEV del" in Text S12), and was unable to recapitulate sustained stomatal closure in response to ABA. The reason for this failure is that the sole remaining regulator of KEV, $Ca^{2+}_c$, oscillates, leading to oscillating closure. We attempted to fix this discrepancy using *boolmore*. Although the algorithm achieved a score improvement (see model "GA - VA-KEV del" in Text S12), it was unable to achieve sustained activation of closure in response to ABA. This shows the importance of having a good starting interaction graph for our workflow. To have a starting model that recapitulates the core behavior of ABA-induced closure, we pursued two ways to eliminate the spurious oscillations in the output node Closure.

First, drawing inspiration from the original model, which assumed self-sustained activity of four nodes that would otherwise exhibit $Ca^{2+}_c$-driven oscillations, we sought to identify a node between KEV and the output node Closure for which a self-sustaining influence would be biologically justified and able to yield sustained closure. After considering multiple nodes, we chose the abstract process node “$H_2O$ efflux”, whose inputs include AnionEM, $K^+$ efflux, Malate, and Aquaporin. Indeed, sustained water efflux from the guard cells through the aquaporin channels is a hallmark of stomatal closure. The improved regulatory function contains a self-edge on the node “$H_2O$ efflux”, and expresses the assumption that water efflux, once started, can be sustained even if $K^+$ efflux stops, if all other conditions continue to be met. This model recapitulated ABA-induced closure and had a score of 311.7/505 (61.7%). We adopted this model as the first baseline model (A).

Second, in an alternative approach, we introduced a global, qualitative change to the 2017 model by implementing a different representation of the dynamics of cytosolic $Ca^{2+}$. The literature on ABA-induced stomatal closure includes significant debate regarding the role of cytosolic $Ca^{2+}$ [62,68]. Substantial spatio-temporal variability was observed, including spatial hotspots, transients, spikes, and oscillations [62]. The emerging conclusion is that cytosolic $Ca^{2+}$ elevation plays an amplifying role in ABA signaling [69]. Nevertheless, there is no consensus on whether the return from the elevated $Ca^{2+}$ level has a regulatory role (in other words, whether it matters if $Ca^{2+}$ has one or multiple increases). In the original and the baseline A model, the oscillation of $Ca^{2+}_c$ drives 10 nodes into an oscillating state. In contrast, the existing experimental observations of 32 elements in response to ABA only indicate an oscillatory activity for $Ca^{2+}_c$. We note that many assays observe a population of guard cells at a single time point; such assays can neither rule out nor support oscillations of certain elements at the level of individual guard cells. As an alternative to the self-sustained activation of $Ca^{2+}_c$ targets assumed in the 2017 model and in the baseline A model, we replaced the two-node negative feedback loop with a single node, ‘$Ca^{2+}_c$ osc’, whose ON state represents the appearance of $Ca^{2+}$ elevation transients and whose OFF state represents an unchanged low cytosolic $Ca^{2+}$ level. We also deleted the four assumed self-edges. The replacement of an oscillating node with a non-oscillating one is a potentially dramatic change because it stabilizes the positive regulation and positive feedback loops in the system. Indeed, this model only had point attractors. We found that the thus-modified model had a score of only 184.2/505 (36.5%), mainly because it did not recapitulate the reduction of ABA-induced closure for single-node interventions. We adopted this model as our second baseline (B), aiming to find out whether a model that lacks oscillations can reach equal agreement with the experimental observations.

### Text S9. Details of the constraints and extra edges used for improving the ABA-induced closure model

#### Constraints on the regulatory functions

We use four types of constraints: fixed functions, edge preservation, logic preservation, and grouped regulation.

We apply the fixed function constraint to 13 nodes, which prevents their update functions from mutating. These update functions of the 2017 model express known biochemical reactions or biophysical mechanisms with especially firm and direct evidence. One such example of a fixed function is cADPR* = $NAD^+$ and ADPRc. This function describes the sole reaction in which cADPR (cyclic ADP ribose) is produced in this system; the reactant of this reaction is $NAD^+$ and it is catalyzed by ADPRc (ADP-ribosyl cyclase).

Apart from fixing these 13 update functions, we also enforce that 9 additional edges are preserved. These are 9 cases of regulator-target pairs that have direct experimental evidence but insufficient evidence to constrain how the regulator is coupled to the other regulators of the target. The edge preservation constraint ensures that the update function of the target node always contains the regulator as a non-redundant input. One such example is the AnionEM→Depolarization edge, which indicates that the efflux of anions promotes the depolarization of the plasma membrane. Every version of the regulatory function of the node Depolarization must contain AnionEM. We also prevent each node from becoming a source node due to edge loss. The exception is GEF1/4/10, which is a source node in the original model but can acquire (and later lose) a new edge from the pool.

We preserve the logical role of 6 additional edges. For these regulator-target pairs, it is known that the activity of the regulator is necessary for the activation of the target. We require that any modification of the function of the target node preserves this necessity. One such example is the necessity of the enzyme activity of RBOH (respiratory burst oxidase homolog) to produce reactive oxygen species (ROS node). A regulatory function of the type ROS* = RBOH and X, where X is an additional regulator of ROS, reflects the necessity of RBOH for the production of ROS.

Finally, we impose a grouped regulation constraint to four groups of nodes. In these cases, it is known that a specific enzyme-catalyzed reaction may contribute to the production of a molecule, but additional possible reactions or regulatory effects allow flexibility to the function. For example, three separate reactions were identified as potentially contributing to the production of PA during stomatal closure, but the relative contributions of these reactions are not known. The 2017 model assumed that each of the three reactions is independently sufficient to produce a sufficient amount of PA that qualifies as its ON state. We allow modifications from this assumption, but we require the enzyme and substrate of each reaction to appear as a group in the regulatory function of PA. Below we summarize the full list of constraints.

#### List of constraints

1. fixed functions: $Ca^{2+}$ ATPase, $Ca^{2+}_c$ ($Ca^{2+}_c$ osc), Closure, DAG, $H_2O$ efflux, InsP3, InsP6, NO, PtdIns(3,5)P2, PtdIns(4,5)P2, RCARs, cADPR, cGMP
2. edge preservation: RCARs ⊣ ABI1, RCARs ⊣ ABI2, RCARs ⊣ HAB1, RCARs ⊣ PP2CA, KEV→$K^+$ efflux, KOUT→$K^+$ efflux, ABI1 ⊣ OST1, ABI2 ⊣ OST1, AnionEM→Depolarization
3. preservation of "necessary" logic: cGMP→8-nitro-cGMP, Depolarization→KOUT, PEPC→Malate, AnionEM ⊣ Malate, NADPH→ROS, RBOH→ROS
4. grouped regulation: (PC, PLDα),(PC, PLDδ),(DAG, DAGK)→PA, (SPHK1/2, Sph)→S1P/PhytoS1P
5. possible source: GEF1/4/10

#### The pool of additional edges

We used a set of 13 experimentally documented interactions among the existing nodes of the network as a pool, from which the algorithm can select edges to add to the model.

1. Ca2c ⊣ ABI2, HAB1, PP2CA, PA ⊣ ABI2, HAB1, PP2CA
These edges originate from the experimental observation that $Ca^{2+}$ inhibits the phosphatase activity of PP2C family members [30]. Model analysis published together with this observation supports that this effect could be mediated by PA.
Indeed, according to [70] PA binds to and inhibits PP2CA; 40% of the phosphatase activity of PP2CA is lost for 100 μM PA.

2. Aquaporin(PIP2;1) → ROS
As described in [71], RBOHs produce extracellular superoxide by transferring electrons from NADPH or FADH2 to oxygen. This extracellular $H_2O_2$ enters the plant cells through plasma membrane aquaporins. Reference [72] shows that knocking out Aquaporin eliminated ROS production under ABA.

3. Actin Reorganization → RBOH
Reference [73] showed that alteration of actin reorganization delays ABA-evoked $H_2O_2$ generation in guard cells. This effect of actin reorganization can explain a previously observed indirect effect of PtdInsP3 and PtdInsP4 (both regulators of actin reorganization) on ABA-induced ROS production [74,75].

4. ROS → Actin Reorganization
Reference [73] found that actin reorganization was impaired in *atrbohd/f* mutants (which lack RBOH), and that exogenous $H_2O_2$ application to *atrbohd/f* mutant leaves induced actin reorganization.

5. PA → Microtubule Depolymerization
In the 2017 model Microtubule Depolymerization is activated by TCTP, which in turn is regulated by $Ca^{2+}_c$. Figure 4f of [76] indicates that PA leads to microtubule depolymerization even in the absence of $Ca^{2+}_c$, suggesting the existence of a $Ca^{2+}$-independent mechanism.

6. $pH_c$ → Vacuolar Acidification
Reference [77] suggests a positive feedback loop in which vacuolar acidification positively affects $pH_c$ (and is necessary for it) and $pH_c$ positively affects vacuolar acidification (and is necessary for it). The first part is incorporated in the 2017 model whereas the second part is not in the model.

7. ABI1 → GEF1/4/10
GEF1/4/10, considered a source node in the 2017 model, was found to be regulated by ABI1 [69]. ABA leads to the degradation of GEF1, likely promoted by CPKs [78]. ABI1 protects GEF1 from degradation.

8. GHR1 → CPK3/21
GHR1 is found to interact with CPK3 [79], probably acting as a scaffold to bring CPK3 to SLAC1.

### Text S10. The Boolean functions of the 2017 model and of the two GA-improved models

In these functions the state variable of each node is represented by the name of the node, and the future state of each node is denoted by using an asterisk. For example C* = A or B means that the state of node C will be updated to 1 when A=1 or B=1. The functions reflect the notation used in the code of our algorithm; certain node names are written slightly differently in the code compared to the manuscript to avoid running errors. The full name of each node can be found in Table S3. The logical operator & indicates AND, | indicates OR and ! indicates NOT.

Boolean functions of the 2017 model:
8-nitro-cGMP* = NO & ROS & cGMP
ABA* = ABA
ABH1* = 1
ABI1* = !PA & ROP11 & !ROS & pHc | !PA & !RCARs & !ROS & pHc
ABI2* = ROP11 & !ROS | !RCARs & !ROS
ADPRc* = 8-nitro-cGMP
ARP_Complex* = 1
Actin_Reorganization* = ARP_Complex & !AtRAC1 & PtdInsP4 & SCAB1 | ARP_Complex & !AtRAC1 & PtdInsP3 & SCAB1
AnionEM* = QUAC1 & SLAH3 | SLAC1
AquaporinPIP2_1* = OST1
AtRAC1* = ABI1 | !ABA
CIS* = cADPR | InsP6 | InsP3
CPK23* = 1
CPK3_21* = Ca2c | CPK3_21
CPK6* = 1
Ca2_ATPase* = Ca2c
Ca2c* = !Ca2_ATPase & CaIM | CIS & !Ca2_ATPase
CaIM* = GHR1 & MRP5 & NtSyp121 | SACC | !ERA1 | !ABH1
Closure* = H2O_Efflux & Microtubule_Depolymerization
DAG* = PLC & PtdIns4_5P2
DAGK* = 1
Depolarization* = KEV & !K_efflux | !H_ATPase & KEV | Ca2c & !K_efflux | Ca2c & !H_ATPase | AnionEM & !K_efflux | AnionEM & !H_ATPase
ERA1* = 1
GAPC1_2* = 1
GCR1* = 1
GEF1_4_10* = 0
GHR1* = !ABI2 & ROS
GPA1* = S1P_PhytoS1P | !GCR1
GTP* = 1
H2O_Efflux* = AnionEM & AquaporinPIP2_1 & K_efflux & !Malate
HAB1* = !RCARs & !ROS
H_ATPase* = !Ca2c & !ROS & !pHc
InsP3* = PLC & PtdIns4_5P2
InsP6* = InsP3
KEV* = Vacuolar_Acidification | Ca2c
KOUT* = Depolarization & pHc | Depolarization & !ROS | Depolarization & !NO
K_efflux* = KEV & KOUT
MPK9_12* = MPK9_12 | Ca2c

MRP5* = 1
Malate* = !ABA & !AnionEM & PEPC
Microtubule_Depolymerization* = TCTP | Microtubule_Depolymerization
NAD* = 1
NADPH* = 1
NIA1_2* = ROS
NO* = NADPH & NIA1_2 & Nitrite
NOGC1* = NO
Nitrite* = 1
NtSyp121* = 1
OST1* = !HAB1 & !PP2CA | !ABI2 & !PP2CA | !ABI2 & !HAB1 | !ABI1 & !PP2CA | !ABI1 & !HAB1 | !ABI1 & !ABI2
PA* = PC & PLDdelta | PC & PLDalpha | DAG & DAGK
PC* = 1
PEPC* = !ABA
PI3P5K* = ABA
PLC* = Ca2c
PLDalpha* = Ca2c & GPA1
PLDdelta* = GAPC1_2 & ROS | NO
PP2CA* = !RCARs & !ROS
PtdIns3_5P2* = PI3P5K
PtdIns4_5P2* = PtdInsP4
PtdInsP3* = 1
PtdInsP4* = 1
QUAC1* = Ca2c & OST1
RBOH* = !ABI1 & GPA1 & OST1 & PA & PtdInsP3 & RCN1 & pHc
RCARs* = ABA
RCN1* = 1
ROP11* = GEF1_4_10
ROS* = NADPH & RBOH
S1P_PhytoS1P* = SPHK1_2 & !SPP1 & Sph
SACC* = Actin_Reorganization
SCAB1* = 1
SLAC1* = !ABI1 & !ABI2 & CPK6 & GHR1 & MPK9_12 & OST1 & !PP2CA & pHc | !ABI1 & !ABI2 & CPK3_21 & GHR1 & MPK9_12 & OST1 & !PP2CA & pHc | !ABI1 & !ABI2 & CPK23 & GHR1 & MPK9_12 & OST1 & !PP2CA & pHc
SLAH3* = !ABI1 & CPK3_21 & CPK6 | !ABI1 & CPK23 & CPK3_21
SPHK1_2* = PA | ABA
SPP1* = 0
Sph* = 1
TCTP* = Ca2c
V-ATPase* = Ca2c
V-PPase* = PtdIns3_5P2
Vacuolar_Acidification* = Vacuolar_Acidification | V-PPase | V-ATPase
cADPR* = ADPRc & NAD
cGMP* = GTP & NOGC1
pHc* = !ABI1 & !ABI2 & OST1 & Vacuolar_Acidification | Ca2c & Vacuolar_Acidification

Boolean function of the GA1-A model:

8-nitro-cGMP* = NO & cGMP
ABA* = ABA
ABH1* = 1

ABI1* = !PA & !RCARs | !PA & !ROS & pHc | ROP11 & pHc
ABI2* = !RCARs & ROP11 | !PA
ADPRc* = 8-nitro-cGMP
ARP_Complex* = 1
Actin_Reorganization* = ARP_Complex & !AtRAC1 & PtdInsP3 & PtdInsP4 | ARP_Complex & !AtRAC1 & PtdInsP3 & ROS | PtdInsP3 & PtdInsP4 & ROS
AnionEM* = QUAC1 & SLAC1
AquaporinPIP2_1* = OST1
AtRAC1* = ABI1 | !ABA
CIS* = InsP3 | cADPR
CPK23* = 1
CPK3_21* = CPK3_21 & Ca2c
CPK6* = 1
Ca2_ATPase* = Ca2c
Ca2c* = !Ca2_ATPase & CaIM | CIS & !Ca2_ATPase
CaIM* = !ABH1 & SACC | GHR1 | MRP5 & NtSyp121 & SACC
Closure* = H2O_Efflux & Microtubule_Depolymerization
DAG* = PLC & PtdIns4_5P2
DAGK* = 1
Depolarization* = AnionEM & !H_ATPase | Ca2c & !H_ATPase | !H_ATPase & KEV
ERA1* = 1
GAPC1_2* = 1
GCR1* = 1
GEF1_4_10* = 0
GHR1* = !ABI2
GPA1* = S1P_PhytoS1P | !GCR1
GTP* = 1
H2O_Efflux* = AnionEM & AquaporinPIP2_1 & K_efflux & !Malate | AnionEM & AquaporinPIP2_1 & H2O_Efflux & !Malate
HAB1* = !RCARs & !PA | !ROS & !PA
H_ATPase* = !pHc
InsP3* = PLC & PtdIns4_5P2
InsP6* = InsP3
KEV* = Ca2c
KOUT* = Depolarization
K_efflux* = KEV & KOUT
MPK9_12* = Ca2c
MRP5* = 1
Malate* = !ABA & !AnionEM & PEPC
Microtubule_Depolymerization* = PA
NAD* = 1
NADPH* = 1
NIA1_2* = ROS
NO* = NADPH & NIA1_2 & Nitrite
NOGC1* = NO
Nitrite* = 1
NtSyp121* = 1
OST1* = !ABI1 & !ABI2 | !ABI1 & !HAB1
PA* = DAG & DAGK & PC & PLDalpha | PC & PLDdelta
PC* = 1
PEPC* = !ABA
PI3P5K* = ABA

PLC* = Ca2c
PLDalpha* = Ca2c
PLDdelta* = GAPC1_2 & NO & ROS
PP2CA* = !RCARs
PtdIns3_5P2* = PI3P5K
PtdIns4_5P2* = PtdInsP4
PtdInsP3* = 1
PtdInsP4* = 1
QUAC1* = OST1
RBOH* = GPA1 & OST1 & PA & PtdInsP3 & RCN1 & pHc
RCARs* = ABA
RCN1* = 1
ROP11* = GEF1_4_10
ROS* = NADPH & RBOH & AquaporinPIP2_1
S1P_PhytoS1P* = SPHK1_2 & Sph
SACC* = Actin_Reorganization
SCAB1* = 1
SLAC1* = !ABI1 & !ABI2 & CPK23 & CPK6 & GHR1 & MPK9_12 | !ABI1 & !ABI2 & CPK23 & GHR1 & MPK9_12 & OST1 | !ABI1 & !ABI2 & CPK6 & GHR1 & OST1 | !ABI2 & CPK23 & CPK3_21 & GHR1 & OST1 & pHc | !ABI2 & CPK23 & GHR1 & MPK9_12 & pHc | !ABI2 & CPK6 & GHR1 & OST1 & !PP2CA & pHc
SLAH3* = CPK23 & CPK3_21 | CPK3_21 & CPK6
SPHK1_2* = PA | ABA
SPP1* = 0
Sph* = 1
TCTP* = Ca2c
V-ATPase* = Ca2c
V-PPase* = PtdIns3_5P2
Vacuolar_Acidification* = V-ATPase | V-PPase & pHc
cADPR* = ADPRc & NAD
cGMP* = GTP & NOGC1
pHc* = !ABI1 & OST1 & Vacuolar_Acidification | !ABI2 & Ca2c | Ca2c & OST1

Boolean functions of the GA1-B model:
8-nitro-cGMP* = NO & cGMP
ABA* = ABA
ABH1* = 1
ABI1* = !PA & !ROS | !RCARs & ROP11 & !ROS & pHc
ABI2* = !RCARs & !Ca2osc & !PA | ROP11 & !Ca2osc | !ROS & !Ca2osc
ADPRc* = 8-nitro-cGMP
ARP_Complex* = 1
Actin_Reorganization* = ARP_Complex & !AtRAC1 & PtdInsP3 & PtdInsP4 & ROS
AnionEM* = QUAC1 & SLAC1
AquaporinPIP2_1* = OST1
AtRAC1* = ABI1 | !ABA
CIS* = InsP6 & cADPR
CPK23* = 1
CPK3_21* = Ca2osc
CPK6* = 1
Ca2osc* = CaIM | CIS
CaIM* = !ABH1 & !ERA1 & GHR1 & MRP5 | !ABH1 & MRP5 & SACC | !ERA1 & GHR1 & NtSyp121 | !ERA1 & MRP5 & SACC | GHR1 & MRP5 & NtSyp121

Closure* = H2O_Efflux & Microtubule_Depolymerization
DAG* = PLC & PtdIns4_5P2
DAGK* = 1
Depolarization* = AnionEM & !H_ATPase | AnionEM & !K_efflux
ERA1* = 1
GAPC1_2* = 1
GCR1* = 1
GEF1_4_10* = 0
GHR1* = !ABI2 & ROS
GPA1* = S1P_PhytoS1P
GTP* = 1
H2O_Efflux* = AnionEM & AquaporinPIP2_1 & K_efflux & !Malate
HAB1* = !RCARs | !ROS
H_ATPase* = !Ca2osc
InsP3* = PLC & PtdIns4_5P2
InsP6* = InsP3
KEV* = Ca2osc
KOUT* = Depolarization & !ROS | Depolarization & pHc
K_efflux* = KEV & KOUT
MPK9_12* = Ca2osc
MRP5* = 1
Malate* = !ABA & !AnionEM & PEPC
Microtubule_Depolymerization* = TCTP | PA
NAD* = 1
NADPH* = 1
NIA1_2* = ROS
NO* = NADPH & NIA1_2 & Nitrite
NOGC1* = NO
Nitrite* = 1
NtSyp121* = 1
OST1* = !ABI1 & !ABI2 | !ABI1 & !PP2CA | !HAB1
PA* = PC & PLDdelta | PC & PLDalpha | DAG & DAGK
PC* = 1
PEPC* = !ABA
PI3P5K* = ABA
PLC* = Ca2osc
PLDalpha* = GPA1
PLDdelta* = GAPC1_2 & NO & ROS
PP2CA* = !RCARs
PtdIns3_5P2* = PI3P5K
PtdIns4_5P2* = PtdInsP4
PtdInsP3* = 1
PtdInsP4* = 1
QUAC1* = OST1
RBOH* = !ABI1 & GPA1 & OST1 & PtdInsP3 & RCN1 & pHc | !ABI1 & OST1 & Actin_Reorganization | GPA1 & OST1 & Actin_Reorganization | OST1 & PA & Actin_Reorganization | PA & PtdInsP3 & RCN1 & Actin_Reorganization | RCN1 & pHc & Actin_Reorganization
RCARs* = ABA
RCN1* = 1
ROP11* = GEF1_4_10
ROS* = NADPH & RBOH & AquaporinPIP2_1
S1P_PhytoS1P* = SPHK1_2 & Sph

SACC* = Actin_Reorganization
SCAB1* = 1
SLAC1* = !ABI1 & !ABI2 & CPK23 & CPK6 & MPK9_12 & pHc | !ABI1 & !ABI2 & CPK3_21 & CPK6 & MPK9_12 & !PP2CA & pHc | !ABI1 & !ABI2 & CPK3_21 & GHR1 & MPK9_12 & pHc | !ABI2 & CPK3_21 & MPK9_12 & OST1 & pHc
SLAH3* = !ABI1 & CPK23 & CPK3_21
SPHK1_2* = ABA
SPP1* = 0
Sph* = 1
TCTP* = Ca2osc
V-ATPase* = Ca2osc
V-PPase* = PtdIns3_5P2
Vacuolar_Acidification* = V-ATPase | V-PPase | pHc
cADPR* = ADPRc & NAD
cGMP* = GTP & NOGC1
pHc* = !ABI1 & OST1 & Vacuolar_Acidification | Ca2osc & OST1 | Ca2osc & Vacuolar_Acidification

### Text S11. Intrinsic limits on the agreement between experiments on ABA-induced closure and models

We analyzed the discrepancies between experimental observations and the results of GA-refined models and identified several types of causes. The first type is made up of inconsistencies between experimental results that cannot be resolved by any model. The second type represents quantitative effects that cannot be reflected by a Boolean model. The third type has to do with the limitations imposed by the interaction network.

**Inconsistencies between experimental results:**

The molecules DAG and InsP3 are the two products of the PLC-catalyzed hydrolysis of PtdIns(4,5)P2. As they are produced by the same reaction, their level should be the same. There is no improvement to the model that can recapitulate both the observation of ABA-induced InsP3 production [80] and the observation of no/very little DAG in the presence of ABA [81].

The pair of observations that (i) depletion of ROS impairs ABA-induced closure [82] and (ii) *gpa1* KO plants do not produce ROS in response to ABA [83], is not compatible with the observation that GPA1 KO leads to wild type ABA-induced closure [84].

**Quantitative effects that cannot be reflected by a Boolean model:**

The Boolean representation of an enzyme-catalyzed reaction requires the activity of the enzyme for the reaction to proceed and yield product. The experimental observation that providing nitrite (the reactant of the reaction that yields NO) leads to stomatal closure [85] cannot be captured in this framework. The likely resolution of the discrepancy is that a non-physiologically large amount of nitrite was provided.

$Ca^{2+}$ release from stores (CIS) and $Ca^{2+}$ flow through the membrane (CaIM) are the two mechanisms of increase in the cytosolic $Ca^{2+}$ level. Any Boolean function would assign a symmetrical influence to them (either both are sufficient, or both are necessary). In contrast, experimental results suggest an unequal contribution of CIS and CaIM [86].

The model characterizes the physiological processes that determine stomatal closure (e.g. $K^+$ efflux, $H_2O$ efflux, Microtubule depolymerization) as necessary regulators of closure, whose disruption leads to Closure=0. Experimental results in which disruption of one of these processes led to reduced (but not absent) ABA-induced closure [76] cannot be recapitulated by the model.

**Limitations imposed by the interaction network:**

If the network contains a linear pathway A→B→C, the results of activating A, B, or C should be the same, as are the results of inhibiting A, B, or C. There were cases in which the perturbation of two nodes that participate in a linear pathway did not yield the same experimental result. For example, providing 8-nitro-cGMP led to a qualitatively smaller degree of closure than providing cADPR, which cannot be captured by the model's 8-nitro-cGMP→ADPRc→cADPR linear pathway. The differences may be due to quantitative effects or to missing edges.

"Actin reorganization" is a node that represents a two-step process: the disassembly and turnover of actin filaments, then their reassembly in a different configuration. The protein SCAB1 bundles and stabilizes actin filaments. The experimental observations that either knockout of SCAB1 or overexpression of SCAB1 delays actin reorganization [87] can only be captured if the two steps of the process are represented by separate nodes.

### Text S12. Summary of the scores and features of the genetic algorithm-modified models as compared to the original 2017 model of ABA-induced stomatal closure.

The name of each model that was refined through the genetic algorithm starts with GA.
The 2017 ABA model's score of 310.5/505 (61.5%) is smaller than the original evaluation of the model (95/112=85%) because our compilation of experiments is larger and our scoring criteria are different.
"VA-KEV del" refers to the 2017 model corrected by deleting an erroneous edge that targets KEV. "GA - VA-KEV del" represents the outcome of the genetic algorithm when starting from the VA-KEV del model. Li (2006) refers to the earliest version of the model of ABA-induced stomatal closure [28]. Waidyarathne (2018) [38] is a model that builds on and expands the Li et al. model. Maheshwari (2019) [30] is a follow-up to the 2017 model. Baseline-A represents the corrected 2017 model (VA-KEV del) augmented with a self-edge on $H_2O$ efflux. Baseline-B represents the model that replaces the $Ca^{2+}{}_c$ and $Ca^{2+}$ ATPase nodes with a merged Ca2+osc node. The GA0 models are GA-improved versions of the respective baselines but without allowing additional edges. The GA1 models allow the addition of edges from a pool of 13 experimentally-supported edges. The GA2 models start from the respective GA1 model and allow the addition of 8 more edges.

| Model | Score | Notes |
|---|---|---|
| Albert (2017) | 310.5 / 505 (61.5%) | wrong regulatory edge for KEV |
| VA-KEV del | 291.7 / 505 (57.8%) | failed to achieve ABA induced closure |
| GA - VA-KEV del | 385.3 / 505 (76.3%) | failed to achieve ABA induced closure |
| Li (2006) | 135.3 / 248 (54.5%) | 39 nodes shared with the 2017 model |

| | | |
|---|---|---|
| Waidyarathne (2018) | 175.8 / 296 (59.4%) | 49 nodes shared with the 2017 model |
| Maheshwari (2019) | 240.0 / 385 (62.3%) | 47 nodes shared with the 2017 model |
| baseline-A | 311.7 / 505 (61.7%) | |
| baseline-B | 184.2 / 505 (36.5%) | alternate $Ca^{2+}$ representation |
| GA0-A | 390.8 / 505 (77.4%) | no extra edges |
| GA0-B | 384.6 / 505 (76.2%) | no extra edges, alternate $Ca^{2+}$ representation |
| GA1-A | 426.9 / 505 (84.5%) | |
| GA1-B | 407.9 / 505 (80.8%) | alternate $Ca^{2+}$ representation |
| GA2-A | 431.2 / 505 (85.4%) | more assumed edges, started from GA1-A |
| GA2-B | 412.8 / 505 (81.7%) | alternate $Ca^{2+}$ representation, more assumed edges, started from GA1-B |

### Text S13. Considering BoNesis or Gitsbe as alternatives to derive a Boolean model from the experimental data on ABA-induced stomatal closure

BoNesis: The BoNesis Python library uses Answer Set Programming for the synthesis of Boolean models based on an interaction network and information describing advanced dynamical properties, such as reachability, bifurcation, minimal trap spaces, stable states, and mutations. Its main application domain is the inference of Boolean models from gene expression data of cellular differentiation and reprogramming processes. We investigated whether the interaction network, edge constraints, and experimental information available for the ABA induced closure system is suitable for BoNesis to infer a Boolean model. The details of this analysis are indicated in the github page https://github.com/kyuhyongpark/boolmore/blob/benchmarks/bonesis/comparison/bonesis/README.md. Here we summarize the key points.

Most of the edge constraints (with the exception of "necessary") can be given to BoNesis by using the AEON file format. As BoNesis can only use purely binary input, experimental observations in an intermediate category had to be left out from the input information. We used the closest interpretation allowed by BoNesis for the remaining experiments. For example, observing "Closure=ON" in the presence of ABA indicates that all trap spaces of the model should have Closure=1. BoNesis does not have a constraint that has this exact meaning. The closest is to require that (i) there exists a trap space with Closure=1, and (ii) all fixed points, if they exist, have Closure=1.

Running the satisfiability check of BoNesis on the compilation of experimental results indicated that no Boolean model can satisfy them all. Therefore, we further restricted the list of experiments, considering only those that were correctly reproduced by either a base model or a GA-improved variant. Such a set of experiments is satisfiable by a documented model. A 24-hour run of BoNesis did not return any solutions. These results indicate that the application domain of BoNesis does not extend to compilations of small-scale perturbation experiments on signal transduction networks.

Gitsbe: The algorithm Gitsbe (Generic Interactions to Specific Boolean Equations) is part of the DrugLogics software package developed by A. Flobak and collaborators [34]. Its inputs are a prior knowledge network and a set of observed steady states for at least a subset of the nodes (i.e. a set of partial fixed points). Gitsbe starts with a default regulatory function in which inhibitors dominate over activators, and then uses a genetic algorithm to eliminate regulators if that improves agreement with the observations. Gitsbe was used as a component of a pipeline to determine synergistic cancer drug combinations [34,88]. We investigated whether the interaction network and experimental information available for the ABA induced closure system is suitable for Gitsbe to infer a Boolean model. The details of this analysis are indicated in the github page https://github.com/kyuhyongpark/boolmore/tree/master/comparison/gitsbe/README.md Here we summarize the key points.

We specified the prior knowledge network corresponding to baseline model A in the closest form interpretable by Gitsbe. Accordingly, we included the experimentally-supported extra edges as part of the initial network, and eliminated the source nodes with constant values. We generated 18 models with Gitsbe and 15 with *boolmore* and scored them on the 429 experimental observations that could be processed by Gitsbe. The best-scoring Gitsbe model has a lower score - 305.0/429 (71.1%) - than the worst-scoring *boolmore* model - 334.3/429 (77.9%). Furthermore, all the Gitsbe-generated models have a biologically invalid function for the node Closure, keeping Microtubule depolymerization as its sole regulator and eliminating its dependence on $H_2O$ efflux. Water efflux from the guard cells is the cause of their deflation, which is a necessary condition of stomatal closure. *Boolmore* can enforce the preservation of key regulators by its use of constraints (as described in Text S9). Gitsbe does not have this capability.

### Text S14. Regulatory function revisions shared by the two GA-improved models

The two GA-refined models preserve the regulatory functions of 30 nodes; the remaining 28 regulatory functions were changed in one or both models. A significant fraction of the modifications to regulatory functions are shared by both models (6 identical modifications and 13 very similar ones).

A general trend in the shared modifications is the deletion of an assumed direct regulatory relationship that can also be explained by a logic chain (a succession of two or more relationships that has the same effect). In some of these cases (e.g., in the regulatory function of 8-nitro-cGMP), we could identify a shortcoming in the reasoning that led to the regulatory function in the original model. Ten edges were deleted in both GA models: $Ca^{2+}_c$→QUAC1, NO⊣KOUT, ROS→8-nitro-cGMP, ROS⊣$H^+$ ATPase, ROS⊣PP2CA, SCAB1→Actin Reorganization (omitted from Figure 4), SLAH3→AnionEM, and self loops on Vacuolar Acidification, MPK9/12 and Microtubule Depolymerization.

Five shared modifications use new edges from the experimentally supported pool, confirming the improvements possible from the incorporation of new biological information. These edges are Aquaporin→ ROS, ROS→ Actin reorganization, PA→ Microtubule depolymerization, PA-| ABI2, $pH_c$→Vacuolar acidification. In the following we describe 10 shared modifications, their support in the experimental literature, and the insights they provide.

ROS* = Aquaporin(PIP2;1) & RBOH

Original: ROS* = RBOH
Both models adopt the experimentally supported Aquaporin(PIP2;1)→ ROS edge [71,72] from the pool.

Microtubule Depolymerization* = PA (GA1-A)
Microtubule Depolymerization* = PA | TCTP (GA1-B)
Original: Microtubule Depolymerization* = TCTP | Microtubule Depolymerization
Both models adopt the experimental-evidence-supported edge from PA to Microtubule depolymerization [70,76] from the pool as a sufficient mechanism. Specifically, [76] indicates that PA leads to microtubule depolymerization even when the cytosolic $Ca^{2+}$ level is clamped (which would abolish the activation of TCTP, the other regulator of the node Microtubule Depolymerization).

ABI2* = !PA (GA1-A)
ABI2* = !$Ca^{2+}$ osc & !PA & !RCARs | !$Ca^{2+}$ osc & !ROS (GA1-B)
Original: ABI2* = !RCARs & !ROS
Both models adopt an indirect (GA1-A) or both direct and indirect (GA1-B) inhibitory effect from $Ca^{2+}_c$ to ABI2, recapitulating the experimental results and prediction from Maheshwari et al. [30]. GA1-A couples this with the loss of the two original inhibitors. This loss does not significantly affect the settings under which ABI2 is activated, as PA is activated by ABA or ROS.

Vacuolar Acidification* = V-PPase & $pH_c$ | V-ATPase (GA1-A)
Vacuolar Acidification* = $pH_c$ | V-PPase | V-ATPase (GA1-B)
Original: Vacuolar Acidification* = Vacuolar Acidification | V-PPase | V-ATPase
Interpretation: The 2017 model included two vacuolar proton pumps (the V-ATPase and the V-PPase) as independent activators of vacuolar acidification. As experimental evidence indicates the existence of a positive feedback loop between the cytosolic and vacuolar pH level [77], we included an edge from $pH_c$ to Vacuolar acidification to the pool. Both models incorporate this edge; both also omit the assumption of self-sustained activity in the 2017 model. The coupling of the effect of $pH_c$ and V-PPase is different in the two functions.

Actin Reorganization* = ARP Complex & !AtRAC1 & PtdInsP3 & PtdInsP4 | ARP Complex & !AtRAC1 & PtdInsP3 & ROS | PtdInsP3 & PtdInsP4 & ROS (GA1-A)
Actin Reorganization* = ARP Complex & !AtRAC1 & PtdInsP3 & PtdInsP4 & ROS (GA1-B)
Original: Actin Reorganization* = (PtdInsP4 | PtdInsP3) & ! AtRAC1 & ARP Complex & SCAB1
Both GA1-A and GA1-B add ROS as a regulator of actin reorganization, but ROS combines with !AtRAC1 with “or” in GA1-A and with “and” in GA1-B. This means that actin reorganization can happen in response to ROS in GA1-A but only in response to ABA in GA1-B. In addition, the effect of SCAB1 on Actin reorganization is lost in both models. This is probably due to the fact that both SCAB1 KO and SCAB1 overexpression only delay actin reorganization [87] .

8-nitro-cGMP* = NO & cGMP
Original: 8-nitro-cGMP* = NO & ROS & cGMP
The published model incorporated the evidence that providing NO or ROS leads to the production of 8-nitro-cGMP, and ROS KO inhibits ABA-induced production of 8-nitro-CGMP

[89], by assuming that both ROS and NO are direct and necessary regulators of 8-nitro-cGMP (in addition to its substrate, cGMP). The GA-improved function indicates that this choice was not optimal. ROS does not need to be a regulator of 8-nitro-cGMP; its sufficient and necessary role in 8-nitro-cGMP production is established via its sufficient and necessary role for NO and cGMP production (via the linear pathway ROS→NIA1/2→NO→NOGC1→cGMP). In addition, assuming ROS to be a direct and necessary regulator of 8-nitro-cGMP contradicts the evidence that NO (which does not lead to ROS production [82]) is sufficient for 8-nitro-cGMP production.

PP2CA* = !RCARs
Original: PP2CA* = !RCARs & !ROS
Interpretation: The 2017 model assumed that ROS inhibits PP2CA because of the evidence that ROS inhibits ABI1, ABI2 and HAB1. The loss of the regulation by ROS, shared by both GA-improved models, suggests that this assumption is not necessary.

QUAC1* = OST1
Original: QUAC1* = $Ca^{2+}_c$ & OST1
Both models omit the regulatory effect of cytosolic $Ca^{2+}$ on the QUAC1 anion channel. This makes the function easier to satisfy and makes sustained QUAC1=1 possible in GA1-A. The 2017 model assumed that $Ca^{2+}_c$ regulates QUAC1 based on the loss of $Ca^{2+}$-induced closure for QUAC1 KO reported in [90]. Both GA1-A and GA1-B recapitulate this observation without having the edge. The lack of necessity for the direct regulation is supported by the experimental observation in [91] of anion current through QUAC1 in response to ABA when the $Ca^{2+}$ level was clamped to 110 nM (which corresponds to the resting $Ca^{2+}_c$ level).

The function of SLAC1 is easier to satisfy.
The published model's regulatory function for SLAC1 incorporated 9 documented direct regulators of the SLAC1 ion channel as well as indirect evidence that MPK9/12 knockout impairs $Ca^{2+}_c$-induced activation of SLAC1. The three CPK nodes were assumed to be interchangeable (were connected by "or"); all the other regulators were assumed to be necessary, meaning that the regulatory function of SLAC1 contained three 8-variable disjunctive ('and'-connected) clauses. The improved models have 4-5 clauses and have fewer (5-7) variables included in each clause.

AnionEM* = QUAC1 & SLAC1
Original: AnionEM* = QUAC1 & SLAH3 | SLAC1
This function omits SLAH3 and makes SLAC1 and QUAC1 necessary for AnionEM. This shared modification of the improved models suggests that although SLAH3 is a valid anion channel, its contribution to AnionEM is much weaker than that of the other two. This prediction is consistent with the finding that SLAH3 knockout does not diminish ABA-induced stomatal closure [92]. The Boolean function of the published model, which assumed that QUAC1 and SLAH3 together can lead to sufficient anion efflux, was suboptimal, as it contradicted the experimental finding that SLAC1 KO eliminated ABA-induced closure [93].

### Text S15. Details of the experimentally testable biological predictions presented in Table 4

A prediction can be extracted from the shared modification to the regulatory function of OST1. Both GA-improved models recapitulate the experimentally observed inhibitory effect of ABI1 on QUAC1 by making ABI1 alone able to inhibit OST1 kinase activity. This results in the follow-up prediction that ABI1=1 leads to the insensitivity of Aquaporin to ABA; this prediction can be experimentally verified via methods used in [94].

The GA-refined models modify the regulatory function of the anion channel SLAC1 such that it is easier to activate. As a consequence, they recapitulate the experimental observation that ROS activates SLAC1. A follow-up prediction is that the resulting anion flow brings the malate concentration below threshold.

Both GA-refined models modify the regulatory function of ROS by incorporating the documented role of aquaporins in bringing ROS into the cell. The necessity of this process for achieving a level of ROS sufficient to regulate its downstream targets implies that the disruption of aquaporins (in *pip2;1* KO mutants) will disrupt NO production, cADPR production (one of the sources of CIS), and the activation of PLD$\delta$ (one of the enzymes that catalyze PA production). Any impairment of NO production in *pip2;1* KO mutants in response to ABA or other closure-inducing signals can be experimentally tested using DAF2 fluorescence.

Yet another prediction is due to the shared features of the regulatory functions of PP2CA in the two models: PP2CA will be active in the absence of ABA or for knockout of the RCARs receptors even if ROS or other internal closure-inducing signal is provided. This prediction is testable by, e.g., assaying PP2CA phosphatase activity in response to ROS.

Causal relationships mediated by chains of interactions (pathways) can also yield new predictions. The 2017 model and the GA-refined models agree in predicting that ROS is sufficient to induce PA production. This is experimentally testable. As the GA-refined models incorporate the new observation that PA is sufficient for microtubule depolymerization, a follow-up testable prediction is that ROS can induce microtubule depolymerization; this can be tested by methods used in [39].

The shared features of the GA-refined models' minimal trap spaces identify further predictions (see Text S16 for a description of these minimal trap spaces). The strong similarity of the GA models' minimal trap spaces in response to providing NO, 8-nitro-cGMP, or cADPR suggests that the experimental observations of three elements ($pH_c$, ROS, PA) made in the case of providing NO can be extrapolated to the other interventions, for which no such assays were performed yet. Perhaps the most interesting of these observations is the lack of production of ROS under externally provided NO [82]. We find that the GA-refined models' minimal trap spaces in response to $Ca^{2+}$ share this feature with the minimal trap spaces found in response to NO. This result leads to the prediction that external $Ca^{2+}$ would yield no or a very limited amount of ROS production. While ROS production in ABA-induced closure has been experimentally documented, the production of ROS in response to external $Ca^{2+}$ has not yet been studied experimentally.

### Text S16. The dynamics of the GA-refined models

Both GA1-A and GA1-B have a single minimal trap space (attractor) in the presence of ABA; these attractors recapitulate the experimental observations of the wild-type response to ABA and are similar to each other. The differences are in the nodes that oscillate in GA1-A due to the oscillations in $Ca^{2+}_c$ and stabilize in their active state in GA1-B due to the activity of the $Ca^{2+}$osc node. Both GA-refined models have a single minimal trap space in the absence of ABA, in contrast to the original model, which has 17 minimal trap spaces. There is an almost perfect agreement between the minimal trap space of GA1-A and GA1-B. A significant agreement also exists between the minimal trap space of GA1-A and GA1-B in the case of ROS-induced stomatal closure.

In contrast, there is a significant difference between the minimal trap spaces in case of externally provided $Ca^{2+}$. We found that the 2017 model has two attractors. Both of these attractors exhibit oscillations in $Ca^{2+}_c$ but they differ in the state of 33 nodes, including Closure. Only one of the attractors recapitulates the experimentally observed robust closure in response to external $Ca^{2+}$ [95]. In contrast, each of GA1-A and GA1-B leads to a single attractor. The attractor of GA1-A features the oscillation of 47 nodes, including Closure. The attractor of GA1-B contains the ON state of the node Closure as well as of documented determinants of closure such as AnionEM. Considering the 6 elements whose status was measured under externally provided $Ca^{2+}$ ($pH_c$, KEV, Microtubule depolymerization, $Ca^{2+}_c$, SLAC1, closure), GA1-A recapitulates 3 measurements and GA1-B recapitulates all 6, thus it is the best among the three models in terms of agreement with experiments.

There is a significant difference between minimal trap spaces in the case of providing NO in the absence of ABA as well. The two GA-refined models rely on different mechanisms to achieve agreement with the experimentally observed moderate closure. GA1-A has the same attractor as in the case of providing external $Ca^{2+}$; this attractor has many oscillating nodes, including Closure. GA1-B has two attractors; one has $Ca^{2+}_c$osc = Closure = 0 and the other has $Ca^{2+}_c$osc = Closure = 1. Considering the 5 elements whose status was assayed under externally provided NO ($pH_c$, ROS, 8-nitro-cGMP, PA, closure), GA1-A recapitulates four experiments, and GA1-B recapitulates all five.

### Text S17. Adapting *boolmore* to multi-level models

Variables with multiple discrete levels are used if the experimental observations fall into more than two categories. For example, experimental observations may indicate a baseline level, a higher-than baseline level, and a low (lower than baseline) level. These observations are represented by assigning state 1 to the baseline level, state 2 to the high (above-baseline) level, and state 0 to the low (below-baseline) level. *Boolmore* can be readily adapted to incorporate multi-level variables by representing such variables by multiple Boolean variables, refining the resulting Boolean model, then mapping groups of Boolean variables back to multi-level variables.

**Defining the Boolean variables and regulatory functions:**

Any node with multiple discrete states (activity levels) can be represented as a set of nodes with Boolean variables [41,42]. Specifically, for any variable x with the levels 0, 1, … m, the so-called van Ham mapping associates m Boolean variables. The first Boolean variable, $b_1$, is 0 if x<1 and is 1 if x≧0, the second Boolean variable, $b_2$, is 0 if x<2 and is 1 if x≧2, and so on. The constraint of the allowed configurations is that the state 1 of a Boolean variable that corresponds to a higher level of x requires the state 1 of all Boolean variables that correspond to lower nonzero levels of x. This allows m+1 configurations of the Boolean variables, having one-to-one correspondence with the m+1 discrete levels of x. Table A shows the van Ham mapping of a three-level variable. Didier et al. proved that this mapping is the sole Boolean mapping, up to cosmetic changes (e.g., by switching 0 and 1), that preserves both the structure of the regulatory network and its dynamical behaviors. We introduce the set of Boolean variables corresponding to the van Ham mapping of each multi-level variable, and express the obligatory relationships between them as constraints. In the example of Table A, the constraint is that $b_1$ is a necessary regulator of $b_2$. We can then use our binary representation for the function of each Boolean variable, and then implement the mutations on this representation as currently done.

| y | z | x* | $b_1$* | $b_2$* |
|---|---|---|---|---|
| 0 | 0 | 0 | 0 | 0 |
| 0 | 1 | 1 | 1 | 0 |
| 1 | 0 | 1 | 1 | 0 |
| 1 | 1 | 2 | 1 | 1 |

Table A. Illustration of the van Ham mapping of a three-level variable x, regulated by two binary variables (y and z), into the Boolean variables $b_1$ and $b_2$. The regulatory function of x can be expressed as “y+z”. The regulatory function of $b_1$ is “y OR z”; the regulatory function of $b_2$ is “y AND z”. In our application of *boolmore* we enforce $b_1$ to be necessary for $b_2$, so that configurations such as “$b_1$=0, $b_2$=1” are not allowed.

The *boolmore*-refined Boolean model can be mapped back to a model with multi-level variables by using the van Ham mapping in reverse. This mapping will yield a table for each multi-level variable. The entries of the table can be expressed via multiple algebraic formalisms [42,96].

**Adapting the representation of perturbations and experimental observations:**

Knockout of a multi-level node x is represented by setting the Boolean variable $b_1$ to 0. The obligatory relationships between Boolean variables, expressed as constraints to the Boolean functions, ensure that the Boolean variables $b_2$= .. =$b_m$ =0. Constitutive activation of a multi-level node is represented by setting each Boolean variable corresponding to it to 1 (see the last row of Table A).

The experimentally observed levels form the basis of the multi-level variable. Observation of the highest level (m) is represented by the state 1 of the Boolean variable $b_m$. By construction, this state requires the state 1 of all the lower-level Boolean variables (see the last row of Table A). Observation of the lowest level (0) is represented by the state 0 of the Boolean variable $b_1$, which ensures the state 0 of all Boolean variables (see the first row of Table A). Observation of an intermediate level i of the multi-level variable is represented by the combination of the state 1 of the Boolean variable $b_i$ with the state 0 of the Boolean variable $b_{i+1}$ (see the second and third row of Table A). The states of the Boolean variables are added as separate elements of the same entry of the table of experimental observations, but their coupled nature is incorporated during scoring.

An observation of the lowest level (0) or an observation of the highest level (m) can only be captured by a single state of the Boolean variables in the model. This state receives a score of 1; all other states receive a score of 0. As there can be some uncertainty in the classification of an observed intermediate state, a partial scoring is warranted. For example, if the experimentally observed state of variable x of Table A is 1, which is represented as two observations for the Boolean variables, $b_1$=1, and $b_2$=0, a partial score (e.g. 0.5) may be given if one of these observations is captured. This can be interpreted as giving a half score for having x activated to level 1, and giving a half score for not having x activated to level 2.

**Case study:**

We translated the three-state model of *C. elegans* nutrition-based lipase regulation [43] to a Boolean model, then refined it with *boolmore.* The model has 12 nodes, including two source nodes that describe nutritional context: “food” and “oxidative stress”. The two source nodes and their receptors are described by Boolean variables. The remaining 8 nodes have three states: basal (corresponding to the level in wild type animals under *ad libitum* feeding), lower than basal (corresponding to knockout or an inhibited state), and higher than basal (corresponding to upregulation due to fasting or oxidative stress). The regulation of the three-level nodes was reported via tables in [43]. Many of these tables did not cover all the input combinations, as there were no observations to inform or constrain the outcome in these cases. The interaction network was published as Figure 7H in Mony et al. [43].

We translated each three-level node into two Boolean nodes. For example, the multi-level node lipl-3 was represented by the Boolean variables lipl-3_basal and lipl-3_high. We determined the values of the Boolean variables using van Ham mapping of the input-output tables of the three-level nodes, and identified the Boolean functions that best fit these values. This Boolean model served as the starting model. Note that this starting model has fewer edges in its interaction graph than we would generally want. For example, considering the relationship of the mRNA mxl-3 and its protein MXL-3, in the starting model MXL-3_basal is regulated by mxl-3_basal but not by mxl-3_high. To allow *boolmore* to freely mutate the functions to allow for each pair of regulator and target that the basal or the high level of the regulator affects the basal or high level of the target , we created a baseline model that  incorporated all possible such edges among Boolean variables. Thus, the baseline model has four edges that correspond to the translation of the protein MXL-3, starting either from mxl-3_basal or mxl-3_high and ending in  MXL-3_basal or MXL-3_high. This setup allows *boolmore* to start the genetic algorithm from the starting model, but freely add edges present in the baseline model (even if they are not present in the starting model).

We used the information regarding  regulatory effects) documented in the article as constraints to the Boolean functions. These include the transcriptional regulation of lipl-3 by HLH-30, the obstruction of HLH-30 activity by MXL-3, and the known information that the insulin-bound receptor DAF-2 activates DAF-16. The relationship of an mRNA to its protein is included in the `necessary` category. Another set of constraints specifies that the Boolean variable that expresses basal expression is a necessary regulator of the Boolean variable that expresses higher-than-basal expression. Finally, as few experimental observations document the conditions for the basal level of DAF-16 and lipl-3, the corresponding Boolean variables are allowed to become constants. All other nodes must retain at least one regulator.

We compiled the results of 81 perturbation experiments from the article (14 results cited from prior literature and the rest reported for the first time). We interpreted the perturbations (knockouts and overexpressions) and observed states as described above. In scoring an observed basal level of variable X, we allowed a partial score of 0.5 for agreement with X_basal=1 and a partial score for agreement with X_high=0.

We found that the starting model’s accuracy was only 52%. We identified that the reason for this low score is that in the presence of food and absence of oxidative stress the positive feedback loop between three nodes (mxl-3, MXL-3, and HSF-1) led to two minimal trap spaces. The application of *boolmore* improved the score to 86% by changing the function of 9 of the 20 Boolean variables. One of the changes was to eliminate the Boolean variable MXL-3_basal from the function of HSF-1_basal, thus disrupting the positive feedback loop and yielding a single minimal trap space in all contexts.

In summary, our case study demonstrates that *boolmore* can be readily adapted to refine multi-level models. All necessary data to run *boolmore* and a detailed report of the performance of the *boolmore*-refined model are available in the github repository https://github.com/kyuhyongpark/boolmore/tree/documentation/case_study/c_elegans

### Table S1. The agreement functions used in our workflow

The agreement function indicates the average value of the respective node in the trap spaces of the model on the x axis and the agreement on the y axis. These are used for hierarchy scoring, as described in Text S3.

| Category | Agreement function | Examples |
|---|---|---|
| ON | 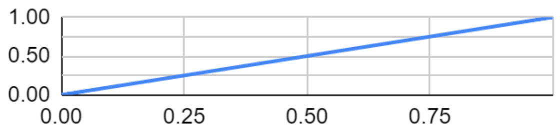  | ABA=1, observing $pH_c$. |
| Some/ON | 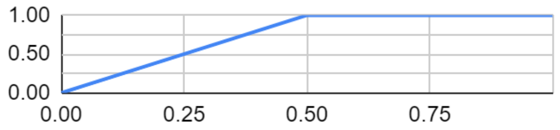  | NO CA, observing Closure. The closure (i.e., reduction in the stomatal aperture) in various reported experiments varied between 30% and 60% of that in response to ABA. |
| Some | 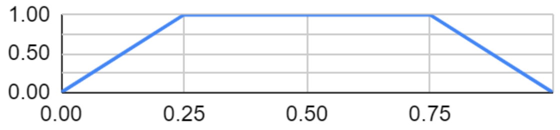  | 8-nitro-cGMP CA, observing Closure. The experimental closure response was around 25% of the response to ABA. |
| OFF/Some | 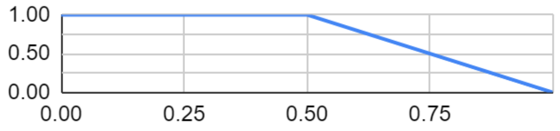  | ABA=1, S1P/PhytoS1P KO, observing Closure. S1P was depleted using a chemical, whose effect may reduce over time. Thus, the experimentally observed weak closure (instead of the expected lack of closure) may be due to the dissipation of the depletion. |
| OFF | 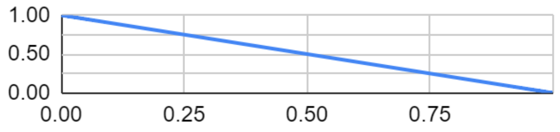  | NO CA, observing ROS. |

### Table S2. Detailed benchmark results

The starting accuracies and the final accuracies are averages of 5 independent runs. We used models from the Cell Collective with size (N) less than 30. We generated artificial training sets of size 10*N, from which a randomly selected 8*N were used as the training set, and the remaining 2*N were used as the validation set. The accuracy is calculated by dividing the score of the model by the respective maximal score. The standard deviation is shown in parentheses.

| model name | size (N) | training start (%) | training final (%) | validation start (%) | validation final (%) |
|---|---|---|---|---|---|
| Arabidopsis thaliana Cell Cycle | 14 | 13.7(±11.3) | 93.1(±2.2) | 7.5(±8) | 87.4(±3.5) |
| Aurora Kinase A in Neuroblastoma | 23 | 38.8(±17) | 98.2(±1.9) | 37(±20.1) | 95.6(±5.6) |
| B cell differentiation | 22 | 44.2(±11.8) | 99.3(±1.3) | 44.7(±15) | 98.4(±1.5) |
| BT474 Breast Cell Line Long-term ErbB Network | 24 | 49.9(±13.8) | 99.5(±0.8) | 50.5(±6.4) | 95.2(±4.6) |
| BT474 Breast Cell Line Short-term ErbB Network | 16 | 49.9(±9.9) | 100(±0) | 46.1(±17.5) | 96.4(±4) |
| Budding Yeast Cell Cycle | 20 | 60.7(±11.4) | 99.4(±0.8) | 55.1(±18.4) | 95(±4.5) |
| Budding Yeast Cell Cycle 2009 | 18 | 23.7(±19.4) | 92.8(±4.6) | 18.8(±17.1) | 84.3(±5.2) |
| CD4+ T Cell Differentiation and Plasticity | 18 | 41.9(±5.8) | 96.3(±1.7) | 44.1(±13.1) | 84.5(±6.2) |
| Cardiac development | 15 | 67.3(±5.7) | 100(±0) | 72(±12.6) | 97.7(±3.2) |
| Cell Cycle Transcription by Coupled CDK and Network Oscillators | 9 | 39.9(±21.9) | 100(±0) | 30.8(±26.6) | 100(±0) |
| Cortical Area Development | 5 | 74.8(±11.2) | 99.5(±1.1) | 73.3(±13.2) | 93(±4.5) |
| Death Receptor Signaling | 28 | 50.9(±12.4) | 97(±4.1) | 43.4(±15.6) | 96.6(±4.6) |
| FA BRCA pathway | 28 | 29.9(±16.9) | 94.2(±2.9) | 25.1(±14.9) | 87.2(±4.1) |
| FGF pathway of Drosophila Signalling Pathways | 23 | 79.9(±9) | 100(±0) | 78.7(±9.6) | 98.7(±2.9) |
| Fanconi anemia and checkpoint recovery | 15 | 38.5(±7.5) | 96.5(±2.5) | 37(±7.1) | 79.7(±9.3) |
| HCC1954 Breast Cell Line Long-term ErbB Network | 23 | 61.8(±6.4) | 99.1(±1.4) | 64.4(±8.3) | 97.4(±2.1) |
| HCC1954 Breast Cell Line Short-term ErbB Network | 16 | 49.8(±6.6) | 99.9(±0.2) | 49(±13) | 97.2(±1.6) |
| HH Pathway of Drosophila Signaling Pathways | 24 | 60.1(±11) | 98.6(±1.9) | 61.2(±13.4) | 95(±6.7) |
| Human Gonadal Sex Determination | 19 | 50.3(±16.1) | 97.5(±2.5) | 45.8(±21.2) | 89.7(±9.3) |
| Iron acquisition and oxidative stress response in aspergillus fumigatus | 22 | 49.2(±30.2) | 99.2(±1.5) | 49.1(±30.5) | 96.6(±2.1) |
| Lac Operon | 13 | 39(±17.9) | 99.4(±0.9) | 29.6(±22.3) | 97.3(±4.2) |
| Mammalian Cell Cycle | 20 | 32.5(±10.6) | 98.1(±1.4) | 25(±14.8) | 90(±5.1) |

| | | | | | |
|---|---|---|---|---|---|
| Mammalian Cell Cycle 2006 | 10 | 30.8(±11) | 98.9(±1.1) | 28.7(±9.5) | 94.5(±4.5) |
| Metabolic Interactions in the Gut Microbiome | 12 | 77.1(±7.6) | 99.6(±0.9) | 74.8(±13.6) | 92.5(±8.4) |
| Neurotransmitter Signaling Pathway | 16 | 38(±18.1) | 100(±0) | 36.3(±16.7) | 100(±0) |
| Oxidative Stress Pathway | 19 | 41.7(±11.6) | 99.7(±0.4) | 42.2(±3.2) | 99.7(±0.6) |
| Predicting Variabilities in Cardiac Gene | 15 | 69.9(±3.8) | 99.8(±0.4) | 65.3(±11.9) | 95.3(±7) |
| Pro-inflammatory Tumor Microenvironment in Acute Lymphoblastic Leukemia | 26 | 50.2(±10.2) | 94(±7.8) | 47.9(±12.8) | 91.2(±7.3) |
| Processing of Spz Network from the Drosophila Signaling Pathway | 24 | 51.6(±12.2) | 99.5(±0.9) | 51.7(±12.4) | 97.9(±2.9) |
| Regulation of the L-arabinose operon of Escherichia coli | 13 | 45.9(±19.8) | 100(±0) | 43.6(±23.3) | 100(±0) |
| SKBR3 Breast Cell Line Long-term ErbB Network | 25 | 47.7(±7.2) | 99.5(±0.6) | 46.8(±8) | 95.3(±4.8) |
| SKBR3 Breast Cell Line Short-term ErbB Network | 16 | 50.6(±8.6) | 100(±0) | 44.8(±8.4) | 97.8(±1.8) |
| Septation Initiation Network | 30 | 54.5(±9.8) | 99.3(±0.5) | 49.5(±7.9) | 98.4(±1.2) |
| T cell differentiation | 23 | 63.1(±9.9) | 100(±0) | 55(±11.5) | 99.2(±0.8) |
| T-LGL Survival Network 2011 Reduced Network | 18 | 29.6(±21) | 99.2(±1.8) | 27.5(±19.1) | 98(±2.8) |
| TOL Regulatory Network | 24 | 60.6(±18.4) | 97.7(±1.3) | 57.5(±20.9) | 91.3(±3.1) |
| Toll Pathway of Drosophila Signaling Pathway | 11 | 40.2(±6.8) | 100(±0) | 36.4(±8.5) | 100(±0) |
| Trichostrongylus retortaeformis | 26 | 34.9(±15.4) | 98.6(±1.3) | 32(±11.9) | 93.3(±5.8) |
| VEGF Pathway of Drosophila Signaling Pathway | 18 | 73.2(±16.2) | 100(±0) | 69.4(±21.8) | 100(±0) |
| Wg Pathway of Drosophila Signalling Pathways | 26 | 69(±19.7) | 100(±0) | 72.1(±21.7) | 98.8(±1.7) |

### Table S3. The full names of the abbreviated node names in the ABA-induced closure models

Most of these node names are adopted from the 2017 ABA-induced closure model [29]. We also indicate the node names used in the code and in Figure 4, when they are different from the original name.

| Node name in the network | Node name used in the code | Node name in Figure 4 | Full name |
|---|---|---|---|
| 8-nitro-cGMP | | 8ncGMP | 8-nitro-cyclic guanosine monophosphate |
| ABA | | | Abscisic acid |
| ABH1 | | (not shown) | ABA hypersensitive |
| ABI1 | | | ABA (abscisic acid)-insensitive 1 |
| ABI2 | | | ABA (abscisic acid)-insensitive 2 |

| | | | |
|---|---|---|---|
| Actin Reorganization | Actin_Reorganization | AR | Actin reorganization |
| ADPRc | | (not shown) | ADP (adenosine diphosphate)-ribosyl cyclase |
| AnionEM | | | Anion efflux through the plasma membrane |
| Aquaporin(PIP2;1) | AquaporinPIP2_1 | Aquapor | Plasma membrane intrinsic protein 2;1 (Aquaporin) |
| ARP Complex | ARP_Complex | (not shown) | Actin related protein complex |
| AtRAC1 | | | small GTPase RAC1 |
| $Ca^{2+}_c$ | Ca2c | $Ca^{2+}$ | Cytosolic calcium |
| $Ca^{2+}$ ATPase | Ca2_ATPase | (not shown) | $Ca^{2+}$ ATPases and $Ca^{2+}/H^+$ antiporters responsible for $Ca^{2+}$ efflux from the cytosol |
| cADPR | | | cyclic ADP-ribose |
| CaIM | | | $Ca^{2+}$ influx across the plasma membrane |
| cGMP | | (not shown) | Cyclic guanosine monophosphate |
| CIS | | | $Ca^{2+}$ influx to the cytosol from intracellular stores |
| Closure | | | Stomatal closure |
| CPK23 | | (not shown) | Calcium-dependent protein kinase 23 |
| CPK3/21 | CPK3_21 | | Calcium-dependent protein kinases 3 and 21 |
| CPK6 | | (not shown) | Calcium-dependent protein kinase 6 |
| DAG | | (not shown) | Diacylglycerol |
| DAGK | | (not shown) | Diacylglycerol kinase |
| Depolarization | | Depolar | Plasma membrane depolarization |
| ERA1 | | (not shown) | Enhanced Response to Abscisic acid1 |
| GAPC1/2 | GAPC1_2 | (not shown) | Glyceraldehyde-3-phosphate dehydrogenase subunits 1 and 2 |
| GCR1 | | (not shown) | putative G protein–coupled receptor |
| GEF1/4/10 | GEF1_4_10 | (not shown) | Guanine exchange factors 1, 4 and 10 |
| GHR1 | | | Guard cell hydrogen peroxide resistant 1 |
| GPA1 | | | Heterotrimeric G protein α subunit 1 |
| GTP | | (not shown) | Guanosine 5'-triphosphate |
| $H^+$ ATPase | H_ATPase | | $H^+$ ATPase at the plasma membrane |
| $H_2O$ Efflux | H2O_Efflux | $H_2O$ eff | water efflux through the plasma membrane |
| HAB1 | | | Hypersensitive to ABA 1 |
| InsP3 | | | Inositol-1,4,5 trisphosphate |
| InsP6 | | (not shown) | Inositol hexakisphosphate |
| $K^+$ Efflux | K_efflux | $K^+$ eff | $K^+$ efflux through the plasma membrane |
| KEV | | | $K^+$ efflux from the vacuole to the cytosol |
| KOUT | | | $K^+$ efflux through slowly activating outwardly-rectifying $K^+$ channels through the plasma membrane |
| Malate | | | Malate |
| Microtubule Depolymerization | Microtubule_Depolymerization | MD | Microtubule depolymerization |
| MPK9/12 | MPK9_12 | | Mitogen-activated protein kinases 9 and 12 |

| | | | |
|---|---|---|---|
| MRP5 | | (not shown) | Multidrug Resistance-associated Protein 5 |
| $NAD^+$ | NAD | (not shown) | Nicotinamide adenine dinucleotide |
| NADPH | | (not shown) | Nicotinamide adenine dinucleotide phosphate |
| NIA1/2 | NIA1_2 | (not shown) | Nitrate reductase 1/2 |
| Nitrite | | (not shown) | Nitrite |
| NO | | | Nitric Oxide |
| NOGC1 | | (not shown) | Nitric Oxide dependent Guanylate Cyclase |
| NtSyp121 | | (not shown) | Tobacco syntaxin-like SNARE (soluble N-ethylmaleimide-sensitive factor) attachment protein receptors |
| OST1 | | | protein kinase OPEN STOMATA 1 |
| PA | | | Phosphatidic acid |
| PC | | (not shown) | Phosphatidyl Choline |
| PEPC | | (not shown) | Phosphoenolpyruvate carboxylase |
| pHc | | | Increase of the cytosolic pH level |
| PI3P5K | | (not shown) | Phosphatidylinositol 3-phosphate 5-kinase |
| PLC | | (not shown) | Phospholipase C |
| PLDα | PLDalpha | | Phospholipase D α1 |
| PLDδ | PLDdelta | | Phospholipase D δ |
| PP2CA | | | Protein Phosphatase 2CA |
| PtdIns(3,5)P2 | PtdIns3_5P2 | (not shown) | Phosphatidylinositol 3,5-bisphosphate |
| PtdIns(4,5)P2 | PtdIns4_5P2 | (not shown) | Phosphatidylinositol 4,5-bisphosphate |
| PtdInsP3 | | (not shown) | Phosphatidylinositol 3-phosphate |
| PtdInsP4 | | (not shown) | Phosphatidylinositol 4-phosphate |
| QUAC1 | | | QUickly activating Anion Channel1 |
| RBOH | | | NADPH oxidases AtRBOH D and F |
| RCARs | | | Regulatory Components of ABA Receptor |
| RCN1 | | (not shown) | Protein phosphatase 2A |
| ROP11 | | (not shown) | Small GTPase ROP11 |
| ROS | | | Reactive oxygen species |
| SACC | | (not shown) | Stretch activated calcium channels |
| S1P / PhytoS1P | S1P_PhytoS1P | S1P | sphingosine-1-phosphate/ phyto sphingosine-1-phosphate |
| SCAB1 | | (not shown) | Stomatal Closure-related Actin Binding protein |
| SLAC1 | | | Slow Anion Channel- associated 1 |
| SLAH3 | | | SLAC1 Homologue 3 |
| Sph | | (not shown) | Sphingosine |
| SPHK1/2 | SPHK1_2 | (not shown) | Sphingosine Kinases 1 and 2 |
| SPP1 | | (not shown) | Sphingoid Phosphate Phosphatase 1 |
| TCTP | | (not shown) | Translationally controlled tumor protein |
| V-ATPase | | (not shown) | Vacuolar proton ATPase |
| Vacuolar Acidification | Vacuolar_Acidification | VA | Vacuolar Acidification |
| V-PPase | | (not shown) | vacuolar proton pyrophosphatase |

### Table S4. Compilation of experimental results used to score the ABA-induced closure model

In our compilation of 505 experiments 244 assayed stomatal aperture and 261 assayed other nodes of the network.

The foundational experiment performed to investigate ABA-induced stomatal closure is to provide 20-50 μM ABA to leaf epidermal peels that were pre-incubated in a solution that opened stomata. The aperture of a population of stomata (usually hundreds) before and after treatment is measured used to establish that ABA induces a significant aperture reduction (i.e., closure). The apertures of a population of stomata under a mock treatment consisting of the solution used to deliver ABA (usually ethanol) are measured as a negative control (to establish that no closure happens).

There are two main types of experimental interventions: the genetic or pharmacological knockout of an element of the network (node) in the presence of ABA, and the external supply or activation of a node in the absence of ABA. Combinatorial interventions usually consist of providing either ABA or a node that was documented to induce stomatal closure combined with the knockout of one or multiple additional nodes. We represent the external supply or activation as “CA” (constitutive activation), and implement it as keeping the node in the ON state. We represent knockout as “KO” and implement it by keeping the node in the OFF state.

One type of experiment required a more subtle representation. $Ca^{2+}$ flow through the plasma membrane (represented by the node CaIM) can be mediated by stretch-activated channels or ROS-activated channels. Certain experiments that measured $Ca^{2+}$ flow through the membrane were not set up to monitor stretch-activated channels. To represent these experiments, we introduced the node SACC to represent stretch-activated channels, made it a regulator of the node CaIM, and assumed that it is inactive in the setting that represents the experiment.

Applying the general method  described in Text S3, the extent of reduction in the stomatal apertures in response to ABA serves as the benchmark to categorize the effect of experimental interventions. Reductions similar in magnitude to the reduction in response to ABA are categorized as “ON”, the lack of reduction observed for mock treatment defines the “OFF” category, and intermediate reductions are categorized as “Some”. Certain experiments (at least 80) were reported in multiple publications, and a fraction of these reported contradictory findings. We accommodate this variability by using the classifications OFF/Some and Some/ON. As indicated in Table S1, a range of model results are deemed consistent with such a hybrid classification, e.g., from 0.5 to 1 for Some/ON. Altogether, there are 248 cases of OFF, 53 cases of OFF/Some, 42 cases of Some, 53 cases of Some/ON and 111 cases of ON in our compilation.

All references are given in PMID, with the only exception of Qu et al. (doi.org/10.1007/s10725-017-0353-5). The key perturbations that lead to stomatal closure in the absence of ABA are underlined and italicized. Experimental observations in the presence of ABA are shown in gray background.

| ID | Source | Perturbation | Experimental result | Observed node | Category | Reference PMIDs |
|---|---|---|---|---|---|---|
| 1 | ABA=0 | WT | no closure | Closure | OFF | |
| 2 | ABA=1 | WT | closure | Closure | ON | |
| 3 | ABA=0 | <u>*ROS CA*</u> | closure(>50%) | Closure | ON | 12773379, |

| | | | | | | |
|---|---|---|---|---|---|---|
| | | | | | | 21719691,<br>22589465,<br>12446847,<br>22730405,<br>21262908,<br>16367958,<br>19690149 |
| 4 | ABA=0 | RBOH KO | no closure | Closure | OFF | 12773379,<br>15064385 |
| 5 | ABA=0 | RBOH KO, *ROS CA* | closure(>50%) | Closure | ON | 12773379,<br>16367958 |
| 6 | ABA=0 | ROS KO | no closure | Closure | OFF | 11500543,<br>16367958 |
| 7 | ABA=1 | ROS KO | no, reduced closure | Closure | OFF/Some | 11500543,<br>16367958 |
| 8 | ABA=1 | RBOH KO | reduced closure<br><br>no closure | Closure | OFF/Some | 12773379,<br>16367958,<br>15064385,<br>19690149 |
| 9 | ABA=1 | PLDα KO, RBOH KO | no closure | Closure | OFF | 19690149 |
| 10 | ABA=0 | *CaIM CA* | closure(>50%) | Closure | ON | 16664518,<br>21719691 |
| 11 | ABA=0 | $Ca^{2+}_c$ KO | no closure | Closure | OFF | 19302418,<br>24271006,<br>15064385,<br>18721267 |
| 12 | ABA=1 | $Ca^{2+}_c$ KO | reduced closure<br><br>/no closure | Closure | OFF/Some | 19302418,<br>24271006,<br>15064385<br>/18721267 |
| 13 | ABA=0 | SCAB1 CA | no closure | Closure | OFF | 21719691 |
| 14 | ABA=1 | SCAB1 CA | slower closure | Closure | Some/ON | 21719691 |
| 15 | ABA=1 | SCAB1 CA | slower actin reorganization | Actin Reorganization | Some/ON | 21719691 |
| 16 | ABA=0 | SCAB1 KO | no closure | Closure | OFF | 21719691 |
| 17 | ABA=1 | SCAB1 KO | slower closure | Closure | Some/ON | 21719691 |
| 18 | ABA=1 | SCAB1 KO | slower actin reorganization | Actin Reorganization | Some/ON | 21719691 |
| 19 | ABA=0 | *ROS CA*, SCAB1 KO | some closure | Closure | Some | 21719691 |
| 20 | ABA=0 | *CaIM CA*, SCAB1 KO | some closure | Closure | Some | 21719691 |
| 21 | ABA=0 | GEF1/4/10 KO | no closure | Closure | OFF | 22908257 |
| 22 | ABA=1 | GEF1/4/10 KO | increased closure | Closure | ON | 22908257,<br>22500990 |
| 23 | ABA=1 | GEF1/4/10 CA | reduced closure, closure | Closure | Some/ON | 22500990 |
| 24 | ABA=0 | SPHK1/2 KO | no closure | Closure | OFF | 18557834 |
| 25 | ABA=1 | SPHK1/2 KO | reduced closure | Closure | Some | 18557834 |
| 26 | ABA=0 | SPHK1/2 CA | no closure | Closure | OFF | 18557834 |
| 27 | ABA=1 | SPHK1/2 CA | increased closure | Closure | ON | 18557834 |
| 28 | ABA=0 | ABI1 KO | no closure | Closure | OFF | 16798945,<br>16614222,<br>19690149 |
| 29 | ABA=1 | ABI1 KO | increased closure<br>/closure | Closure | ON | 16798945<br>/16614222,<br>19690149 |
| 30 | ABA=0 | HAB1 KO | no closure | Closure | OFF | 16798945 |

| | | | | | | |
|---|---|---|---|---|---|---|
| 31 | ABA=1 | HAB1 KO | increased closure | Closure | ON | 16798945 |
| 32 | ABA=0 | ABI1 KO, HAB1 KO | no closure | Closure | OFF | 16798945 |
| 33 | ABA=1 | ABI1 KO, HAB1 KO | increased closure | Closure | ON | 16798945 |
| 34 | ABA=0 | ABI1 KO, *ROS CA* | closure | Closure | ON | 19690149 |
| 35 | ABA=0 | ABI1 KO, *NO CA* | closure | Closure | ON | 19690149 |
| 36 | ABA=0 | ABI1 CA | no closure | Closure | OFF | 10488243, 16614222, 12446847, 19690149 |
| 37 | ABA=1 | ABI1 CA | no closure | Closure | OFF | 10488243, 16614222, 12446847, 19690149 |
| 38 | ABA=0 | ABI1 CA, *ROS CA* | no closure | Closure | OFF | 19690149 |
| 39 | ABA=0 | ABI2 KO | no closure | Closure | OFF | 11208021 |
| 40 | ABA=1 | ABI2 KO | closure | Closure | ON | 11208021 |
| 41 | ABA=0 | ABI2 CA | no closure | Closure | OFF | 11208021, 10488243, 12446847 |
| 42 | ABA=1 | ABI2 CA | no closure | Closure | OFF/Some | 11208021, 10488243, 12446847 |
| 43 | ABA=0 | RCARs CA | no closure | Closure | OFF | 25969135 |
| 44 | ABA=1 | RCARs CA | increased closure | Closure | ON | 25969135 |
| 45 | ABA=0 | RCARs KO | no closure | Closure | OFF | 19874541 |
| 46 | ABA=1 | RCARs KO | reduced closure | Closure | Some | 19874541 |
| 47 | ABA=0 | *CaIM CA*, RCARs KO | some closure, closure | Closure | Some/ON | 19874541 |
| 48 | ABA=0 | GCR1 KO | no closure | Closure | OFF | 15155892 |
| 49 | ABA=1 | GCR1 KO | increased closure | Closure | ON | 15155892 |
| 50 | ABA=0 | TCTP CA | no closure | Closure | OFF | 22610367 |
| 51 | ABA=1 | TCTP CA | increased closure | Closure | ON | 22610367 |
| 52 | ABA=0 | S1P/PhytoS1P KO | no closure | Closure | OFF | 11279499 |
| 53 | ABA=1 | S1P/PhytoS1P KO | no, reduced closure | Closure | OFF/Some | 11279499 |
| 54 | ABA=0 | OST1 KO | no closure | Closure | OFF | 24033256, 31179540, 15064385 |
| 55 | ABA=1 | OST1 KO | no closure | Closure | OFF | 24033256, 30361234, 31179540, 15064385 |
| 56 | ABA=0 | OST1 CA | no closure | Closure | OFF | 24033256 |
| 57 | ABA=1 | OST1 CA | increased closure | Closure | ON | 24033256 |
| 58 | ABA=0 | ERA1 KO | no, some closure | Closure | OFF/Some | 9765153 |
| 59 | ABA=1 | ERA1 KO | increased closure | Closure | ON | 9765153 |
| 60 | ABA=0 | ABH1 KO | no, some closure | Closure | OFF/Some | 11525733 |
| 61 | ABA=1 | ABH1 KO | increased closure | Closure | ON | 11525733 |
| 62 | ABA=0 | PLDα KO | no closure | Closure | OFF | 16614222, 22932846, 19690149, Qu et al., 24271006, |

| | | | | | | |
|---|---|---|---|---|---|---|
| | | | | | | 19690149 |
| 63 | ABA=1 | PLDα KO | no closure<br>reduced closure<br>reduced closure | Closure | OFF/Some | 16614222, 19690149, Qu et al., 19690149, 24271006, 22392280 |
| 64 | ABA=0 | PLDα KO, *ROS CA* | closure | Closure | ON | 19690149, 22392280 |
| 65 | ABA=0 | PLDδ KO | no closure | Closure | OFF | 22589465, 22932846, 22392280 |
| 66 | ABA=1 | PLDδ KO | no closure<br>reduced closure, closure | Closure | OFF | 22589465, 22392280 |
| 67 | ABA=0 | PLDδ KO, *ROS CA* | no closure<br>closure | Closure | OFF | 22589465, 22392280 |
| 68 | ABA=0 | *PA CA* | some closure, closure | Closure | Some/ON | 10518598, 16614222, 24271006, Qu et al., 22392080 |
| 69 | ABA=1 | *PA CA* | closure | Closure | ON | 10518598 |
| 70 | ABA=0 | PLDα KO, PLDδ KO | no closure | Closure | OFF | 10518598, 22932846, 22392280 |
| 71 | ABA=1 | PLDα KO, PLDδ KO | reduced closure | Closure | Some | 10518598, 22392280 |
| 72 | ABA=0 | PLDα KO, PLDδ KO, *ROS CA* | some closure, closure | Closure | Some/ON | 22392280 |
| 73 | ABA=0 | *PA CA*, PLDα KO, PLDδ KO | closure (more than PA CA)<br>some closure (same as PA CA) | Closure | Some/ON | 10518598, 22392080 |
| 74 | ABA=0 | DAG CA | no closure | Closure | OFF | 10518598 |
| 75 | ABA=0 | ABI1 KO, *PA CA* | some closure, closure | Closure | Some/ON | 16614222 |
| 76 | ABA=0 | ABI1 CA, *PA CA* | no closure | Closure | OFF | 16614222 |
| 77 | ABA=0 | *PA CA*, PLDα KO | some closure, closure | Closure | Some/ON | 16614222, 24271006, 22392080 |
| 78 | ABA=0 | *PA CA*, PLDδ KO | some closure, closure | Closure | Some/ON | 22392080 |
| 79 | ABA=0 | GPA1 KO, *PA CA* | some closure, closure | Closure | Some/ON | 16614222 |
| 80 | ABA=0 | GPA1 KO, *PA CA*, PLDα KO | some closure, closure | Closure | Some/ON | 16614222 |
| 81 | ABA=0 | ABI1 KO, PLDα KO | no closure | Closure | OFF | 16614222 |
| 82 | ABA=1 | ABI1 KO, PLDα KO | closure | Closure | ON | 16614222 |
| 83 | ABA=0 | GPA1 KO | no closure | Closure | OFF | 16614222, 12789341 |
| 84 | ABA=1 | GPA1 KO | closure | Closure | ON | 16614222, 11408655, 12789341 |
| 85 | ABA=0 | GPA1 KO, PLDα KO | no closure | Closure | OFF | 16614222 |
| 86 | ABA=1 | GPA1 KO, PLDα KO | no closure | Closure | OFF | 16614222 |
| 87 | ABA=0 | PP2CA KO | no closure | Closure | OFF | Qu et al., 16361522 |
| 88 | ABA=0 | *PA CA*, PP2CA KO | some closure, closure (less than PA CA) | Closure | Some/ON | Qu et al. |
| 89 | ABA=0 | $Ca^{2+}_c$ KO, PLDα KO | no closure | Closure | OFF | 24271006 |

| | | | | | | |
|---|---|---|---|---|---|---|
| 90 | ABA=1 | $Ca^{2+}_c$ KO, PLDα KO | no, reduced closure | Closure | OFF/Some | 24271006 |
| 91 | ABA=0 | *PA CA*, $Ca^{2+}_c$ KO | some closure, closure | Closure | Some/ON | 24271006 |
| 92 | ABA=0 | *PA CA*, $Ca^{2+}_c$ KO, PLDα KO | some closure, closure | Closure | Some/ON | 24271006 |
| 93 | ABA=0 | CPK3/21 KO | no closure | Closure | OFF | 17032064, 21994053 |
| 94 | ABA=1 | CPK3/21 KO | reduced closure, closure | Closure | Some/ON | 17032064 |
| 95 | ABA=0 | CPK6 KO | no closure | Closure | OFF | 17032064, 21994053 |
| 96 | ABA=1 | CPK6 KO | reduced closure, closure | Closure | Some/ON | 17032064 |
| 97 | ABA=0 | CPK3/21 KO, CPK6 KO | no closure | Closure | OFF | 17032064 |
| 98 | ABA=1 | CPK3/21 KO, CPK6 KO | no, reduced closure | Closure | OFF/Some | 17032064 |
| 99 | ABA=0 | cGMP CA | no closure | Closure | OFF | 23396828 |
| 100 | ABA=1 | NOGC1 KO, cGMP CA | reduced closure, closure | Closure | Some/ON | 23396828 |
| 101 | ABA=0 | cADPR KO | no closure<br>no, some closure<br>/some closure | Closure | OFF/Some | 9861057,<br>10518598<br>/19847112 |
| 102 | ABA=1 | cADPR KO | slower closure<br>/reduced closure | Closure | Some/ON | 9861057<br>/19847112,<br>10518598 |
| 103 | ABA=0 | ADPRc KO | no closure | Closure | OFF | 9861057 |
| 104 | ABA=1 | ADPRc KO | no, reduced closure | Closure | OFF/Some | 9861057 |
| 105 | ABA=0 | PI3P5K KO | no closure | Closure | OFF | 23757398 |
| 106 | ABA=1 | PI3P5K KO | reduced, slower closure | Closure | Some/ON | 23757398 |
| 107 | ABA=0 | V-ATPase KO | no closure | Closure | OFF | 23757398 |
| 108 | ABA=1 | V-ATPase KO | reduced, slower closure | Closure | Some/ON | 23757398 |
| 109 | ABA=0 | V-PPase KO | no closure | Closure | OFF | 23757398 |
| 110 | ABA=1 | V-PPase KO | reduced, slower closure | Closure | Some/ON | 23757398 |
| 111 | ABA=0 | Vacuolar Acidification KO | no closure | Closure | OFF | 23757398 |
| 112 | ABA=1 | Vacuolar Acidification KO | reduced closure | Closure | Some | 23757398 |
| 113 | ABA=0 | PLC KO | no closure<br><br><br>/some closure | Closure | OFF/Some | 9990101,<br>17996010,<br>10518598<br>/19847112 |
| 114 | ABA=1 | PLC KO | reduced closure<br><br>/closure | Closure | Some/ON | 9990101,<br>10518598<br>/19847112 |
| 115 | ABA=0 | InsP3 KO | no closure | Closure | OFF | 9990101 |
| 116 | ABA=1 | InsP3 KO | no, reduced closure | Closure | OFF/Some | 9990101 |
| 117 | ABA=1 | PLDα KO, PLDδ KO, cADPR KO | no closure | Closure | OFF | 10518598 |
| 118 | ABA=1 | PLC KO, PLDα KO, PLDδ KO | no, reduced closure | Closure | OFF/Some | 10518598 |
| 119 | ABA=1 | PLC KO, cADPR KO | no closure | Closure | OFF | 10518598 |
| 120 | ABA=1 | PLC KO, PLDα KO, PLDδ KO, cADPR KO | no closure | Closure | OFF | 10518598 |
| 121 | ABA=0 | $pH_c$ KO | no closure | Closure | OFF | 16968132,<br>18721267,<br>15064385 |
| 122 | ABA=1 | $pH_c$ KO | reduced closure<br><br>no, reduced closure<br>/no closure | Closure | OFF/Some | 11408655,<br>18721267,<br>15064385<br>/16968132 |

| | | | | | | |
|---|---|---|---|---|---|---|
| 123 | ABA=0 | OST1 KO, $pH_c$ KO | no closure | Closure | OFF | 15064385 |
| 124 | ABA=1 | OST1 KO, $pH_c$ KO | no closure | Closure | OFF | 15064385 |
| 125 | ABA=1 | GPA1 KO, $pH_c$ KO | no, reduced closure | Closure | OFF/Some | 11408655 |
| 126 | ABA=0 | *NO CA*, RBOH KO | closure | Closure | ON | 16367958 |
| 127 | ABA=0 | PtdInsP4 KO | no closure | Closure | OFF | 12368494 |
| 128 | ABA=1 | PtdInsP4 KO | no, reduced closure | Closure | OFF/Some | 12368494 |
| 129 | ABA=0 | PtdInsP3 KO | no closure | Closure | OFF | 12368494 |
| 130 | ABA=1 | PtdInsP3 KO | no, reduced closure | Closure | OFF/Some | 12368494 |
| 131 | ABA=1 | PtdInsP3 KO, PtdInsP4 KO | no, reduced closure | Closure | OFF/Some | 12368494 17160388 |
| 132 | ABA=0 | ROP11 CA | no closure | Closure | OFF | 22233300 |
| 133 | ABA=1 | ROP11 CA | reduced closure | Closure | Some | 22233300 |
| 134 | ABA=0 | ROP11 KO | no closure | Closure | OFF | 22233300 |
| 135 | ABA=1 | ROP11 KO | increased closure | Closure | ON | 22233300 |
| 136 | ABA=0 | KOUT KO | no closure | Closure | OFF | 12671068, 15064385 |
| 137 | ABA=1 | KOUT KO | reduced closure | Closure | Some | 12671068, 15064385 |
| 138 | ABA=0 | $K^+$ efflux KO | no closure | Closure | OFF | 12671068 |
| 139 | ABA=1 | $K^+$ efflux KO | reduced closure | Closure | Some | 12671068 |
| 140 | ABA=0 | KOUT KO | no ROS | ROS | OFF | 15064385 |
| 141 | ABA=1 | KOUT KO | ROS | ROS | ON | 15064385 |
| 142 | ABA=0 | KOUT KO, $pH_c$ KO | no closure | Closure | OFF | 15064385 |
| 143 | ABA=1 | KOUT KO, $pH_c$ KO | reduced closure | Closure | Some | 15064385 |
| 144 | ABA=0 | PP2CA CA | no closure | Closure | OFF | 16361522 |
| 145 | ABA=1 | PP2CA CA | reduced closure | Closure | Some | 16361522 |
| 146 | ABA=1 | PP2CA KO | increased closure | Closure | ON | 16361522, Qu et al. |
| 147 | ABA=0 | $H^+$ ATPase CA | no closure | Closure | OFF | 24367097 |
| 148 | ABA=1 | $H^+$ ATPase CA | reduced closure | Closure | Some | 24367097 |
| 149 | ABA=0 | *AtRAC1 KO* | some closure, closure | Closure | Some/ON | 11459830 |
| 150 | ABA=0 | AtRAC1 CA | no closure | Closure | OFF | 11459830 |
| 151 | ABA=1 | AtRAC1 CA | reduced closure | Closure | Some | 11459830 |
| 152 | ABA=0 | QUAC1 KO | no closure | Closure | OFF | 20626656 |
| 153 | ABA=1 | QUAC1 KO | no, reduced closure | Closure | OFF/Some | 20626656 |
| 154 | ABA=0 | *CalM CA*, QUAC1 KO | no closure | Closure | OFF | 20154005 |
| 155 | ABA=0 | GAPC1/2 KO | no closure | Closure | OFF | 22589465 |
| 156 | ABA=1 | GAPC1/2 KO | reduced closure | Closure | Some | 22589465 |
| 157 | ABA=0 | GAPC1/2 KO, *ROS CA* | some closure | Closure | Some | 22589465 |
| 158 | ABA=0 | GAPC1/2 KO, PLDδ KO | no closure | Closure | OFF | 22589465 |
| 159 | ABA=1 | GAPC1/2 KO, PLDδ KO | no, reduced closure | Closure | OFF/Some | 22589465 |
| 160 | ABA=0 | GAPC1/2 KO, PLDδ KO, *ROS CA* | no, some closure | Closure | OFF/Some | 22589465 |
| 161 | ABA=0 | MPK9/12 KO | no closure | Closure | OFF | 19910530 |
| 162 | ABA=1 | MPK9/12 KO | no closure | Closure | OFF/Some | 19910530 |
| 163 | ABA=0 | MPK9/12 KO, *ROS CA* | some closure | Closure | Some | 19910530 |
| 164 | ABA=0 | SLAC1 KO | no closure | Closure | OFF | 18305484 |

| | | | | | | |
|---|---|---|---|---|---|---|
| 165 | ABA=1 | SLAC1 KO | no closure | Closure | OFF | 18305484 |
| 166 | ABA=0 | ARP Complex KO | no, some closure | Closure | OFF/Some | 22570440 |
| 167 | ABA=1 | ARP Complex KO | no, reduced closure | Closure | OFF/Some | 22570440 |
| 168 | ABA=0 | ARP Complex KO, *CaIM CA* | some closure | Closure | Some | 22570440 |
| 169 | ABA=0 | ARP Complex KO | no actin reorganization | Actin Reorganization | OFF | 22570440 |
| 170 | ABA=1 | ARP Complex KO | no, reduced actin reorganization | Actin Reorganization | OFF/Some | 22570440 |
| 171 | ABA=0 | Aquaporin(PIP2;1) KO | no closure | Closure | OFF | 26163575 |
| 172 | ABA=1 | Aquaporin(PIP2;1) KO | reduced closure | Closure | Some | 26163575 |
| 173 | ABA=0 | GHR1 KO | no closure | Closure | OFF | 22730405, 30361234 |
| 174 | ABA=1 | GHR1 KO | no closure | Closure | OFF | 22730405, 30361234 |
| 175 | ABA=0 | GHR1 KO, *ROS CA* | no closure | Closure | OFF | 30361234, 22730405 |
| 176 | ABA=0 | OST1 KO, *ROS CA* | some closure /closure (same as ROS CA) | Closure | Some/ON | 30361234 /22730405 |
| 177 | ABA=0 | *CaIM CA*, GHR1 KO | closure | Closure | ON | 22730405 |
| 178 | ABA=0 | RCN1 KO | no, some closure | Closure | OFF/Some | 12417706 |
| 179 | ABA=1 | RCN1 KO | reduced closure | Closure | Some | 12417706 |
| 180 | ABA=0 | SPP1 KO | no closure | Closure | OFF | 21910031 |
| 181 | ABA=1 | SPP1 KO | increased closure | Closure | ON | 21910031 |
| 182 | ABA=0 | NIA1/2 KO | no, some closure | Closure | OFF/Some | 12446847 |
| 183 | ABA=1 | NIA1/2 KO | no closure | Closure | OFF | 12446847, 16367958 |
| 184 | ABA=0 | Microtubule Depolymerization KO | no closure | Closure | OFF | 24271006, Qu et al. |
| 185 | ABA=1 | Microtubule Depolymerization KO | no, reduced closure | Closure | OFF/Some | 24271006, Qu et al. |
| 186 | ABA=0 | Microtubule Depolymerization CA | no closure | Closure | OFF | 24271006, Qu et al. |
| 187 | ABA=1 | Microtubule Depolymerization CA | closure | Closure | ON | 24271006, Qu et al. |
| 188 | ABA=0 | Microtubule Depolymerization CA, PLDα KO | no closure | Closure | OFF | 24271006, Qu et al. |
| 189 | ABA=1 | Microtubule Depolymerization CA, PLDα KO | reduced closure, closure | Closure | Some/ON | 24271006, Qu et al. |
| 190 | ABA=0 | Microtubule Depolymerization CA, PP2CA KO | no closure | Closure | OFF | Qu et al. |
| 191 | ABA=1 | Microtubule Depolymerization CA, PP2CA KO | increased closure | Closure | ON | Qu et al. |
| 192 | ABA=0 | Microtubule Depolymerization KO, PLDα KO | no closure | Closure | OFF | Qu et al. |
| 193 | ABA=1 | Microtubule Depolymerization KO, PLDα KO | no closure | Closure | OFF | Qu et al. |
| 194 | ABA=0 | Microtubule Depolymerization KO, PP2CA KO | no closure | Closure | OFF | Qu et al. |
| 195 | ABA=1 | Microtubule Depolymerization KO, PP2CA KO | closure | Closure | ON | Qu et al. |
| 196 | ABA=0 | NOGC1 KO | no closure | Closure | OFF | 23396828 |

| | | | | | | |
|---|---|---|---|---|---|---|
| 197 | ABA=1 | NOGC1 KO | no closure | Closure | OFF | 23396828 |
| 198 | ABA=0 | *NO CA*, NOGC1 KO | no closure | Closure | OFF | 23396828 |
| 199 | ABA=0 | MRP5 KO | no closure | Closure | OFF | 17098742 |
| 200 | ABA=1 | MRP5 KO | reduced closure | Closure | Some | 17098742 |
| 201 | ABA=0 | *$pH_c$ CA* | some closure, closure(>50%) | Closure | Some/ON | 18721267 |
| 202 | ABA=1 | *$pH_c$ CA* | increased closure | Closure | ON | 18721267 |
| 203 | ABA=0 | NO KO | no closure | Closure | OFF | 18721267 |
| 204 | ABA=1 | NO KO | no closure | Closure | OFF | 18721267,<br>16367958 |
| 205 | ABA=1 | SLAH3 KO | closure | Closure | ON | 21586729 |
| 206 | ABA=0 | NtSyp121 KO | no closure | Closure | OFF | 19825544 |
| 207 | ABA=1 | NtSyp121 KO | no closure | Closure | OFF | 19825544 |
| 208 | ABA=0 | *S1P/PhytoS1P CA* | some closure, closure(>50%) | Closure | Some/ON | 12789341 |
| 209 | ABA=0 | GPA1 KO, *S1P/PhytoS1P CA* | no, some closure | Closure | OFF/Some | 12789341 |
| 210 | ABA=0 | Nitrite CA | some closure, closure(>50%) | Closure | Some/ON | 12446847 |
| 211 | ABA=0 | Nitrite CA, NIA1/2 KO | no closure | Closure | OFF | 12446847 |
| 212 | ABA=0 | *NO CA* | some closure, closure(>50%)<br>/some closure(<25%) | Closure | Some/ON | 12446847,<br>19825544,<br>17996010,<br>18721267,<br>16367958,<br>19690149<br>/23396828,<br>22932846 |
| 213 | ABA=0 | NIA1/2 KO, *NO CA* | some closure, closure(>50%) | Closure | Some/ON | 12446847,<br>16367958 |
| 214 | ABA=0 | NIA1/2 KO, *ROS CA* | closure(>50%)<br>/no, some closure* | Closure | Some/ON | 12446847<br>/16367958 |
| 215 | ABA=0 | *NO CA*, NOGC1 KO, cGMP CA | some closure, closure (similar to NO CA) | Closure | Some/ON | 23396828 |
| 216 | ABA=0 | GPA1 KO, *ROS CA* | closure (same as ROS CA) | Closure | ON | 21262908 |
| 217 | ABA=0 | Actin Reorganization KO, *ROS CA* | some closure | Closure | Some | 24372484 |
| 218 | ABA=0 | *8-nitro-cGMP CA* | some closure(~25%) | Closure | Some | 23396828 |
| 219 | ABA=0 | *8-nitro-cGMP CA*, SLAC1 KO | no closure | Closure | OFF | 23396828 |
| 220 | ABA=0 | *cADPR CA* | closure(>50%) | Closure | ON | 9861057 |
| 221 | ABA=0 | ABI1 CA, *AtRAC1 KO* | some closure, closure | Closure | Some/ON | 11459830 |
| 222 | ABA=0 | ABI1 CA, AtRAC1 CA | no closure | Closure | OFF | 11459830 |
| 223 | ABA=0 | *InsP3 CA* | some closure, closure(~25%) | Closure | Some/ON | 2388697 |
| 224 | ABA=0 | *InsP3 CA* | $Ca^{2+}_c$ oscillation | $Ca^{2+}_c$ | Some | 2388697 |
| 225 | ABA=0 | CalM KO | no closure | Closure | OFF | 2388697 |
| 226 | ABA=0 | CalM KO | no $Ca^{2+}_c$ | $Ca^{2+}_c$ | OFF | 2388697 |
| 227 | ABA=0 | CalM KO, *InsP3 CA* | some closure, closure(~25%) | Closure | Some/ON | 2388697 |
| 228 | ABA=0 | CalM KO, *InsP3 CA* | $Ca^{2+}_c$ oscillation | $Ca^{2+}_c$ | Some | 2388697 |
| 229 | ABA=0 | *NO CA*, NtSyp121 KO | some closure | Closure | Some | 19825544 |
| 230 | ABA=0 | *NO CA*, PLC KO | no closure | Closure | OFF | 17996010 |
| 231 | ABA=0 | *NO CA*, PLDδ KO | no closure | Closure | OFF | 22932846 |
| 232 | ABA=0 | *NO CA*, PLDα KO | some closure, closure (same as NO CA) | Closure | Some/ON | 22932846,<br>19690149 |

| | | | | | | |
|---|---|---|---|---|---|---|
| 233 | ABA=0 | *NO CA*, PLDα KO, PLDδ KO | no closure | Closure | OFF | 22932846 |
| 234 | ABA=0 | *8-nitro-cGMP CA*, cADPR KO | no closure | Closure | OFF | 23396828 |
| 235 | ABA=0 | *8-nitro-cGMP CA*, $Ca^{2+}_c$ KO | no, some closure | Closure | OFF/Some | 23396828 |
| 236 | ABA=0 | GCR1 KO, *S1P/PhytoS1P CA* | closure(>50%) | Closure | ON | 15155892 |
| 237 | ABA=0 | *CaIM CA*, CPK3/21 KO | some closure | Closure | Some | 21994053 |
| 238 | ABA=0 | *CaIM CA*, CPK6 KO | some closure | Closure | Some | 21994053 |
| 239 | ABA=0 | ABI1 CA, *NO CA* | no closure | Closure | OFF | 12446847, 19690149 |
| 240 | ABA=0 | ABI2 CA, *NO CA* | no closure | Closure | OFF | 12446847 |
| 241 | ABA=0 | *CaIM CA*, TCTP CA | increased closure compared to CaIM CA | Closure | ON | 22610367 |
| 242 | ABA=0 | *CaIM CA*, PLDα KO | no closure | Closure | OFF | 24271006 |
| 243 | ABA=0 | NO KO, *ROS CA* | no, some closure | Closure | OFF/Some | 16367958 |
| 244 | ABA=0 | *NO CA*, ROS KO | closure | Closure | ON | 16367958 |
| 245 | ABA=0 | WT | no ROS | ROS | OFF | 23946352, 16367958, 23396828, 15064385, 24033256, 12746515, 19690149, 22730405 |
| 246 | ABA=1 | WT | ROS | ROS | ON | 23946352, 16367958, 23396828, 15064385, 24033256, 12746515, 19690149, 22730405, 22392280 |
| 247 | ABA=0 | WT | no NO | NO | OFF | 23946352, 18721267, 16367958, 23396828, 19690149 |
| 248 | ABA=1 | WT | NO | NO | ON | 23946352, 18721267, 16367958, 23396828, 19690149, 22392280 |
| 249 | ABA=0 | WT | no $pH_c$ | $pH_c$ | OFF | 23946352, 18721267, 20739306, 15064385, 22392280 |
| 250 | ABA=1 | WT | $pH_c$ | $pH_c$ | ON | 23946352, 18721267, 20739306, 15064385, 22392280 |
| 251 | ABA=0 | RCARs KO | no ROS | ROS | OFF | 23946352 |
| 252 | ABA=1 | RCARs KO | no ROS | ROS | OFF | 23946352 |
| 253 | ABA=0 | RCARs KO | no NO | NO | OFF | 23946352 |
| 254 | ABA=1 | RCARs KO | no NO | NO | OFF | 23946352 |

| | | | | | | |
|---|---|---|---|---|---|---|
| 255 | ABA=0 | RCARs KO | no $pH_c$ | $pH_c$ | OFF | 23946352 |
| 256 | ABA=1 | RCARs KO | no $pH_c$ | $pH_c$ | OFF | 23946352 |
| 257 | ABA=0 | NO CA | some $pH_c$ | $pH_c$ | Some | 18721267 |
| 258 | ABA=1 | NO CA | $pH_c$ | $pH_c$ | ON | 18721267 |
| 259 | ABA=0 | NO KO | no $pH_c$ | $pH_c$ | OFF | 18721267 |
| 260 | ABA=1 | NO KO | reduced $pH_c$ | $pH_c$ | Some | 18721267 |
| 261 | ABA=0 | $Ca^{2+}_c$ KO | no $pH_c$ | $pH_c$ | OFF | 18721267, 15064385 |
| 262 | ABA=1 | $Ca^{2+}_c$ KO | no $pH_c$ /$pH_c$ | $pH_c$ | OFF | 18721267 /15064385 |
| 263 | ABA=0 | $Ca^{2+}_c$ KO | no ROS | ROS | OFF | 15064385 |
| 264 | ABA=1 | $Ca^{2+}_c$ KO | ROS | ROS | ON | 15064385 |
| 265 | ABA=0 | $pH_c$ KO | no NO | NO | OFF | 18721267 |
| 266 | ABA=1 | $pH_c$ KO | no NO | NO | OFF | 18721267 |
| 267 | ABA=0 | $pH_c$ CA | NO | NO | ON | 18721267 |
| 268 | ABA=1 | $pH_c$ CA | NO | NO | ON | 18721267 |
| 269 | ABA=0 | $Ca^{2+}_c$ KO | no NO | NO | OFF | 18721267 |
| 270 | ABA=1 | $Ca^{2+}_c$ KO | no NO | NO | OFF | 18721267 |
| 271 | ABA=0 | ROS CA | NO | NO | ON | 16367958, 19690149 |
| 272 | ABA=0 | NIA1/2 KO | no, some NO | NO | OFF/Some | 16367958 |
| 273 | ABA=1 | NIA1/2 KO | no NO | NO | OFF | 16367958 |
| 274 | ABA=0 | NIA1/2 KO, ROS CA | no NO | NO | OFF | 16367958 |
| 275 | ABA=0 | Nitrite CA | NO | NO | ON | 16367958 |
| 276 | ABA=0 | NIA1/2 KO, Nitrite CA | no NO | NO | OFF | 16367958 |
| 277 | ABA=0 | RBOH KO | no NO | NO | OFF | 16367958 |
| 278 | ABA=1 | RBOH KO | no NO | NO | OFF | 16367958 |
| 279 | ABA=0 | RBOH KO, ROS CA | NO | NO | ON | 16367958 |
| 280 | ABA=0 | ABI1 KO | no NO | NO | OFF | 19690149 |
| 281 | ABA=1 | ABI1 KO | NO | NO | ON | 19690149 |
| 282 | ABA=0 | NO CA | no ROS | ROS | OFF | 16367958 |
| 283 | ABA=1 | NO KO | ROS | ROS | ON | 16367958 |
| 284 | ABA=0 | WT | no CalM | CalM | OFF | 24033256 |
| 285 | ABA=1 | WT | CalM | CalM | ON | 24033256 |
| 286 | ABA=0 | WT | no SACC | SACC | OFF | 24033256 |
| 287 | ABA=0 | SACC KO | no CalM | CalM | OFF | 21262908 |
| 288 | ABA=1 | SACC KO | CalM | CalM | ON | 21262908 |
| 289 | ABA=0 | SACC KO, GPA1 KO | no CalM | CalM | OFF | 21262908 |
| 290 | ABA=1 | SACC KO, GPA1 KO | no CalM | CalM | OFF | 21262908 |
| 291 | ABA=0 | GPA1 KO | no ROS | ROS | OFF | 21262908 |
| 292 | ABA=1 | GPA1 KO | no ROS | ROS | OFF | 21262908 |
| 293 | ABA=0 | Aquaporin(PIP2;1) KO | no ROS | ROS | OFF | 26163575 |
| 294 | ABA=1 | Aquaporin(PIP2;1) KO | no ROS | ROS | OFF | 26163575, 28784763 |
| 295 | ABA=0 | WT | no Aquaporin(PIP2;1) | Aquaporin(PIP2;1) | OFF | 28784763 |
| 296 | ABA=1 | WT | Aquaporin(PIP2;1) | Aquaporin(PIP2;1) | ON | 28784763 |

| | | | | | | |
|---|---|---|---|---|---|---|
| 297 | ABA=0 | OST1 KO | no Aquaporin(PIP2;1) | Aquaporin(PIP2;1) | OFF | 28784763 |
| 298 | ABA=1 | OST1 KO | no Aquaporin(PIP2;1) | Aquaporin(PIP2;1) | OFF | 28784763 |
| 299 | ABA=0 | WT | no SLAC1 | SLAC1 | OFF | 22730405, 19910530 |
| 300 | ABA=1 | WT | SLAC1 | SLAC1 | ON | 22730405, 19910530 |
| 301 | ABA=0 | OST1 KO | no SLAC1 | SLAC1 | OFF | 22730405 |
| 302 | ABA=1 | OST1 KO | no SLAC1 | SLAC1 | OFF | 22730405 |
| 303 | ABA=0 | GHR1 KO | no SLAC1 | SLAC1 | OFF | 22730405 |
| 304 | ABA=1 | GHR1 KO | no, reduced SLAC1 | SLAC1 | OFF/Some | 22730405 |
| 305 | ABA=1 | ABI1 CA | no $pH_c$ | $pH_c$ | OFF | 20739306 |
| 306 | ABA=1 | ABI2 CA | no $pH_c$ | $pH_c$ | OFF | 20739306 |
| 307 | ABA=1 | OST1 KO | no $pH_c$ | $pH_c$ | OFF | 20739306 |
| 308 | ABA=0 | CalM CA | $pH_c$ | $pH_c$ | ON | 20739306 |
| 309 | ABA=0 | CalM CA, ABI1 CA | $pH_c$ | $pH_c$ | ON | 20739306 |
| 310 | ABA=0 | CalM CA, ABI2 CA | $pH_c$ | $pH_c$ | ON | 20739306 |
| 311 | ABA=0 | CalM CA, OST1 KO | $pH_c$ | $pH_c$ | ON | 20739306 |
| 312 | ABA=0 | PLDα KO, PLDδ KO | no $pH_c$ | $pH_c$ | OFF | 22392280 |
| 313 | ABA=1 | PLDα KO, PLDδ KO | no $pH_c$ | $pH_c$ | OFF | 22392280 |
| 314 | ABA=0 | WT | no $Ca^{2+}_c$ | $Ca^{2+}_c$ | OFF | 20739306, 31179540, 22392280 |
| 315 | ABA=1 | WT | $Ca^{2+}_c$ oscillation | $Ca^{2+}_c$ | Some | 20739306, 31179540, 22392280 |
| 316 | ABA=1 | $pH_c$ KO | no, reduced $Ca^{2+}_c$ | $Ca^{2+}_c$ | OFF/Some | 20739306 |
| 317 | ABA=0 | OST1 KO | no $Ca^{2+}_c$ | $Ca^{2+}_c$ | OFF | 31179540 |
| 318 | ABA=1 | OST1 KO | no $Ca^{2+}_c$ | $Ca^{2+}_c$ | OFF | 31179540 |
| 319 | ABA=0 | PLDα KO, PLDδ KO | no $Ca^{2+}_c$ | $Ca^{2+}_c$ | OFF | 22392280 |
| 320 | ABA=1 | PLDα KO, PLDδ KO | $Ca^{2+}_c$ oscillation | $Ca^{2+}_c$ | Some | 22392280 |
| 321 | ABA=0 | WT | no actin reorganization | Actin Reorganization | OFF | 11459830, 18088331 |
| 322 | ABA=1 | WT | actin reorganization | Actin Reorganization | ON | 11459830, 18088331 |
| 323 | ABA=0 | ABI1 CA | no actin reorganization | Actin Reorganization | OFF | 11459830 |
| 324 | ABA=1 | ABI1 CA | no actin reorganization | Actin Reorganization | OFF | 11459830 |
| 325 | ABA=0 | PtdInsP3 KO | no actin reorganization | Actin Reorganization | OFF | 18088331 |
| 326 | ABA=1 | PtdInsP3 KO | no, reduced actin reorganization | Actin Reorganization | OFF/Some | 18088331 |
| 327 | ABA=0 | PtdInsP4 KO | no actin reorganization | Actin Reorganization | OFF | 18088331 |
| 328 | ABA=1 | PtdInsP4 KO | no, reduced actin reorganization | Actin Reorganization | OFF/Some | 18088331 |
| 329 | ABA=1 | PtdInsP3 KO, PtdInsP4 KO | no, reduced actin reorganization | Actin Reorganization | OFF/Some | 18088331 |
| 330 | ABA=1 | PLC KO | actin reorganization | Actin Reorganization | ON | 18088331 |
| 331 | ABA=1 | ROS KO | no, reduced actin reorganization | Actin Reorganization | OFF/Some | 18088331 |
| 332 | ABA=0 | RBOH KO | no actin reorganization | Actin Reorganization | OFF | 24372484 |

| | | | | | | |
|---|---|---|---|---|---|---|
| 333 | ABA=1 | RBOH KO | no actin reorganization | Actin Reorganization | OFF/Some | 24372484 |
| 334 | ABA=0 | WT | AtRAC1 | AtRAC1 | ON | 11459830 |
| 335 | ABA=1 | WT | no AtRAC1 | AtRAC1 | OFF | 11459830 |
| 336 | ABA=0 | ABI1 CA | AtRAC1 | AtRAC1 | ON | 11459830 |
| 337 | ABA=1 | ABI1 CA | AtRAC1 | AtRAC1 | ON | 11459830 |
| 338 | ABA=0 | WT | no 8-nitro-cGMP | 8-nitro-cGMP | OFF | 23396828 |
| 339 | ABA=1 | WT | 8-nitro-cGMP | 8-nitro-cGMP | ON | 23396828 |
| 340 | ABA=0 | NO CA | some 8-nitro-cGMP, 8-nitro-cGMP | 8-nitro-cGMP | Some/ON | 23396828 |
| 341 | ABA=1 | NO KO | no 8-nitro-cGMP | 8-nitro-cGMP | OFF | 23396828 |
| 342 | ABA=1 | NOGC1 KO | no 8-nitro-cGMP | 8-nitro-cGMP | OFF | 23396828 |
| 343 | ABA=1 | NOGC1 KO, cGMP CA | 8-nitro-cGMP | 8-nitro-cGMP | ON | 23396828 |
| 344 | ABA=0 | NO CA, NOGC1 KO | no 8-nitro-cGMP | 8-nitro-cGMP | OFF | 23396828 |
| 345 | ABA=0 | NO CA, NOGC1 KO, cGMP CA | 8-nitro-cGMP | 8-nitro-cGMP | ON | 23396828 |
| 346 | ABA=1 | ROS KO | no 8-nitro-cGMP | 8-nitro-cGMP | OFF | 23396828 |
| 347 | ABA=0 | WT | no $K^+$ efflux | $K^+$ efflux | OFF | 11027317 |
| 348 | ABA=1 | WT | $K^+$ efflux | $K^+$ efflux | ON | 11027317 |
| 349 | ABA=0 | WT | no KEV | KEV | OFF | 11027317 |
| 350 | ABA=1 | WT | KEV | KEV | ON | 11027317 |
| 351 | ABA=0 | CalM CA($Ca^{2+}_c$ oscillation) | some KEV, KEV | KEV | Some/ON | 11027317 |
| 352 | ABA=0 | WT | no microtubule depolymerization | Microtubule Depolymerization | OFF | 22402260, Qu et al. |
| 353 | ABA=1 | WT | microtubule depolymerization | Microtubule Depolymerization | ON | 22402260, Qu et al. |
| 354 | ABA=0 | PLDα KO | no microtubule depolymerization | Microtubule Depolymerization | OFF | Qu et al. |
| 355 | ABA=1 | PLDα KO | reduced microtubule depolymerization | Microtubule Depolymerization | Some | Qu et al. |
| 356 | ABA=0 | PP2CA KO | no microtubule depolymerization | Microtubule Depolymerization | OFF | Qu et al. |
| 357 | ABA=1 | PP2CA KO | microtubule depolymerization | Microtubule Depolymerization | ON | Qu et al. |
| 358 | ABA=0 | PA CA | microtubule depolymerization | Microtubule Depolymerization | ON | Qu et al. |
| 359 | ABA=0 | PA CA, PP2CA KO | some microtubule depolymerization, microtubule depolymerization | Microtubule Depolymerization | Some/ON | Qu et al. |
| 360 | ABA=0 | $Ca^{2+}_c$ KO | no microtubule depolymerization | Microtubule Depolymerization | OFF | 24271006 |
| 361 | ABA=1 | $Ca^{2+}_c$ KO | no, reduced microtubule depolymerization | Microtubule Depolymerization | OFF/Some | 24271006 |
| 362 | ABA=0 | $Ca^{2+}_c$ KO, PA CA | microtubule depolymerization | Microtubule Depolymerization | ON | 24271006 |
| 363 | ABA=0 | $Ca^{2+}_c$ KO, PLDα KO | no, some microtubule depolymerization | Microtubule Depolymerization | OFF/Some | 24271006 |
| 364 | ABA=1 | $Ca^{2+}_c$ KO, PLDα KO | no, reduced microtubule depolymerization | Microtubule Depolymerization | OFF/Some | 24271006 |
| 365 | ABA=0 | $Ca^{2+}_c$ KO, PA CA, PLDα KO | microtubule depolymerization | Microtubule Depolymerization | ON | 24271006 |
| 366 | ABA=0 | CalM CA | microtubule depolymerization | Microtubule | ON | 24271006 |

| | | | | | | |
|---|---|---|---|---|---|---|
| | | | | Depolymerization | | |
| 367 | ABA=0 | CalM CA, PLDα KO | some microtubule depolymerization | Microtubule Depolymerization | Some | 24271006 |
| 368 | ABA=0 | WT | no vacuolar acidification | Vacuolar Acidification | OFF | 23757398 |
| 369 | ABA=1 | WT | vacuolar acidification | Vacuolar Acidification | ON | 23757398 |
| 370 | ABA=0 | PI3P5K KO | no vacuolar acidification | Vacuolar Acidification | OFF | 23757398 |
| 371 | ABA=1 | PI3P5K KO | no, reduced vacuolar acidification | Vacuolar Acidification | OFF/Some | 23757398 |
| 372 | ABA=0 | PI3P5K KO | no $pH_c$ | $pH_c$ | OFF | 23757398 |
| 373 | ABA=1 | PI3P5K KO | no, reduced $pH_c$ | $pH_c$ | OFF/Some | 23757398 |
| 374 | ABA=0 | V-PPase KO | no vacuolar acidification | Vacuolar Acidification | OFF | 23757398 |
| 375 | ABA=1 | V-PPase KO | reduced vacuolar acidification | Vacuolar Acidification | Some | 23757398 |
| 376 | ABA=0 | $pH_c$ KO | no vacuolar acidification | Vacuolar Acidification | OFF | 23757398 |
| 377 | ABA=1 | $pH_c$ KO | no, reduced vacuolar acidification | Vacuolar Acidification | OFF/Some | 23757398 |
| 378 | ABA=0 | $pH_c$ KO | no ROS | ROS | OFF | 15064385 |
| 379 | ABA=1 | $pH_c$ KO | reduced ROS* | ROS | Some | 15064385 |
| 380 | ABA=0 | OST1 KO | no CalM | CalM | OFF | 24033256 |
| 381 | ABA=1 | OST1 KO | no CalM | CalM | OFF | 24033256 |
| 382 | ABA=0 | OST1 CA | no CalM | CalM | OFF | 24033256 |
| 383 | ABA=1 | OST1 CA | increased CalM | CalM | ON | 24033256 |
| 384 | ABA=0 | OST1 KO | no ROS | ROS | OFF | 24033256, 15064385 |
| 385 | ABA=1 | OST1 KO | no ROS | ROS | OFF | 24033256, 15064385 |
| 386 | ABA=0 | OST1 CA | no ROS | ROS | OFF | 24033256 |
| 387 | ABA=1 | OST1 CA | increased ROS | ROS | ON | 24033256 |
| 388 | ABA=0 | RBOH KO | no ROS | ROS | OFF | 15064385 |
| 389 | ABA=1 | RBOH KO | no ROS | ROS | OFF | 15064385 |
| 390 | ABA=0 | OST1 KO, RBOH KO | no ROS | ROS | OFF | 15064385 |
| 391 | ABA=1 | OST1 KO, RBOH KO | no ROS | ROS | OFF | 15064385 |
| 392 | ABA=0 | KOUT KO, RBOH KO | no ROS | ROS | OFF | 15064385 |
| 393 | ABA=1 | KOUT KO, RBOH KO | no ROS | ROS | OFF | 15064385 |
| 394 | ABA=0 | ABI1 CA | no ROS | ROS | OFF | 11701885, 19690149 |
| 395 | ABA=1 | ABI1 CA | no ROS /ROS | ROS | OFF | 11701885 /19690149 |
| 396 | ABA=0 | ABI2 CA | no ROS | ROS | OFF | 11701885 |
| 397 | ABA=1 | ABI2 CA | ROS | ROS | ON | 11701885 |
| 398 | ABA=0 | ABI1 KO | no ROS | ROS | OFF | 19690149 |
| 399 | ABA=1 | ABI1 KO | ROS | ROS | ON | 19690149 |
| 400 | ABA=0 | PtdInsP3 KO | no ROS | ROS | OFF | 12746515 |
| 401 | ABA=1 | PtdInsP3 KO | no, reduced ROS | ROS | OFF/Some | 12746515 |

| | | | | | | |
|---|---|---|---|---|---|---|
| 402 | ABA=0 | PtdInsP4 KO | no ROS | ROS | OFF | 12746515 |
| 403 | ABA=1 | PtdInsP4 KO | ROS | ROS | ON | 12746515 |
| 404 | ABA=1 | PtdInsP3 KO, PtdInsP4 KO | no ROS | ROS | OFF | 12746515 |
| 405 | ABA=0 | PLDδ KO | no NO | NO | OFF | 22932846 |
| 406 | ABA=1 | PLDδ KO | NO | NO | ON | 22932846 |
| 407 | ABA=0 | PLDδ KO | no ROS | ROS | OFF | 22932846 |
| 408 | ABA=1 | PLDδ KO | ROS<br>reduced ROS, ROS | ROS | Some/ON | 22932846, 22392280 |
| 409 | ABA=0 | WT | no PA | PA | OFF | 19690149 |
| 410 | ABA=1 | WT | PA | PA | ON | 19690149, 10518598, 22392280 |
| 411 | ABA=0 | PLDα KO | no PA | PA | OFF | 19690149 |
| 412 | ABA=1 | PLDα KO | reduced PA | PA | Some | 19690149, 22392280 |
| 413 | ABA=0 | PLDα KO | no, some ROS | ROS | OFF/Some | 19690149 |
| 414 | ABA=1 | PLDα KO | no, reduced ROS<br>reduced ROS | ROS | Some | 19690149, 22392280 |
| 415 | ABA=1 | PLDα KO, PLDδ KO | no, reduced ROS | ROS | OFF/Some | 22392280 |
| 416 | ABA=0 | PLDα KO | no NO | NO | OFF | 19690149 |
| 417 | ABA=1 | PLDα KO | no NO | NO | OFF | 19690149 |
| 418 | ABA=1 | PLDα KO, PLDδ KO | no NO | NO | OFF | 22392280 |
| 419 | ABA=0 | PLDα KO, ROS CA | NO | NO | ON | 19690149 |
| 420 | ABA=0 | WT | no AnionEM | AnionEM | OFF | 19847112 |
| 421 | ABA=1 | WT | AnionEM | AnionEM | ON | 19847112 |
| 422 | ABA=0 | cADPR KO | no AnionEM | AnionEM | OFF | 19847112 |
| 423 | ABA=1 | cADPR KO | no AnionEM | AnionEM | OFF | 19847112 |
| 424 | ABA=0 | PLC KO | no AnionEM | AnionEM | OFF | 19847112 |
| 425 | ABA=1 | PLC KO | no AnionEM | AnionEM | OFF | 19847112 |
| 426 | ABA=1 | WT | Depolarization | Depolarization | ON | 19847112 |
| 427 | ABA=1 | cADPR KO | no Depolarization | Depolarization | OFF | 19847112 |
| 428 | ABA=1 | PLC KO | no Depolarization | Depolarization | OFF | 19847112 |
| 429 | ABA=0 | SACC KO, ERA1 KO | no CaIM | CaIM | OFF | 12119381 |
| 430 | ABA=1 | SACC KO, ERA1 KO | increased CaIM | CaIM | ON | 12119381 |
| 431 | ABA=0 | ERA1 KO | no SLAC1 | SLAC1 | OFF | 12119381 |
| 432 | ABA=1 | ERA1 KO | SLAC1 | SLAC1 | ON | 12119381 |
| 433 | ABA=0 | $Ca^{2+}_c$ KO | no QUAC1 | QUAC1 | OFF | 23452338 |
| 434 | ABA=1 | $Ca^{2+}_c$ KO | QUAC1 | QUAC1 | ON | 23452338 |
| 435 | ABA=1 | ABI1 CA, $Ca^{2+}_c$ KO | no QUAC1 | QUAC1 | OFF | 23452338 |
| 436 | ABA=1 | $Ca^{2+}_c$ KO, OST1 KO | no QUAC1 | QUAC1 | OFF | 23452338 |
| 437 | ABA=1 | PLDα KO, PLDδ KO | no PA | PA | OFF | 10518598, 22392280 |
| 438 | ABA=1 | PLDδ KO | reduced PA | PA | Some | 22392280 |
| 439 | ABA=0 | WT | no PLDα | PLDα | OFF | 10518598 |
| 440 | ABA=1 | WT | reduced PLDα, PLDα | PLDα | Some/ON | 10518598 |
| 441 | ABA=0 | WT | no PLDδ | PLDδ | OFF | 10518598 |

| | | | | | | |
|---|---|---|---|---|---|---|
| 442 | ABA=1 | WT | reduced PLDδ, PLDδ | PLDδ | Some/ON | 10518598 |
| 443 | ABA=1 | WT | no, reduced DAG | DAG | OFF/Some | 10518598 |
| 444 | ABA=0 | ROS CA | some SLAC1, SLAC1 | SLAC1 | Some/ON | 22730405 |
| 445 | ABA=0 | OST1 KO, ROS CA | some SLAC1, SLAC1 | SLAC1 | Some/ON | 22730405 |
| 446 | ABA=0 | GHR1 KO, ROS CA | no, some SLAC1 | SLAC1 | OFF/Some | 22730405 |
| 447 | ABA=0 | SACC KO, ROS CA | some CalM, CalM | CalM | Some/ON | 21262908 |
| 448 | ABA=0 | SACC KO, GPA1 KO, ROS CA | some CalM, CalM | CalM | Some/ON | 21262908 |
| 449 | ABA=0 | S1P/PhytoS1P CA | $Ca^{2+}_c$ oscillation | $Ca^{2+}_c$ | Some | 11279499 |
| 450 | ABA=0 | CalM CA | $Ca^{2+}_c$ oscillation | $Ca^{2+}_c$ | Some | 11429606 |
| 451 | ABA=0 | AtRAC1 KO | some actin reorganization, actin reorganization | Actin Reorganization | Some/ON | 11459830 |
| 452 | ABA=0 | NO CA | some PA, PA | PA | Some/ON | 17996010 |
| 453 | ABA=0 | PLC KO | no PA | PA | OFF | 17996010 |
| 454 | ABA=0 | NO CA, PLC KO | no, some PA | PA | OFF/Some | 17996010 |
| 455 | ABA=0 | ROS CA | some 8-nitro-cGMP, 8-nitro-cGMP | 8-nitro-cGMP | Some/ON | 23396828 |
| 456 | ABA=0 | ROS CA | some actin reorganization | Actin Reorganization | Some | 18088331 |
| 457 | ABA=0 | CalM CA | some SLAC1, SLAC1 | SLAC1 | Some/ON | 19910530 |
| 458 | ABA=0 | MPK9/12 KO | no SLAC1 | SLAC1 | OFF | 19910530 |
| 459 | ABA=1 | MPK9/12 KO | no SLAC1 | SLAC1 | OFF | 19910530 |
| 460 | ABA=0 | CalM CA, MPK9/12 KO | no SLAC1 | SLAC1 | OFF | 19910530 |
| 461 | ABA=0 | PA CA | no $Ca^{2+}_c$ | $Ca^{2+}_c$ | OFF | 10518598 |
| 462 | ABA=0 | GHR1 KO | no ROS | ROS | OFF | 22730405 |
| 463 | ABA=1 | GHR1 KO | ROS | ROS | ON | 22730405 |
| 464 | ABA=0 | ABI1 CA | no NO | NO | OFF | 12446847, 19690149 |
| 465 | ABA=1 | ABI1 CA | NO | NO | ON | 12446847, 19690149 |
| 466 | ABA=0 | ABI2 CA | no NO | NO | OFF | 12446847 |
| 467 | ABA=1 | ABI2 CA | NO | NO | ON | 12446847 |
| 468 | ABA=0 | PA CA | RBOH | RBOH | ON | 19690149 |
| 469 | ABA=0 | PA CA | ROS<br>some ROS (less than ABA=1) | ROS | Some/ON | 19690149, 22392080 |
| 470 | ABA=0 | PA CA, PLDα KO, PLDδ KO | some ROS (less than ABA=1) | ROS | Some | 22392080 |
| 471 | ABA=0 | PA CA | no NO | NO | OFF | 22392080 |
| 472 | ABA=0 | PA CA, PLDα KO, PLDδ KO | no NO | NO | OFF | 22392080 |
| 473 | ABA=0 | PA CA | $pH_c$ | $pH_c$ | ON | 22392080 |
| 474 | ABA=0 | PA CA, PLDα KO, PLDδ KO | $pH_c$ | $pH_c$ | Some/ON | 22392080 |
| 475 | ABA=0 | PC CA | no RBOH* | RBOH | OFF | 19690149 |
| 476 | ABA=0 | PC CA | no ROS* | ROS | OFF | 19690149 |
| 477 | ABA=0 | PA CA, RBOH KO | no ROS | ROS | OFF | 19690149 |
| 478 | ABA=0 | WT | no OST1 | OST1 | OFF | 26192964 |
| 479 | ABA=1 | WT | OST1 | OST1 | ON | 26192964 |
| 480 | ABA=0 | ABI1 KO, ABI2 KO, HAB1 KO, PP2CA KO | no OST1 | OST1 | OFF | 26192964 |
| 481 | ABA=1 | ABI1 KO, ABI2 KO, HAB1 KO, PP2CA KO | OST1 | OST1 | ON | 26192964 |

| | | | | | | |
|---|---|---|---|---|---|---|
| 482 | ABA=0 | CPK6 KO, CPK23 KO | no OST1 | OST1 | OFF | 26192964 |
| 483 | ABA=1 | CPK6 KO, CPK23 KO | OST1 | OST1 | ON | 26192964 |
| 484 | ABA=0 | CPK6 KO | no OST1 | OST1 | OFF | 26192964 |
| 485 | ABA=1 | CPK6 KO | OST1 | OST1 | ON | 26192964 |
| 486 | ABA=0 | WT | ABI1 | ABI1 | ON | 19407143 |
| 487 | ABA=1 | WT | no ABI1 | ABI1 | OFF | 19407143 |
| 488 | ABA=0 | WT | ABI2 | ABI2 | ON | 19407143 |
| 489 | ABA=1 | WT | no ABI2 | ABI2 | OFF | 19407143 |
| 490 | ABA=0 | WT | HAB1 | HAB1 | ON | 19874541 |
| 491 | ABA=1 | WT | no HAB1 | HAB1 | OFF | 19407142 |
| 492 | ABA=0 | WT | PP2CA | PP2CA | ON | 22198272 |
| 493 | ABA=1 | WT | no PP2CA | PP2CA | OFF | 22198272 |
| 494 | ABA=0 | WT | $H^+$ ATPase | $H^+$ ATPase | ON | 15563626, 23946352 |
| 495 | ABA=1 | WT | no $H^+$ ATPase | $H^+$ ATPase | OFF | 15563626, 23946352 |
| 496 | ABA=0 | WT | Malate | Malate | ON | 12392716 |
| 497 | ABA=1 | WT | no Malate | Malate | OFF | 24419583 |
| 498 | ABA=0 | WT | no InsP3 | InsP3 | OFF | 12226236 |
| 499 | ABA=1 | WT | InsP3 | InsP3 | ON | 12226236 |
| 500 | ABA=0 | WT | no S1P/PhytoS1P | S1P/PhytoS1P | OFF | 12789341 |
| 501 | ABA=1 | WT | S1P/PhytoS1P | S1P/PhytoS1P | ON | 12789341 |
| 502 | ABA=0 | WT | no SPHK1/2 | SPHK1/2 | OFF | 12789341 |
| 503 | ABA=1 | WT | SPHK1/2 | SPHK1/2 | ON | 12789341 |
| 504 | ABA=0 | WT | no RCARs | RCARs | OFF | 19874541 |
| 505 | ABA=1 | WT | RCARs | RCARs | ON | 19874541 |
